# Normative modeling of brain morphometry in Clinical High-Risk for Psychosis

**DOI:** 10.1101/2023.01.17.523348

**Authors:** Shalaila S Haas, Ruiyang Ge, Ingrid Agartz, G. Paul Amminger, Ole A Andreassen, Peter Bachman, Inmaculada Baeza, Sunah Choi, Tiziano Colibazzi, Vanessa L Cropley, Camilo de la Fuente-Sandoval, Bjørn H Ebdrup, Adriana Fortea, Paolo Fusar-Poli, Birte Yding Glenthøj, Louise Birkedal Glenthøj, Kristen M Haut, Rebecca A Hayes, Karsten Heekeren, Christine I Hooker, Wu Jeong Hwang, Neda Jahanshad, Michael Kaess, Kiyoto Kasai, Naoyuki Katagiri, Minah Kim, Jochen Kindler, Shinsuke Koike, Tina D Kristensen, Jun Soo Kwon, Stephen M Lawrie, Jimmy Lee, Imke LJ Lemmers-Jansen, Ashleigh Lin, Xiaoqian Ma, Daniel H Mathalon, Philip McGuire, Chantal Michel, Romina Mizrahi, Masafumi Mizuno, Paul Møller, Ricardo Mora-Durán, Barnaby Nelson, Takahiro Nemoto, Merete Nordentoft, Dorte Nordholm, Maria A Omelchenko, Christos Pantelis, Jose C Pariente, Jayachandra M Raghava, Francisco Reyes-Madrigal, Jan I Røssberg, Wulf Rössler, Dean F Salisbury, Daiki Sasabayashi, Ulrich Schall, Lukasz Smigielski, Gisela Sugranyes, Michio Suzuki, Tsutomu Takahashi, Christian K Tamnes, Anastasia Theodoridou, Sophia I Thomopoulos, Paul M Thompson, Alexander S Tomyshev, Peter J Uhlhaas, Tor G Værnes, Therese AMJ van Amelsvoort, Theo GM van Erp, James A Waltz, Christina Wenneberg, Lars T Westlye, Stephen J Wood, Juan H Zhou, Dennis Hernaus, Maria Jalbrzikowski, René S Kahn, Cheryl M Corcoran, Sophia Frangou, the ENIGMA Clinical High Risk for Psychosis Working Group

## Abstract

**Importance:** The lack of robust neuroanatomical markers of psychosis risk has been traditionally attributed to heterogeneity. A complementary hypothesis is that variation in neuroanatomical measures in the majority of individuals at psychosis risk may be nested within the range observed in healthy individuals.

**Objective:** To quantify deviations from the normative range of neuroanatomical variation in individuals at clinical high-risk for psychosis (CHR-P) and evaluate their overlap with healthy variation and their association with positive symptoms, cognition, and conversion to a psychotic disorder.

**Design, Setting, and Participants:** Clinical, IQ and FreeSurfer-derived regional measures of cortical thickness (CT), cortical surface area (SA), and subcortical volume (SV) from 1,340 CHR-P individuals [47.09% female; mean age: 20.75 (4.74) years] and 1,237 healthy individuals [44.70% female; mean age: 22.32 (4.95) years] from 29 international sites participating in the ENIGMA Clinical High Risk for Psychosis Working Group.

**Main Outcomes and Measures:** For each regional morphometric measure, z-scores were computed that index the degree of deviation from the normative means of that measure in a healthy reference population (N=37,407). Average deviation scores (ADS) for CT, SA, SV, and globally across all measures (G) were generated by averaging the respective regional z-scores. Regression analyses were used to quantify the association of deviation scores with clinical severity and cognition and two-proportion z-tests to identify case-control differences in the proportion of individuals with infranormal (z<-1.96) or supranormal (z>1.96) scores.

**Results:** CHR-P and healthy individuals overlapped in the distributions of the observed values, regional z-scores, and all ADS vales. The proportion of CHR-P individuals with infranormal or supranormal values in any metric was low (<12%) and similar to that of healthy individuals. CHR-P individuals who converted to psychosis compared to those who did not convert had a higher percentage of infranormal values in temporal regions (5-7% vs 0.9-1.4%). In the CHR-P group, only the ADS_SA_ showed significant but weak associations (|β|<0.09; P_FDR_<0.05) with positive symptoms and IQ.

**Conclusions and Relevance:** The study findings challenge the usefulness of macroscale neuromorphometric measures as diagnostic biomarkers of psychosis risk and suggest that such measures do not provide an adequate explanation for psychosis risk.

**Key points:** *Question:* Is the risk of psychosis associated with brain morphometric changes that deviate significantly from healthy variation?

*Findings:* In this study of 1340 individuals high-risk for psychosis (CHR-P) and 1237 healthy participants, individual-level variation in macroscale neuromorphometric measures of the CHR-P group was largely nested within healthy variation and was not associated with the severity of positive psychotic symptoms or conversion to a psychotic disorder.

*Meaning:* The findings suggest the macroscale neuromorphometric measures have limited utility as diagnostic biomarkers of psychosis risk.

## Introduction

Schizophrenia is a mental disorder characterized by psychotic and cognitive symptoms^1^ and significant psychosocial disability.^2^ Similar abnormalities are also present in individuals at clinical high-risk for psychosis (CHR-P) who typically experience attenuated or brief psychotic symptoms^3^ and cognitive difficulties, and an elevated risk of developing psychosis at rate of 20% at 2 years and 35% at 10 years.^5^ A better understanding of the neurobiology of CHR states holds the promise of improving early detection and preventive strategies.^6^

Multiple magnetic resonance imaging (MRI) studies have focused on identifying neuroanatomical alterations in CHR-P compared to healthy individuals (HI). Two meta-analyses of these studies have highlighted cortical thickness (CT) reductions of small effect size in the frontotemporal regions of CHR individuals^7,8^ while a mega-analysis of brain morphometric data from 1792 CHR-P and 1377 HI from the CHR-P Working Group of the Enhancing Neuroimaging Genetics through Meta-analysis (ENIGMA) Consortium found that such CT reductions were widespread (Cohen d range of -0.17 to -0.09).^9^

Recently psychiatric neuroimaging has turned to normative modeling, which quantifies individual-level deviation in brain-derived phenotypes relative to a normative reference population.^10^ The advantage of this approach is that it can test whether psychiatric disorders are associated with substantial deviation from healthy variation in measures of brain organization. Normative modeling has yet to be applied to CHR-P states, but there are two studies on patients with established schizophrenia that are of direct relevance.^11,12^ In both studies, brain morphometry measures with values below the 5th or above the 95th percentile of the normative range were respectively considered infranormal and supranormal. Lv et al. (2020)^11^ calculated normative models of CT from 195 HI and applied them to 322 individuals with schizophrenia; 10-15% of patients showed infranormal CT values in temporal and ventromedial frontal regions and 3% of patients had supranormal values mainly in the paracentral lobule. Wolfers and colleagues (2021)^12^ developed normative models from voxel-based morphometry data from three samples of HI (N1=400, N2=312, N3=256) and applied them to data from corresponding samples of patients with schizophrenia (N1=94, N2=105, N3=163); only a low percentage of voxels (<2%) had extreme values in patients across samples; voxels with infranormal values were mostly located within temporal, medial frontal, and posterior cingulate regions.

It is currently unknown whether regional deviations from healthy neuroanatomical variation in brain morphometry might be present in CHR-P individuals and whether they explain substantial variance in positive symptoms or cognition. At least one study has suggested that normative deviation is better than raw volumes in predicting psychotic symptoms.^13^ Addressing these questions is important for two reasons. First, vulnerabilities during brain development, as inferred from the presence of deviations from normative neuroanatomical trajectories, may set the scene for the brain changes observed in established cases of schizophrenia. Second, deviation from healthy variation in brain neuroanatomy may prove informative in identifying those CHR-P individuals that convert or experience more severe clinical presentations. To test these hypotheses, the current study derived age- and sex-specific normative models of regional morphometry from an independent dataset of HI and applied them to the ENIGMA CHR-P Working Group sample which represents the largest available dataset of individual-level morphometric measures from CHR-P individuals.^9^

## Methods

### Study Sample

The study sample was derived from the pooled dataset of CHR-P and HI held by the ENIGMA CHR-P Working Group (eMethods and eTable 1). At each site, CHR-P status was ascertained using either the Structured Interview for Prodromal Syndromes (SIPS) or the Comprehensive Assessment of At-Risk Mental States (CAARMS) (eMethods and eTable 2). Additional site-specific eligibility criteria are shown in eTable 1. At each site, whole-brain T1-weighted MRI data obtained from each participant (eMethods and eTable 3) were parcellated and segmented using standard FreeSurfer pipelines (https://surfer.nmr.mgh.harvard.edu/) to yield estimates of total intracranial volume (ICV), regional measures of CT (N=68), surface area (SA) (N=68), and subcortical volume (SV) (N=14) (eTable 4). These measures were then assessed using the ENIGMA consortium quality assessment pipeline.^14-17^ Ethical approval for data collection and sharing was obtained by the Institutional Review Board at each site. Participant data were shared after all identifying information was removed.

**Table 1.** Characteristics of the Sample at Clinical High-Risk for Psychosis (CHR-P)

| **All CHR-P individuals** |  |
| --- | --- |
| Age, years, mean (SD) | 20.75 (4.74) |
| Female Sex, number (%) | 631 (47.09%) |
| SIPS Positive Symptoms Score, mean (SD)<sup>a</sup> | 10.93 (4.66) |
| CAARMS Positive Symptoms Scores, mean (SD)<sup>b</sup> | 10.37 (4.03) |
| IQ, mean z-score (SD)<sup>c</sup> | -0.21 (1.00) |
| Prescribed Antipsychotic Medication, number (%)<sup>d</sup> | 243 (18.63%) |
| Months of follow-up, mean (SD)<sup>e</sup> | 19.71 (13.97) |
| Converters, N (%)<sup>e</sup> | 157 (14.31%) |
| **CHR-P individuals that converted to a psychotic disorder** |  |
| Age, years, mean (SD) | 20.09 (4.68) |
| Female Sex, number (%) | 64 (40.76%) |
| SIPS Positive Symptoms Score, mean (SD)<sup>f</sup> | 12.12 (5.06) |
| CAARMS Positive Symptoms Scores, mean (SD)<sup>f</sup> | 10.71 (4.24) |
| IQ, mean z-score (SD)<sup>g</sup> | -0.29 (1.03) |
| Prescribed Antipsychotic Medication, number (%)<sup>h</sup> | 32 (20.38%) |
| CAARMS = Comprehensive Assessment of At-Risk Mental States; SIPS = Structured Interview for Psychosis-risk Syndromes; SD = standard Deviation. Positive symptom ratings at the time of scanning were available for the entire sample (N = 1340), assessed either with SIPS or CAARMS; |  |
| <sup>a</sup> SIPS was used to assess CHR-P status in 806 CHR-P participants; <sup>b</sup> CAARMS was used to assess CHR-P status in 534 CHR-P participants; <sup>c</sup> Estimates of IQ were available in 924 CHR-P participants; z-scores were used to accommodate site differences in the instruments used (eTable 1); <sup>d</sup> Medication status at the time of scanning was available in 1304 CHR-P individuals; <sup>e</sup> Conversion status was known for 1097 CHR-P participants but information about the length of the follow-up period was available for 975 CHR-P individuals; <sup>f</sup> SIPS and CAARMS were respectively used to assess 115 and 42 individuals who converted; <sup>g</sup> IQ estimates were available in 109 CHR-P individuals that converted; <sup>h</sup> Medication status was available in 157 CHR-P individuals that converted |  |

The current study sample comprises participants who had both high-quality brain morphometric data and complete SIPS or CAARMS ratings at the time of their scan (eMethods and eFigure 1). Based on these criteria we included 1,340 CHR-P individuals [47.09% female; age range: 9.5 to 39 years; mean (SD) age: 20.75 (4.74) years] and 1,237 HI [44.70% female; age range: 12 to 39.87 years; mean (SD) age: 22.32 (4.95) years] (Table 1, eTables 5 and 6). Conversion status at a mean follow-up time of 19.71 (13.97) months was available for 1,097 CHR-P individuals (Table 1 and eTable 6). Individuals that converted to a psychotic disorder (CHR-PC) (n=157) had significantly higher positive symptoms at the time of scanning (mean z-score [SD] = 0.21 [1.08]) than those who did not convert (CHR-PNC) (N=940) (mean z-score [SD] = -0.05 [1.01]; T = 2.99; P = 0.003), but the two groups did not differ in age, sex, or IQ (all P > 0.07).

**Figure 1.**
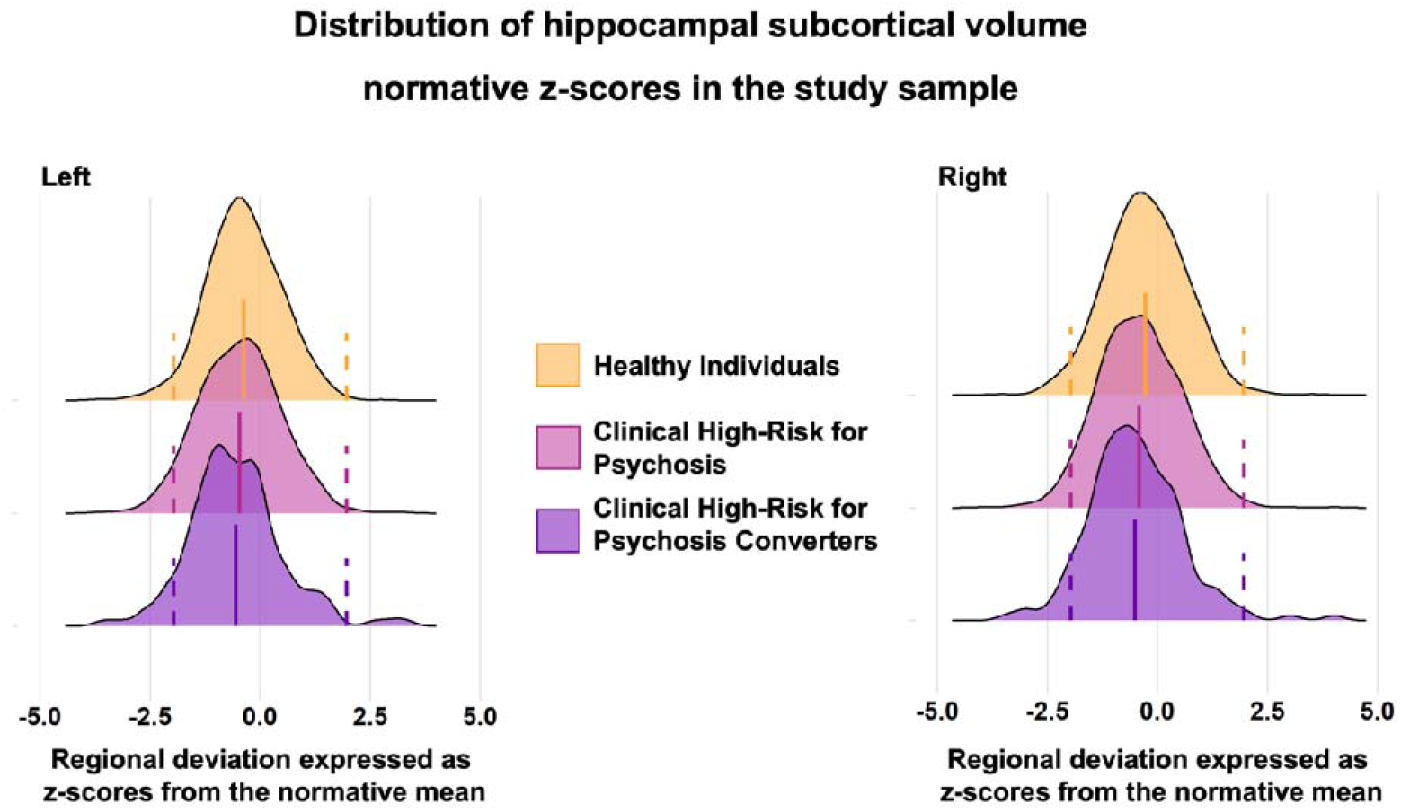
Distribution of hippocampal subcortical volume normative z-scores in the study sample. The figure presents the distribution of the left and right hippocampus regional normative z-scores in healthy individuals (HI), individuals at clinical high-risk for psychosis (CHR-P) and CHR-P that converted to full-blown psychosis (CHR-PC). The results for the remaining regional normative z-score are presented in the eVideo. The dotted lines represent the cutoffs for infranormal and supranormal values at z = |1.96|.

### Clinical Data

The ratings of CAARMS and SIPS converge only for positive symptoms (eMethods and eTable 2); these ratings were converted to z-scores to enable cross-site harmonization. Similarly, IQ estimates were converted to z-scores to accommodate the different instruments used across sites (eTable 1). Information was also available on medication exposure at the time of scanning.

### Normative models of brain morphometry

The normative models for each of the regional CT, SA, and SV measures (eTable 4) were generated using CentileBrain, an empirically validated framework for normative models of brain morphometry developed using data from an independent multi-site sample of 37,407 HI (53.3% female; aged 3-90 years) (https://centilebrain.org/).^18^ Details of the sample, procedures, model performance, and code are presented in the eMethods. Fractional polynomial regression was used to generate sex-specific models for each measure while accounting for site using ComBat-GAM harmonization.^19^ The ICV, mean CT, and mean SA were included in the models of the regional measures of SV, CT, and SA respectively.

### Computing deviation scores of regional morphometric measures

The CentileBrain model parameters were then applied to each regional CT, SA, and SV measure of the CHR-P and HI of the ENIGMA sample. For each measure in each participant, we estimated the degree of normative deviation from the reference population mean as a z-score computed by subtracting the predicted (Ŷ) from the raw value (Y_o_) of that measure and then dividing the difference by the root mean square error of the model (eFigure 2).^20,21^ A positive/negative z-score indicates that the value of the corresponding morphometric measure is higher/lower than the normative mean. As per previous literature,^15,16^ we defined regional z-scores as infranormal when below z=-1.96 or supranormal when above z=1.96, corresponding to the 5th and 95th percentile respectively. Intermediate values (i.e., between z=-1.96 and z=1.96) were designated as “within normal range”.

**Figure 2.**
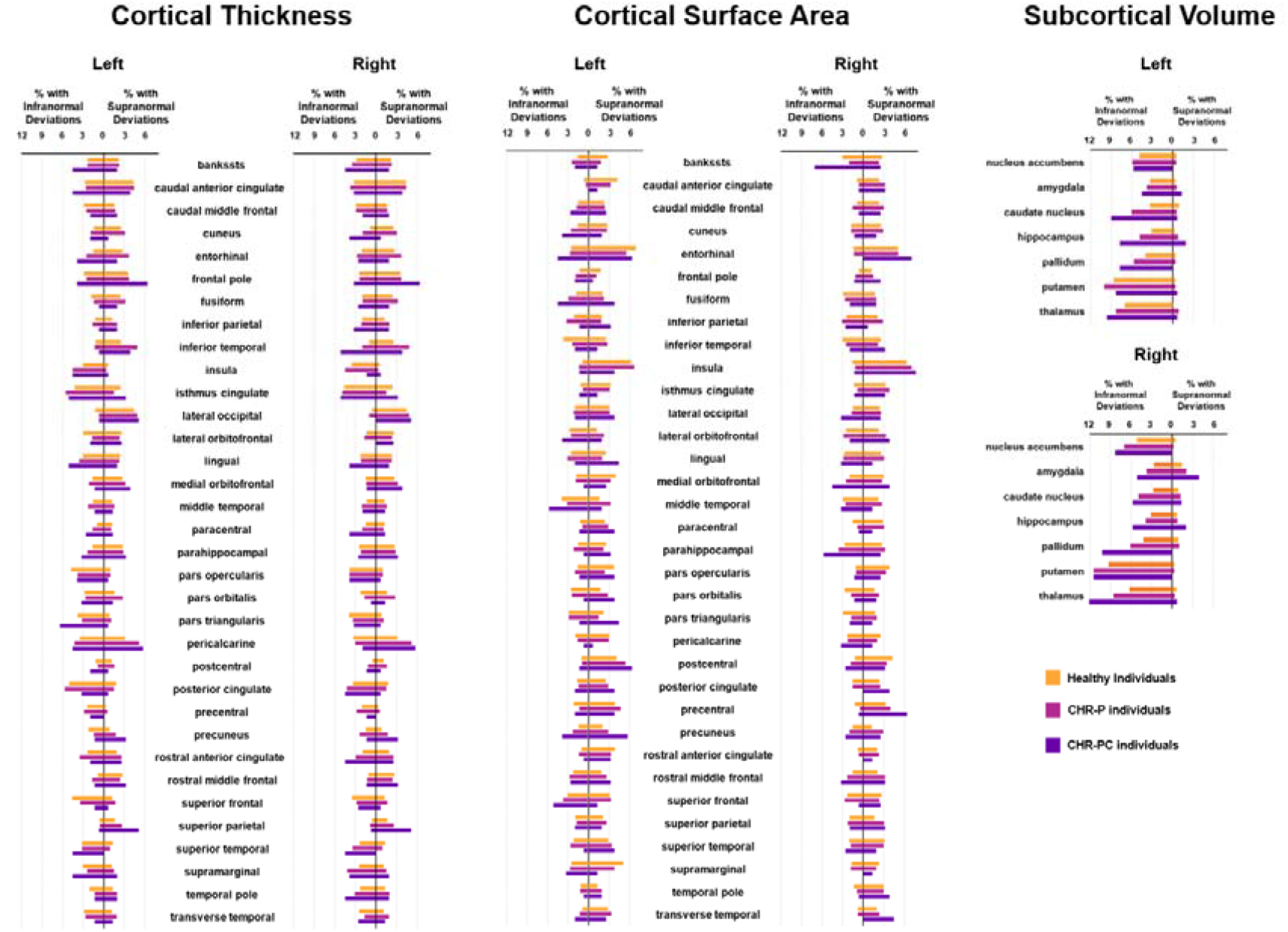
Percentage of subjects with infra- or supranormal regional normative z-scores. The proportion of healthy individuals, clinical high-risk for psychosis (CHR-P), and cinical high-risk for psychosis converters with infranormal (left) and supranormal deviations (right) are presented for each hemisphere for cortical thickness, cortical surface area, and subcortical volume.

### Computation of average deviation scores

We averaged the regional z-scores in each participant to generate an “average deviation score” for CT (ADS_CT_), SA (ADS_SA_), and SV (ADS_SV_). ADS were not weighted for the size of the region to enhance reproducibility. Positive or negative ADS values indicate a general pattern of deviations that are above or below the normative reference values. ADS scores were further averaged to generate a “global average deviation score” (ADS_G_). Using the same criteria as for the z-scores, each ADS was also designated as infranormal, supranormal, or within the normal range. In supplemental analyses, we also explored alternate definitions of ADS, by averaging positive and negative z-scores separately for each neuroimaging phenotype (eMethods).

### Statistical Analyses

Statistical significance across all tests performed was set at P_FDR_<0.05 as per the Benjamini-Hochberg false discovery rate (FDR) correction for multiple comparisons. The robustness of the results was confirmed using a leave-one-site-out approach.

The following analyses were conducted: (i) we calculated the percentage of CHR-P and HI from the ENIGMA sample that had supranormal or infra-normal z-scores in any regional measure and in any of ADS_CT_, ADS_SA_, ADS_SV,_ and ADS_G_ (eMethods); group differences in the proportion of individuals with supra-or infranormal z-scores were examined using the two-proportions z-test implemented in R version 4.1.2; (ii) within the CHR-P group, linear regression (implemented with the *“lm*” function in R version 4.1.2) was used to assess associations between positive symptoms and IQ with each regional z-score, the observed value of each regional morphometric measure, and each ADS; age was included as predictor in all aforementioned regression models due to its significant association with positive symptoms and IQ (P_FDR_<0.05), while sex was included only in the models with observed data as the z-scores and ADSs were derived from sex-specific models; (iii) we repeated the above analyses separately for CHR-PC and CHR-PNC individuals; and (iv) in HI, we conducted regression analyses to assess associations between IQ and the brain regional z-scores, observed values, and ADS.

Supplemental testing involved group comparisons of mean regional and ADS scores, the use of an alternate parcellation template, and repeating the aforementioned analyses in subsamples of CHR-P sub-syndromes (i.e., attenuated psychotic symptoms syndrome, brief intermittent psychotic symptoms syndrome, and genetic risk and functional deterioration syndrome); additional analyses focused on medication status and alternate ADS definitions (eMethods).

## Results

### Infra- and supranormal deviations in brain morphometry in CHR-P and HI

The distributions of the z-scores and observed values of all regional brain morphometric measures of CHR-P and HI showed complete overlap. The left and right hippocampus are used as exemplars in Figure 1 while the corresponding figures for all other regions are included in the eVideo. Moreover, the distribution overlap was independent of parcellation used to define subregions (eFigure 3 and 4).

**Figure 3.**
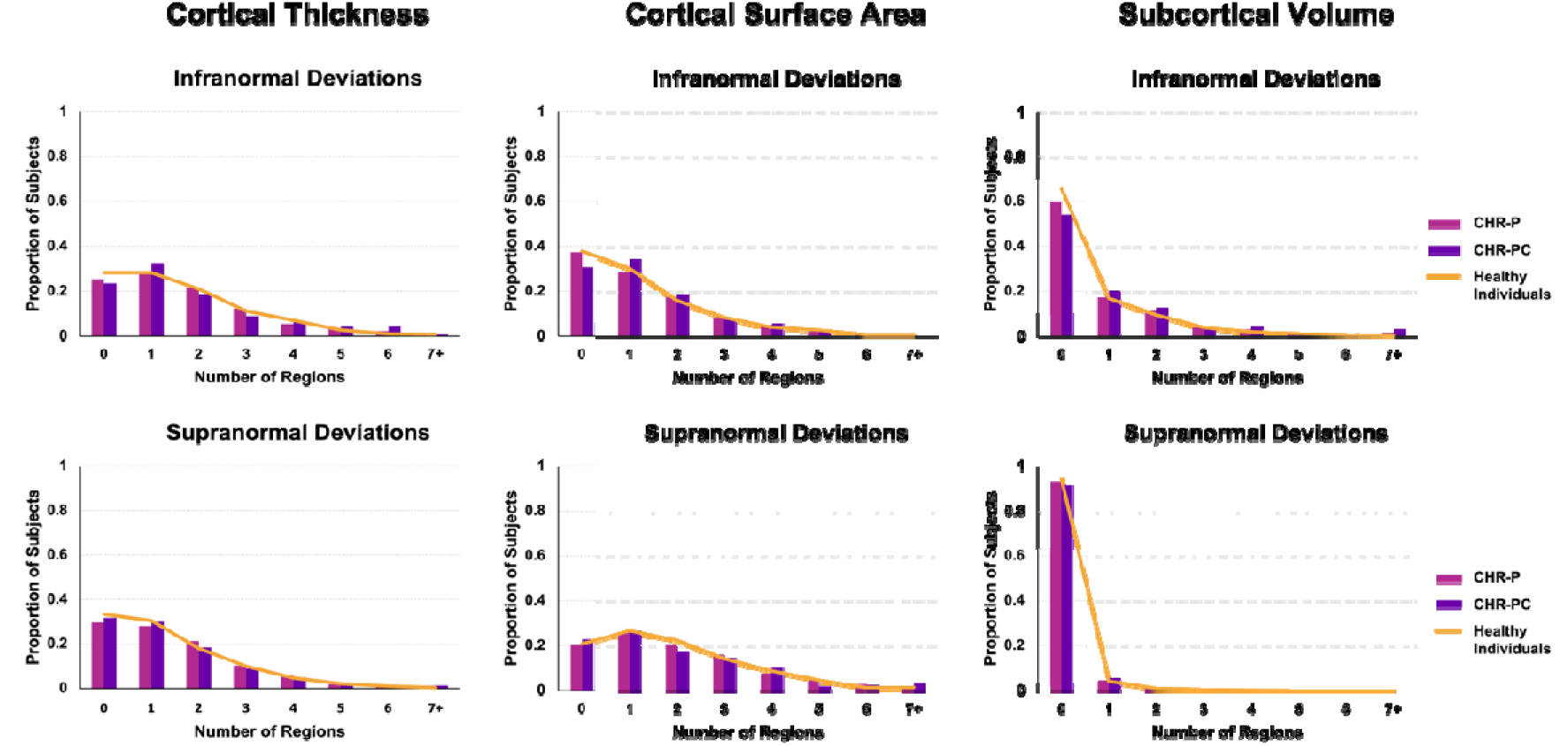
Distribution of the total number of regions with infra- or supranormal regional normative z-scores. Bar plots show the distribution of the total number of regions per individual with infra-normal (top row) and supra-normal (bottom row) deviations from the normative model for A) cortical thickness, B) cortical surface area, and C) subcortical volume separately in healthy individuals, clinical high-risk for psychosis (CHR-P) individuals, and clinical high-risk for psychosis converters (CHR-PC).

**Figure 4.**
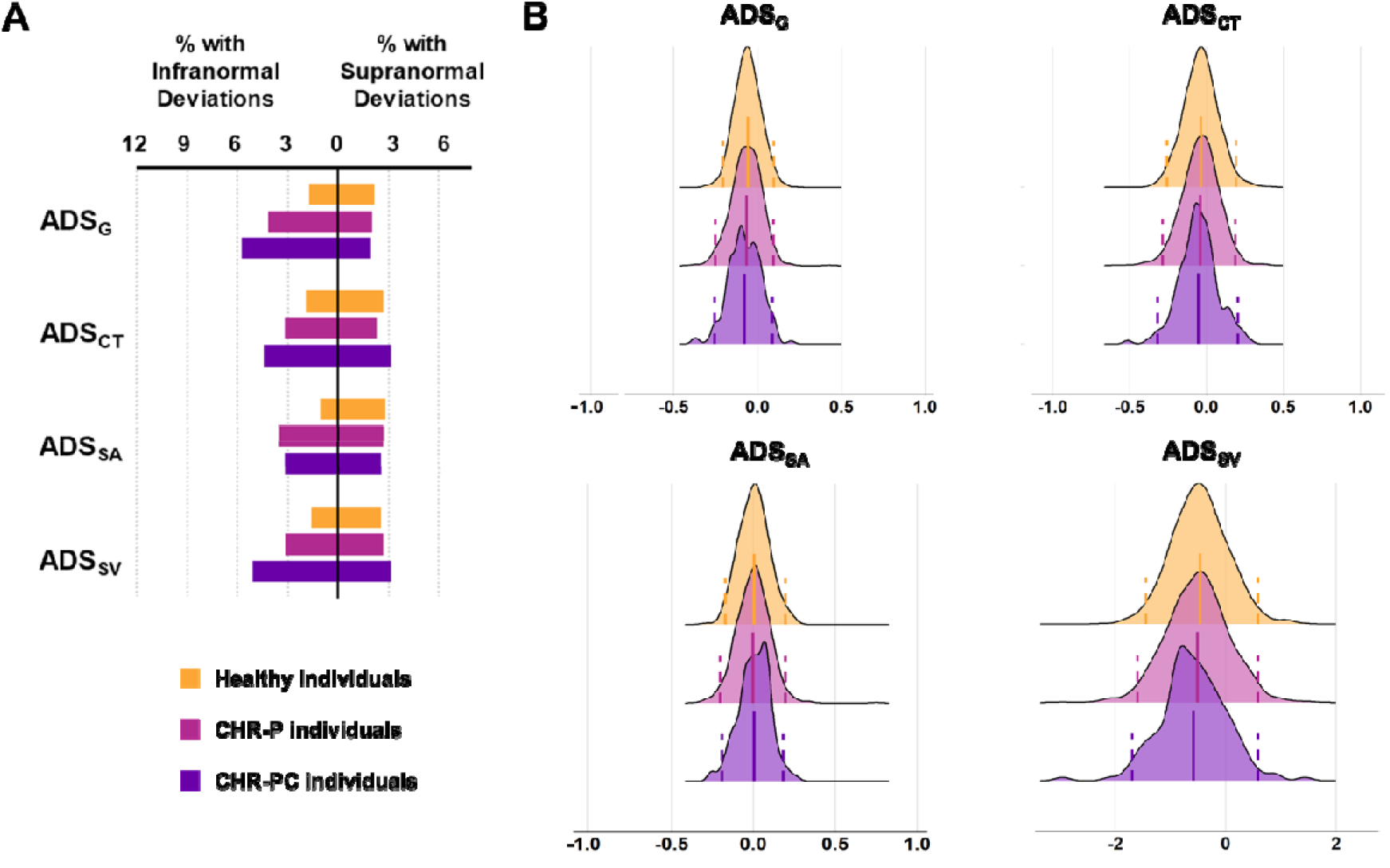
Distributions of average deviation scores and percentage of subjects with infra- or supranormal regional normative z-scores. A) Percentage of healthy, clinical high-risk for psychosis (CHR-P), and clinical high-risk for psychosis converters (CHR-PC) with supra-or infranormal global average deviation score (ADS_G_), the average deviation score for cortical thickness (ADS_CT_), average deviation score for cortical surface area (ADS_SA_), and average deviation score for subcortical volumes (ADS_SV_). B) The distributions of the average deviation scores in CHR-P (magenta color), CHR-PC (purple color), and healthy individuals (yellow color) for ADS_G_, ADS_CT_, ADS_SA_, and ADS_SV_. The dotted lines represent the cutoffs for infranormal and supranormal values at z = |1.96|.

The percentage of CHR-P and HI with supra-or infranormal z-scores in each morphometric measure are shown in Figure 2. Infranormal regional CT z-scores were noted in 0.52-5.67% of CHR-P individuals and 0.49-5.01% of HI; the corresponding range for supranormal z-scores were 0.37-5.15% and 0.32-5.50%. Infranormal regional SA z-scores were noted in 0.30-3.66% of CHR-P individuals and 0.65-3.88% of HI; the corresponding range for supranormal z-scores were 1.12-7.01% and 1.21-6.95%. Infranormal regional SV z-scores were noted in 3.73-11.42% of CHR-P individuals and 2.67-9.30% of HI; the corresponding range for supranormal z-scores were 0.07-2.01% and 0.08-1.37%. Infranormal z-scores in any regional CT, SA, and SV were observed in 74.63%, 62.76%, and 40.00% across CHR-P, respectively; the corresponding infranormal z-scores of HI were 71.71%, 62.25%, and 34.93% (Figure 3). Supranormal z-scores in any regional CT, SA, and SV were observed in 70.37%, 79.70%, and 6.64% across CHR-P, respectively; the corresponding supranormal z-scores in HI is 66.61%, 79.22%, and 5.58% (Figure 3). There were no significant group differences in the percentage of individuals with supra-or infranormal regional values (P_FDR_>0.05; eTable 7). Compared to unmedicated CHR-individuals, those medicated had a greater proportion with supranormal regional z-scores for the surface area of the left lateral occipital lobe (χ^2^ = 13.92, P_FDR_ = 0.03) but no other differences (eTable 8).

### Supra- and infranormal average deviation scores

The percentage of CHR-P and HI with supra-or infranormal ADS values are shown in Figure 4A. The distributions in both groups showed a near complete overlap (Figure 4B). Infranormal ADS_CT_, ADS_SA_, ADS_SV_, and ADS_G_ were respectively observed in 3.21%, 3.51%, 3.13%, and 4.18% of CHR-P individuals; the corresponding percentages in HI were 1.94%, 1.05%, 1.62%, and 1.70%. Supranormal ADS_CT_, ADS_SA_, ADS_SV_, and ADS_G_ were respectively observed in 2.31%, 2.69%, 2.69%, and 2.01% of CHR-P individuals; the corresponding percentages in HI were 2.67%, 2.83%, 2.59%, and 2.18%. A significantly higher percentage of CHR-P individuals had infranormal ADS_G_ (χ^2^ = 12.82, P_FDR_ = 6.85E-4), ADS_SV_ (χ^2^ = 5.68, P_FDR_ = 0.02), and ADS_SA_ (χ^2^ = 16.01, P_FDR_ = 2.53E-4) (eTable 9). There were no differences in the percentage of CHR-P individuals with infra-or supranormal ADS depending on their medication status (P_FDR_>0.05; eTable 10).

### Associations of regional z-scores and observed values with positive symptoms and IQ

Within the CHR-P group, positive associations were noted only between IQ and the z-scores of the left caudate volume (left: β= 0.11, P_FDR_ = 0.05), and surface area of the left cuneus (β= 0.11, P_FDR_ = 0.05) (eFigure 5A and B; eTable 11). When analyses were repeated using the observed regional morphometric values, there were no significant associations with IQ or positive symptoms (eTable 12). The same pattern of results for z-scores and observed values was observed when CHR-P individuals exposed to medication were excluded. In HI, no significant associations were noted either between IQ and z-scores or between IQ and the observed values (P_FDR_ > 0.05) (eTable 13).

### Associations of average deviation scores with positive symptoms and IQ

Within the CHR-P group, positive symptoms were negatively associated with ADS_SA_ (β = - 0.08, P_FDR_ = 0.02; eFigure 5C), while IQ was positively associated with ADS_SA_ (β = 0.09, P_FDR_ = 0.02) and ADS_G_ (β = 0.10, P_FDR_ = 0.01) (eFigure 5D and E; eTable 14). This pattern of associations was robust to medication status and leave-one-out analysis (eFigure 6). In HI, positive associations were also present between IQ and ADS_SA,_ and ADS_G_ (eTable 14).

### CHR-P individuals who converted to a psychotic disorder

The percentage of CHR-PC and CHR-PNC individuals with infranormal and supranormal regional z-scores and ADS are shown in Figure 3 and eTable 15). There was a significantly greater percentage of CHR-PC (5.10%) than HI (0.89%) with infranormal z-scores for the thickness of the right inferior temporal lobe (χ^2^ = 15.34, P_FDR_ = 0.01) and a significantly greater percentage of CHR-PC (7.01%) than CHR-PNC (1.38%) with infranormal z-scores for the surface area of the right banks of the superior temporal sulcus (χ^2^ = 17.34, P_FDR_ = 4.69E-3). No further differences were identified. As for the entire CHR-P sample, IQ was positively associated with ADS_SA_ (β = 0.26, P_FDR_ = 0.02) and ADS_G_ (β = 0.21, P_FDR_ = 0.05) in CHR-PC individuals. Because of the smaller sample size, associations between positive symptoms and ADS_SA_ were no longer significant within the CHR-PC group but retained the same direction (β = -0.12, P_FDR_ = 0.20). No other significant associations were found between regional z-scores or ADS and IQ or positive symptoms in the CHR-PC or CHR-PNC subsamples (eTable 16).

### Supplemental Analyses

Group differences in regional z-scores and ADS between HI and the entire CHR-P or the CHR-PC group were of small effect sizes (Cohen’s |d| <0.26) (eTable 17). Similarly, the effect size differences in the above metrics in CHR-P exposed or not exposed to antipsychotics were also negligible (Cohen’s |d| <0.24) (eTable 17). Analyses of CHR syndromes (eResults) and alternate ADS did not provide additional insights (eTable 18).

## Discussion

This study found that variation in regional neuromorphometric measures in CHR-P individuals was nested within the healthy distribution while extreme deviations were present in a minority of CHR-P individuals and at proportions similar to those observed in healthy individuals. However, a greater proportion of CHR-P individuals had infranormal ADS_CT_, ADS_SA_, and ASD_SV_ values. Additionally, a higher percentage of CHR-PC individuals had infranormal values in temporal regions but none of the regional z-scores had meaningful associations with the severity of positive symptoms.

Prior case-control studies, including a study by Jalbrzikowski and colleagues who also used the ENIGMA CHR-P Working Group dataset, have reported subtle decrements in regional brain morphometry in CHR-P individuals.^7-9^ These findings are aligned with the observation of a higher proportion of CHR-P individuals had infranormal values for ADS. In the same dataset, Baldwin and colleagues,^22^ showed that individual-level heterogeneity was similar in CHR-P and healthy individuals and was not predictive of increased clinical severity. The current study extends our understanding of the role of brain morphometry for psychosis by showing that regional neuroanatomical variation in CHR-P individuals is nested within normative variation. A small minority of CHR-PC patients had pronounced decrements in the cortical thickness and surface area of temporal regions reinforcing the relevance of these regions for psychosis risk ^7-9^ and syndromal schizophrenia. ^11,12,23^

Regional deviation from normative patterns in the CHR-P individuals did not show meaningful associations with the severity of positive symptoms. The only exception was that higher ADS_SA_, indicating an overall pattern of positive regional deviations in cortical surface area, was associated with less severe positive symptoms in the CHR-P group (regardless of conversion status). The ADS_SA_ was also positively associated with IQ both in CHR-P and healthy individuals. The strength of these associations was low (|β|<0.20). Nevertheless, these findings resonate with prior reports of higher IQ being associated with greater cortical surface area expansion^24,25^ and may reflect the integrity of cellular processes relating to neurite remodelling and intra-cortical myelination that play a key role in cortical surface area expansion during early adulthood.^26, 27^

## Limitations

The study includes the largest neuroimaging dataset of CHR-P individuals and robust normative models derived from an independent reference sample. As is common with large-scale studies, the data were collected at multiple sites using different scanners and protocols. Although we accounted for site effects using MRI data harmonization and tested the robustness of the results using leave-one-site-out analyses, residual effects cannot be fully excluded but are unlikely to have influenced the overall pattern of the results. The neuroimaging data of the CHR individuals are cross-sectional and do not capture potential longitudinal changes that may be more informative^28^.

## Conclusions

In this study, regional variation in the neuroanatomy of CHR-P individuals was nested within the normal variation. The degree of neuroanatomical normative deviation showed minimal associations with positive symptoms and conversion status. These findings question the usefulness of neuromorphometry as a diagnostic biomarker of CHR-states.

## Acknowledgments

This study presents independent research funded by multiple agencies. The funding sources had no role in the study design, data collection, analysis, and interpretation of the data. The views expressed in the manuscript are those of the authors and do not necessarily represent those of any of the funding agencies. SSH is supported by NIH National Institute of Mental Health, grant T32MH122394. VLC has received investigator grant 1177370 and project grant 1065742 from the National Health and Medical Research Council. CdlF-S was supported by grants 261895 and 320662 from the Consejo Nacional de Ciencia y Tecnología (CONACyT), grants from CONACyT’s Sistema Nacional de Investigadores (SNI), and grant R21 MH117434 from the National Institutes of Health. LBG was supported by the TrygFoundation; the Danish Research Council on Independent Research; the Lundbeck Foundation Center for Clinical Intervention and Neuropsychiatric Schizophrenia Research, CINS. KMH received support from NIMH grant R01MH105246. RAH has received grants from the University of Pittsburgh Medical Center. CIH was supported by grant R01 MH105246 from the National Institutes of Health. WJH was supported by a grant of the Korea Health Technology R&D Project through the Korea Health Industry Development Institute (KHIDI) funded by the Ministry of Health & Welfare, Republic of Korea (grant number: HI19C1080). KK and SK received supports from Japan Agency for Medical Research and Development (AMED) under Grant Number JP18dm0307001, JP18dm0307004, and JP19dm0207069, and JST Moonshot R&D (JPMJMS2021). MKi has received grant 2019R1C1C1002457 from the National Research Foundation of Korea and grant 21-BR-03-01 from the KBRI basic research program through Korea Brain Research Institute, funded by the Ministry of Science & ITC. TK was supported by a Brain and Behavior Research Foundation 2021 NARSAD Young Investigator Grant (ID 30112). JSK has received grant 2020M3E5D9079910 from the National Research Foundation of Korea and grant 21-BR-03-01 from the KBRI basic research program through Korea Brain Research Institute, funded by the Ministry of Science & ITC. JL received grant NMRC/TCR/003/2008 from the National Medical Research Council Translational and Clinical Research Flagship Programme. AL is supported by a National Health and Medical Research Council (NHMRC) Emerging Leadership Fellowship 2010063. RM has received grants R01MH100043 and R01MH113564 from the National Institute of Mental Health and grants from the Brain and Behavior Research Foundation and Canadian Institutes of Health Research. BN is supported by a National Health and Medical Research Council (NHMRC) Senior Research Fellowship 1137687. TN has received grants from Otsuka as well as personal fees from Astellas, Eisai, Janssen Pharmaceuticals, Meiji Seika Pharma, Sumitomo Dainippon Pharma, and Takeda. MN is supported by Mental Health Center Copenhagen, Mental Health Services in the Capital Region of Denmark, and University of Copenhagen.DN has received grants R25-A2701 and R287-2018-1485 from the Lundbeck Foundation and grants from the Mental Health Services Capital Region of Denmark. PL-O and FR-M were supported by grants from SNI. WR has received grants from the Zurich Program for Sustainable Development of Mental Health Services. DFS was supported by the US National Institutes of Health, R01MH113533. DS has received grant JP18K15509 and JP22K15745 from the JSPS KAKENHI. US has received grant 569259 from the National Health and Medical Research Council of Australia. GS is supported by the Fundació Clínic Recerca Biomèdica, the Brain and Behavior Research Foundation (NARSAD Young Investigator Award 2017, grant ID: 26731), the Alicia Koplowitz Foundation and the Spanish Ministry of Health, Instituto de Salud Carlos III “Health Research Fund” (PI15/0444; PI18/0242; PI18/00976; PI2100330). MS has received grant JP20H03598 from the Japan Society for the Promotion of Science KAKENHI and grant JP19dk0307069s0203 from the Japan Agency for Medical Research and Development. TT was supported by JSPS KAKENHI Grant Number JP18K07550. CKT is supported by the Research Council of Norway (#223273, #288083, #323951) and the South-Eastern Norway Regional Health Authority (#2019069, #2021070, #2023012, #500189). PMT and SIT are supported in part by NIMH grants R01MH123163, R01MH121246, and R01MH116147. PJU has received grant MR/L011689/1 from the UK Medical Research Council. JAW has received grant 5R01MH115031 from the National Institute of Mental Health. SJW was funded by several grants from the National Health & Medical Research Council of Australia. Core funding for ENIGMA was provided by the NIH Big Data to Knowledge (BD2K) program under consortium grant U54 EB020403 to PMT. This work was supported in part through the computational resources and staff expertise provided by Scientific Computing at the Icahn School of Medicine at Mount Sinai.

