## Supplementary Material for "Normative modeling of brain morphometry in Clinical High-Risk for Psychosis"


**eMethods.**
**eResults.**
**eTable 1.** Information on the samples contributed by site to the Clinical High-Risk for Psychosis Working Group of the ENIGMA Consortium
**eTable 2.** SIPS and the CAARMS items and Clinical High-Risk criteria
**eTable 3.** Neuroimaging acquisition protocols and FreeSurfer version per site
**eTable 4.** FreeSurfer-derived morphometric measures
**eTable 5.** Demographic characteristics of study participants per site
**eTable 6.** Characteristics of Clinical High-Risk individuals defined with either the SIPS or CAARMS criteria
**eTable 7.** Percentage of individuals with infra- or supranormal normative regional z-scores based on group
**eTable 8.** Percentage of CHR-P with infra- or supranormal normative regional z-scores according to medication status
**eTable 9.** Percentage of individuals with infra- or supranormal normative average deviation scores based on group
**eTable 10.** Percentage of CHR-P with infra- or supranormal normative average deviation scores according to medication status
**eTable 11.** Associations between regional normative z-scores and positive symptoms and IQ in CHR-P
**eTable 12.** Associations between observed brain morphometric measures with positive symptoms and IQ in CHR-P
**eTable 13.** Associations between either regional z-scores or observed brain morphometric measures with IQ in healthy individuals
**eTable 14.** Associations between average deviation scores with positive symptoms and IQ in CHR-P and healthy individuals
**eTable 15.** Percentage of CHR-P with infra- or supranormal normative regional z-score according to conversion status.
**eTable 16.** Associations between regional z-scores and average deviation scores with positive symptoms and IQ based on conversion status
**eTable 17.** Effect size (Cohen's d) of group differences
**eTable 18.** Association between positive or negative average deviation scores with positive symptoms and IQ
**eFigure 1.** Flow diagram for study sample selection of Clinical High-Risk individuals and healthy individuals
**eFigure 2.** Illustrative Representation of Normative Modeling
**eFigure 3.** Distribution of normative regional z-scores for cortical thickness based on the Schaefer 400 parcellation
**eFigure 4.** Distribution of normative regional z-scores for cortical surface area based on the Schaefer 400 parcellation
**eFigure 5.** Associations between regional and average deviation scores and clinical measures in CHR-P.
**eFigure 6.** Associations between average deviation scores with the positive symptoms and IQ based on medication exposure and removing one site at a time.
**eReferences.**


This supplementary material has been provided by the authors to give readers additional information about their work.

**eMethods.**

**Clinical High-Risk for Psychosis Working Group of the ENIGMA Consortium**

The working group has pooled together clinical, neuroimaging, and cognitive data from multiple research sites around the world. Site-specific eligibility criteria and site-specific assessments used to determine clinical high-risk status for psychosis (CHR-P) are described in eTables 1 and 2.

**Determination of Clinical High-Risk status for Psychosis**

Clinical high-risk (CHR-P) status for Psychosis in each recruitment site was determined using either the Comprehensive Assessment of At-Risk Mental States (CAARMS)[1] or the Structured Interview for Prodromal Syndromes/Psychosis-Risk Syndromes (SIPS)[2,3]. Both are semi-structured interviews conducted by trained clinicians. The items of the instruments and the criteria required to meet CHR status are described in eTable 2.

The CAARMS consists of seven subscales: positive symptoms, cognitive change, emotional disturbances, negative symptoms, behavioral change, motor change, and general psychopathology. The positive symptoms subscale and an assessment of social and occupational functioning are used to determine CHR-P eligibility and syndrome subgroup. The positive symptoms subscale consists of four positive symptoms. Each symptom is rated on a scale from 0 (absent) to 6 (endorsed at a fully psychotic level). Based on different criteria (eTable 2) a CHR-P participant could be classified as having either attenuated psychotic symptoms (APS) or brief limited intermittent psychotic symptoms (BIPS) or a “trait/vulnerability” syndrome.

The SIPS has four subscales: positive, negative, disorganization, and general symptoms. The positive symptoms subscale was used to determine CHR-P eligibility and consists of five positive symptoms. Each symptom is rated on a scale from 0-6 as for the CAARMS. Based on different criteria (eTable 2) a participant could be classified as having APS, BIPS or a “genetic risk and deterioration” (GRD) syndrome (equivalent to the trait/vulnerability syndrome).

**Structural Neuroimaging**

At each recruitment site, participants underwent structural magnetic resonance imaging (sMRI) to obtain T1-weighted scans. Details of the acquisition protocols are provided in eTable 3. Images were parcellated and segmented at each respective site using standard pipelines implemented using FreeSurfer analysis software (https://surfer.nmr.mgh.harvard.edu/) to yield regional measures of cortical thickness (CT), cortical surface area (SA), and subcortical volume (SV) (eTable 4). Information on the FreeSurfer version used per site is shown in eTable 3. Quality assessment of the FreeSurfer output used standardized ENIGMA procedures (http://enigma.ini.usc.edu/protocols/imaging-protocols/).

**Study Sample**

For the current study sample, both CHR and healthy individuals were selected from the total sample available based on the availability of high-quality imaging data (<5% of missing regional measures or ICV values per individual scan) and completed CAARMS or SIPS ratings. eFigure 1 shows the selection flow diagram of the current study sample.

**Study Sample Characteristics**

eTable 5 presents the demographic breakdown of the study sample by site. Of the 1340 CHR individuals, 806 (60.15%) were assessed using SIPS and 534 (39.85%) using CAARMS. Their clinical characteristics are presented in eTable 6.

**Normative modeling**

We used the CentileBrain framework (https://centilebrain.org/) to generate sex-specific normative models for each of the FreeSurfer-extracted regional measure of cortical thickness (CT), Surface Area (SA), and Subcortical Volume (SV) (eTable 4) estimated using Multivariate Fractional Polynomial Regression (MFPR)[4] in a sample of 37,407 healthy individuals (53.3% female) with an age range of 3-90 years pooled from 81 datasets. Normative models for each morphometric measures were estimated using the following procedures: (i) data preparation: Sex-specific subsamples of each pooled sample were randomly split into a training subset (80%) and a test subset (20%) stratified by scanning site. In each subset, data were mean-centered after extreme values in each site-dataset were identified and removed using the interquartile range (IQR) method; (ii) site harmonization was implemented using ComBat-GAM[5]; (iii) normative model of each neuroimaging measure were generated using MFPR implemented using the “mfp” package in R and the closed test procedure (known as RA2) to select the most appropriate fractional polynomial. Initially, models were trained using 5-fold crossvalidation (5F-CV) in the corresponding sex-specific training subset with age being the only explanatory variable and then model parameters were tested in the corresponding sex-specific test subset. Model performance was evaluated using the Mean Absolute Error (MAE) and Root Mean Square Error (RMSE); the MAE is the average of the absolute differences (i.e., errors) between the predicted and the actual value of the outcome variable and the RMSE is the standard deviation of the prediction errors (iv) model optimization involves the inclusion in the models of covariates pertaining to acquisition and global neuroimaging features (i.e., ICV for SV models, mean cortical thickness for CT models and mean cortical surface area for SA models.

For each FreeSurfer-extracted measure, individualized z-scores were generated by subtracting the predicted value ($\hat{Y}$) from the observed value ($Y_o$) of that measure divided by the RMSE of the model[6-8].

**Proportion of subjects with infra- and supranormal regional normative z-scores and average deviation scores**

In order to identify the proportion of CHR-P with infra- and supranormal regional z-scores, for example, the percentage of CHR-P individuals with supranormal values for a region X was computed as: [(N of CHR-P individuals with supranormal values in X/N of CHR-P individuals)*100]. The same method was applied to identify the proportion of CHR-P with infranormal regional z-scores. Similarly, in order to identify the proportion of CHR-P with infra- or supranormal average deviation scores (ADS), for example, the percentage of CHR-P individuals with supranormal $ADS_{CT}$ was computed as: [(N of CHR-P individuals with supranormal ADSCT values/N of CHR-P individuals)*100]. This method was applied to identify both infra- and supranormal ADS for CT ($ADS_{CT}$), SA ($ADS_{SA}$), SV($ADS_{SV}$), and globally across all measures ($ADS_{G}$). The same was repeated in for all measures in healthy individuals.

**Alternate definitions of the average deviation scores**

We computed alternate average deviation scores for each neuroimaging phenotype and a global index based on the sign of the regional z-scores. Accordingly, regions with positive z-scores for cortical thickness were averaged to form the positive $ASD_{CT}$ (P-$ASD_{CT}$), and regions with negative z-scores for cortical thickness were averaged to form the negative $ASD_{CT}$ (N-$ASD_{CT}$). The same process was used to compute the P-$ASD_{SA}$ and N-$ASD_{SA}$ for cortical surface area, the P-$ASD_{SV}$ and N-$ASD_{SV}$ for the subcortical volumes, and P-$ADS_{G}$ and N-$ADS_{G}$ globally across all measures.

**eResults.**

**Distribution of regional normative z-scores and observed regional values in healthy individuals and individuals at clinical high-risk for psychosis**

eVideo. Distributions of regional normative z-scores and observed regional values in healthy individuals and individuals at clinical high-risk for psychosis (CHR-P), and individuals at clinical high-risk for psychosis that converted to full-blown psychosis (CHR-PC).

**Associations between average deviation scores with positive symptoms and IQ**

The association between ASDs with positive symptoms and IQ in CHR-P individuals was not influenced by medication status (eFigure 3A and B). The leave-one-site-out analyses replicated these findings with two exceptions: the association between $ASD_{SA}$ with positive symptoms was not present after excluding the Copenhagen site (11.79% of the CHR sample) and the association between $ADS_{SA}$ and IQ was not present after excluding the UCSF site (4.8% of the CHR sample) (eFigure 3C and D).

**CHR syndromes**

Of the 1340 CHR-P individuals, 1057 were classified as having attenuated symptoms (APS), 28 as having brief limited intermittent symptoms (BIPS) and 53 as having genetic risk and deterioration syndrome (GRD). As the sample size for BIPS and GRD was too small, the analyses described in the main manuscript were repeated for those with APS. The results recapitulated those reported for the total sample.

**Alternate average deviation scores**

We computed alternate average deviations scores by averaging separately positive or negative regional z-scores for the imaging phenotype (eMethods) and computed associations between each of these ADS and the positive symptom scores and IQ (eTable 18).

| eTable 1. Information on the samples contributed by site to the Clinical High-Risk for Psychosis Working Group of the ENIGMA Consortium | | | | | | | |
|---|---|---|---|---|---|---|---|
| **#** | **Full Site Name** | **Abbreviated Site Name** | **Major Eligibility Criteria for Healthy and CHR-P participants** | **Site sample (N) and mean age (SD) in years** | **Criteria for CHR-P status** | **Inclusion Criteria for Healthy Participants** | **IQ Method** |
| **1** | Vrije Universiteit Amsterdam, The Netherlands | Amsterdam | • 16-31 years<br>• Sufficient command of the Dutch language<br>• IQ ≥80 | Healthy=23; 23.45 (2.80)<br>CHR-P=16; 23.64 (2.51) | • Met CAARMS criteria | • CAARMS and SOFAS score <55<br>• No family history of psychiatric disorders | NA |
| **2** | New York State Psychiatric Institute, Columbia University, USA | Columbia1 | • 15-35 years<br>• No major medical or neurological disorder diagnosis<br>• IQ >70<br>• No lifetime substance use disorder<br>• No lifetime history of traumatic head injury<br>• No imminent risk of harm to self or others | Healthy=9; 24.35 (4.05)<br>CHR-P=17; 22.89 (4.96) | • Met SIPS criteria | • No SCID Axis I and/or Axis II Cluster C disorder diagnosis<br>• Participants Not adopted | NA |
| **3** | New York State Psychiatric Institute, Columbia University, USA | Columbia3 | • 14-30 years<br>• No lifetime major medical or neurological disorders<br>• IQ >70 | Healthy=17; 22.91 (3.68)<br>CHR-P=53; 21.27 (4.01) | • Met SIPS criteria<br>• CHR symptoms do not occur solely in the context of substance use | • No lifetime psychiatric disorder diagnosis<br>• No lifetime history of meeting criteria for psychosis-risk syndrome via the SIPS<br>• No current substance abuse | NA |
| **4** | Mental Health Center Copenhagen and CINS, Mental Health Center Glostrup, University of Copenhagen, Denmark | Copenhagen_1<br>Copenhagen_2 | • 18-40 years<br>• No organic brain disorder diagnosis (e.g., epilepsy, inflammatory brain disease)<br>• Fluent in Danish<br>• IQ >70<br>• Not currently pregnant | Healthy=58; 24.78 (3.30)<br>CHR-P=158; 24.21 (4.22) | • Met CAARMS criteria<br>• No psychiatric symptoms that could be explained by a physical illness or acute drug intoxication<br>• No diagnosis of a serious developmental disorder (e.g. Asperger's syndrome)<br>• Lifetime neuroleptic exposure equivalents less | • No lifetime DSM-IV psychiatric disorder diagnosis | WAIS-III |

| eTable 1. Information on the samples contributed by site to the Clinical High-Risk for Psychosis Working Group of the ENIGMA Consortium | | | | | | | |
|---|---|---|---|---|---|---|---|
| **#** | **Full Site Name** | **Abbreviated Site Name** | **Major Eligibility Criteria for Healthy and CHR-P participants** | **Site sample (N) and mean age (SD) in years** | **Criteria for CHR-P status** | **Inclusion Criteria for Healthy Participants** | **IQ Method** |
| | | | | | than haloperidol dose of >50 mg<br>• No current mood stabilizer use or recreational use of ketamine | | |
| **5** | Central South University, China | CSU | • 13-30 years<br>• No lifetime neurological disease or organic brain abnormalities (examined by neuroradiologist blinded to group allocation)<br>• Fluent in Chinese<br>• IQ >70<br>• Not currently pregnant<br>• No lifetime drug or alcohol dependence | Healthy=55; 21.47 (3.20)<br>CHR=49; 19.49 (5.05) | • Met SIPS criteria<br>• No lifetime psychotropic medication use | | WISC-I+PC |
| **6** | University Hospital of Child and Adolescent Psychiatry and Psychotherapy, University of Bern, Switzerland | FETZ Bern | • 8-40 years<br>• No current neurological disorder diagnosis<br>• Fluent in German, French or English<br>• IQ >70 (via clinical impression) | Healthy=15; 20.11 (6.02)<br>CHR-P=39; 19.14 (4.81) | • Met SIPS criteria | • No lifetime psychotic disorder diagnosis | NA |
| **7** | Institute of Neuroscience and Psychology, University of Glasgow, Scotland | Glasgow | • 16-35 years<br>• No organic cause for presentation<br>• Fluent in English<br>• IQ >70 | Healthy=45; 22.89 (3.61)<br>CHR-P=75; 22.45 (4.82) | • Met CAARMS criteria | • No current psychiatric disorder diagnosis<br>• No current substance abuse<br>• No first-degree relative with a psychotic disorder diagnosis | NART |

| eTable 1. Information on the samples contributed by site to the Clinical High-Risk for Psychosis Working Group of the ENIGMA Consortium | | | | | | | |
|---|---|---|---|---|---|---|---|
| **#** | **Full Site Name** | **Abbreviated Site Name** | **Major Eligibility Criteria for Healthy and CHR-P participants** | **Site sample (N) and mean age (SD) in years** | **Criteria for CHR-P status** | **Inclusion Criteria for Healthy Participants** | **IQ Method** |
| **8** | Heidelberg University Hospital, Germany | Heidelberg | • 14-18 years<br>• No lifetime neurological disorder diagnosis<br>• Fluent in German<br>• IQ >85<br>• No illegal drug use in past 12 months | Healthy=31; 15.71 (0.90)<br>CHR-P=22; 15.14 (1.08) | • Met SIPS criteria<br>• No lifetime DSM-IV Axis I diagnosis<br>• No psychotropic medication exposure | • No lifetime DSM-IV Axis-I disorder diagnosis<br>• No psychotropic medication prescription<br>• Does not meet APS psychosis risk syndrome via SIPS | WISC-IV |
| **9** | August Pi i Sunyer Biomedical Research Institute, Barcelona, Spain | IDIBAPS | • 10-17 years<br>• No lifetime neurological disorder or traumatic brain injury diagnosis<br>• Fluent in Spanish or Catalan<br>• IQ >70 | Healthy=44; 16.00 (1.53)<br>CHR-P=60; 15.32 (1.80) | • Met SIPS criteria<br>• No autism spectrum disorder | • No current psychiatric diagnosis<br>• No current psychotropic medication use<br>• No lifetime psychotic disorder<br>• No first- or second-degree relative with psychotic disorder diagnosis | WAIS-III<br>WAIS-IV<br>WISC-IV<br>WISC-V |
| **10** | Icahn School of Medicine at Mount Sinai, USA | ISMMS | 15-35 years<br>• No major medical or neurological disorder diagnosis<br>• No lifetime traumatic head injury<br>• Fluent in English<br>• IQ >70<br>• No MRI contraindications<br>• No lifetime substance use disorder<br>• No imminent risk of harm to self or others | Healthy=11; 27.89 (3.95)<br>CHR-P=24; 23.43 (5.55) | • Met SIPS criteria | • No SCID Axis I diagnosis, Axis II Cluster C disorder<br>• Participants not adopted | NA |

**eTable 1. Information on the samples contributed by site to the Clinical High-Risk for Psychosis Working Group of the ENIGMA Consortium**

| # | Full Site Name | Abbreviated Site Name | Major Eligibility Criteria for Healthy and CHR-P participants | Site sample (N) and mean age (SD) in years | Criteria for CHR-P status | Inclusion Criteria for Healthy Participants | IQ Method |
|---|---|---|---|---|---|---|---|
| **11** | Institute of Psychiatry, Psychology & Neuroscience, King's College London, UK | London | • 18-35 years<br>• No lifetime neurological disorder diagnosis<br>• Fluent in English<br>• IQ within normal range<br>• No drug or alcohol dependence, as specified in DSM-IV<br>• No recreational drug use in past two weeks | Healthy=12; 26.42 (4.78)<br>CHR-P=46; 23.52 (4.64) | • Met CAARMS criteria | • No lifetime psychiatric disorder diagnosis<br>• No current psychotropic medication use | NA |
| **12** | Maastricht University, The Netherlands | Maastricht | • 12-55 years<br>• No lifetime neurological disorder diagnosis<br>• Fluent in Dutch<br>• IQ >70 | Healthy=37; 25.52 (5.73)<br>CHR-P=39; 20.13 (4.18) | • Met SIPS criteria<br>• No lifetime psychotic disorder for more than one week, based on DSM-IV criteria<br>• Clear evidence that the psychosis-risk syndrome is not due to a non-schizophrenia-spectrum psychiatric disorder | • No DSM-IV psychiatric disorder diagnosis (lifetime)<br>• No psychotropic medication prescription (lifetime)<br>• No first- or second-degree relative with a psychotic disorder diagnosis | DART |
| **13** | University of Melbourne, Australia | Melbourne | • 14-30 years<br>• No history of substantial head injury, seizures, neurologic diseases, impaired thyroid function, corticosteroid use<br>• Fluent in English<br>• No alcohol or substance abuse or dependence | Healthy=90; 21.76 (3.59)<br>CHR-P=18; 18.39 (2.87) | • Met CAARMS criteria<br>• No known organic cause for presentation<br>• Lifetime neuroleptic exposure is less than total lifetime haloperidol use of >15mg | • No lifetime psychiatric disorder diagnosis<br>• No family history of psychotic illness | WASI<br>WASI-II |

| eTable 1. Information on the samples contributed by site to the Clinical High-Risk for Psychosis Working Group of the ENIGMA Consortium | | | | | | | |
|---|---|---|---|---|---|---|---|
| **#** | **Full Site Name** | **Abbreviated Site Name** | **Major Eligibility Criteria for Healthy and CHR-P participants** | **Site sample (N) and mean age (SD) in years** | **Criteria for CHR-P status** | **Inclusion Criteria for Healthy Participants** | **IQ Method** |
| **14** | Instituto Nacional de Neurología y Neurocirugía, Mexico City, Mexico | Mexico City | • 13-34 years<br>• No lifetime neurological disorder or traumatic brain injury diagnosis<br>• No lifetime drug or alcohol dependence | Healthy=37; 20.97 (3.4)<br>CHR-P=30; 19.7 (4.22) | • Met SIPS criteria | • No lifetime diagnosis of any psychiatric disorder<br>• No first- or second-degree relative with psychotic disorder | NA |
| **15** | Mental Health Research Center Moscow, Russia | MHRC | • 16-28 years<br>• Male<br>• Right-handed<br>• No lifetime neurological disorder diagnosis<br>• No lifetime drug or alcohol dependence | Healthy=33; 22.52 (2.52)<br>CHR=18; 20.03 (2.61) | • Met SIPS criteria<br>• No lifetime mental or behavioural disorders due to psychoactive substance use | • No lifetime psychiatric disorder<br>• No lifetime psychotropic medication prescription<br>• No family history of psychiatric or neurological disorders | NA |
| **16** | Maryland Psychiatric Research Center, University of Maryland School of Medicine, USA | MPRC | • 15-35 years<br>• No major medical or neurological disorder diagnosis<br>• No lifetime traumatic head injury<br>• Fluent in English<br>• IQ >70<br>• No MRI contraindications<br>• No lifetime substance use disorder<br>• No imminent risk of harm to self or others | Healthy=19; 18.00 (4.28)<br>CHR=29; 17.28 (3.21) | • Met SIPS criteria | • No lifetime psychiatric disorder | WASI<br>WASI-II |
| **17** | University of Newcastle, Australia | Newcastle | • 13-25 years<br>• No antipsychotic pharmacotherapy<br>• IQ ≥70<br>• No lifetime drug abuse or dependence | Healthy=17; 20.25 (1.91)<br>CHR-P=43; 19.53 (2.12) | • Met CAARMS criteria<br>• Drop of 30% in GAF in the past 12 months | • No psychiatric disorder diagnosis<br>• No lifetime treatment for depression or anxiety<br>• No first-degree family | WASI-II |

| eTable 1. Information on the samples contributed by site to the Clinical High-Risk for Psychosis Working Group of the ENIGMA Consortium | | | | | | | |
|---|---|---|---|---|---|---|---|
| **#** | **Full Site Name** | **Abbreviated Site Name** | **Major Eligibility Criteria for Healthy and CHR-P participants** | **Site sample (N) and mean age (SD) in years** | **Criteria for CHR-P status** | **Inclusion Criteria for Healthy Participants** | **IQ Method** |
| | | | diagnosis<br>• No head injury with loss of consciousness (>15 mins)<br>• No organic brain impairment<br>• No hearing impairment (>20dB SPL)<br>• No history of nasal trauma. | | | member with schizophrenia | |
| **18** | NORMENT, University of Oslo and Oslo University Hospital, Norway | Oslo Region | • 15-30 years<br>• No organic cause for presentation<br>• Fluent in Norwegian<br>• IQ >70 | Healthy=62; 19.90 (3.62) CHR-P=20; 20.08 (3.61) | • Met SIPS criteria<br>• No antipsychotic medication equivalent to a dose of ≥5 mg Olanzapine per day (current or ≥4 weeks lifetime)<br>• CHR symptoms not substance-induced | • No lifetime psychiatric disorder diagnosis<br>• No lifetime psychotropic medication use<br>• No lifetime psychotic disorder diagnosis<br>• No first-degree relative with psychotic disorder diagnosis<br>• No lifetime substance abuse disorder diagnosis | WASI |
| **19** | University of Pittsburgh, USA | Pitt | • 12-40 years<br>• No lifetime neurological disorder diagnosis<br>• Fluent in English<br>• IQ >80 | Healthy=60; 22.93 (5.53) CHR-P=25; 20.87 (5.4) | • Met SIPS criteria<br>• psychosis-risk syndrome not due to a non-schizophrenia-spectrum psychiatric disorder | • No lifetime disorder diagnosis<br>• No lifetime psychotropic medication exposure<br>• No lifetime psychosis risk syndrome (SIPS criteria)<br>• No first-degree relative with a psychotic disorder diagnosis | WASI |

| **eTable 1. Information on the samples contributed by site to the Clinical High-Risk for Psychosis Working Group of the ENIGMA Consortium** | | | | | | | |
|---|---|---|---|---|---|---|---|
| **#** | **Full Site Name** | **Abbreviated Site Name** | **Major Eligibility Criteria for Healthy and CHR-P participants** | **Site sample (N) and mean age (SD) in years** | **Criteria for CHR-P status** | **Inclusion Criteria for Healthy Participants** | **IQ Method** |
| **20** | Rush University Medical Center, Chicago, IL USA | RUMC | • 12-35 years<br>• No lifetime major medical illness or neurological disorder diagnosis that could confound behavioral or neural measurements, incl. diabetes, epilepsy, multiple sclerosis, or traumatic brain injury with severity 7+ on TBI scale<br>• IQ >70 | Healthy=29; 23.88 (3.22)<br>CHR-P=62; 22.74 (3.93) | • Met SIPS criteria<br>• Psychosis-risk syndrome not due to a non-schizophrenia-spectrum psychiatric disorder | • No current Axis I/II disorders<br>• Positive symptoms rated 1 or lower on the SIPS | WASI<br>WRAT |
| **21** | Institute of Mental Health, Singapore and National University of Singapore, Singapore | Singapore | • 14-29 years<br>• No lifetime neurological disorder diagnosis<br>• Fluent in English<br>• No history of intellectual disability<br>• No lifetime history of illicit substance use | Healthy=52; 22.01 (4.21)<br>CHR-P=99; 21.93 (3.59) | • Met CAARMS criteria<br>• No current antipsychotic or mood stabilizer use<br>• No medical cause associated with psychotic symptoms | • No lifetime psychiatric disorder diagnosis<br>• No first-degree relative with a psychotic disorder diagnosis | NA |
| **22** | Seoul National University Hospital, South Korea | SNUH | • 15-34 years<br>• Female or male<br>• No lifetime neurological disorder diagnosis or traumatic brain injury<br>• Fluent in Korean<br>• IQ >70<br>• No MRI contraindications | Healthy=72; 21.25 (2.51)<br>CHR-P=71; 20.72 (3.81) | • Met SIPS criteria | • No lifetime psychiatric disorder diagnosis, including psychotic disorder diagnosis<br>• No current psychotropic medication prescription<br>• No first- or second-degree relative with a psychotic disorder diagnosis<br>• No lifetime neurological disorder diagnosis | K-WAIS |

| **eTable 1. Information on the samples contributed by site to the Clinical High-Risk for Psychosis Working Group of the ENIGMA Consortium** | | | | | | | |
|---|---|---|---|---|---|---|---|
| **#** | **Full Site Name** | **Abbreviated Site Name** | **Major Eligibility Criteria for Healthy and CHR-P participants** | **Site sample (N) and mean age (SD) in years** | **Criteria for CHR-P status** | **Inclusion Criteria for Healthy Participants** | **IQ Method** |
| **23** | Stavanger University Hospital, Norway | Stavanger | • 13-65 years<br>• Female, male<br>• No known neurological or endocrine disorders related to CHR symptoms<br>• Able to speak a Scandinavian language<br>• IQ >70<br>• No MRI contraindications | Healthy=33; 17.03 (3.10)<br>CHR=P=35; 16.26 (1.88) | • Met SIPS criteria<br>• Symptoms not better accounted for by major Axis I, Axis II , or substance use disorder<br>• No current antipsychotic medication use<br>• Not more than 4 weeks of lifetime antipsychotic medication use | • No lifetime psychiatric disorder diagnosis<br>• No lifetime psychotropic medication exposure<br>• No lifetime psychosis risk syndrome (SIPS criteria)<br>• No first-degree relatives with a psychotic disorder diagnosis<br>• No current substance use disorder | WAIS-IV |
| **24** | Department of Neuro-psychiatry, Toho University School of Medicine, Japan | Toho | • 16-35 years<br>• No lifetime neurological disorder diagnosis<br>• Fluent in Japanese<br>• IQ >70<br>• No MRI contraindications | Healthy=15; 22.87 (2.64)<br>CHR=P=34; 23.71 (6.9) | • Met SIPS criteria | • No lifetime psychiatric disorder diagnosis<br>• No psychotropic medication exposure | WAIS-III |
| **25** | Department of Neuropsychiatry Graduate School of Medicine, The University of Tokyo | Tokyo | • 15-30 years old<br>• No neurological disorder diagnosis or traumatic brain injury with known cognitive consequences and/or loss of consciousness >5 mins<br>• Fluent in Japanese<br>• IQ ≥70<br>• No substance use disorder or substance abuse<br>• No history of | Healthy=25; 22.08 (2.84)<br>CHR-P=38; 20.92 (3.55) | • Met SIPS criteria | • No psychiatric condition<br>• No psychotropic medication use<br>• No first-degree relatives with a psychotic disorder diagnosis | JART25 |

| eTable 1. Information on the samples contributed by site to the Clinical High-Risk for Psychosis Working Group of the ENIGMA Consortium | | | | | | | |
|---|---|---|---|---|---|---|---|
| # | Full Site Name | Abbreviated Site Name | Major Eligibility Criteria for Healthy and CHR-P participants | Site sample (N) and mean age (SD) in years | Criteria for CHR-P status | Inclusion Criteria for Healthy Participants | IQ Method |
| | | | electroconvulsive therapy | | | | |
| **26** | Centre for Addiction and Mental Health, University of Toronto, Canada | Toronto | • 18-40 years<br>• No current neurological disorder diagnosis<br>• Fluent in English or French<br>• IQ >70 (via clinical impression)<br>• Not currently pregnant<br>• No current drug or alcohol abuse<br>• No lifetime history of severe head trauma | Healthy=36; 25.37 (5.1)<br>CHR-P=25; 20.84 (1.9) | • Met SIPS criteria | • No lifetime psychiatric disorder diagnosis<br>• No psychotropic medication prescription<br>• No relatives with a psychiatric disorder diagnosis | RBANS |
| **27** | Graduate School of Medicine and Pharmaceutical Sciences, University of Toyama, Japan | Toyama | • 12-42 years<br>• No lifetime neurological disorder diagnosis<br>• Fluent in Japanese<br>• No MRI contraindications | Healthy =139; 25.04 (4.24)<br>CHR-P=73; 18.59 (4.11) | • Met CAARMS criteria | • No lifetime psychiatric disorder diagnosis<br>• No lifetime psychotropic medication exposure<br>• No first-degree relative with a psychiatric disorder diagnosis<br>• Physically healthy at time of MRI scanning<br>• No history of severe obstetric complications, serious head trauma, serious medical disease (e.g. neurological illness, thyroid dysfunction, diabetes, and | JART50 |

| eTable 1. Information on the samples contributed by site to the Clinical High-Risk for Psychosis Working Group of the ENIGMA Consortium | | | | | | | |
|---|---|---|---|---|---|---|---|
| **#** | **Full Site Name** | **Abbreviated Site Name** | **Major Eligibility Criteria for Healthy and CHR-P participants** | **Site sample (N) and mean age (SD) in years** | **Criteria for CHR-P status** | **Inclusion Criteria for Healthy Participants** | **IQ Method** |
| | | | | | | hypertension), steroid use, or substance abuse. | |
| **28** | University of California, San Francisco, USA | UCSF | • 11-30 years<br>• No lifetime neurological disorder diagnosis<br>• Fluent in English<br>• No MRI contraindications<br>• No substance dependence (past 12 months, excluding nicotine)<br>• No head injury<br>• Good physical health | Healthy=100;<br>23.9 (7.6)<br>CHR-P=65;<br>19.55 (4.46) | • Met SIPS criteria | • No lifetime DSM-IV Axis I disorder diagnosis<br>• No first-degree relatives with a psychotic disorder diagnosis | WASI-II |
| **29** | Psychiatric Hospital, University of Zurich, Switzerland | Zurich | • 13-35 years<br>• No neurological disorder diagnosis, no organic brain abnormalities (confirmed by neuroradiologist blinded to group allocation)<br>• Fluent in German<br>• IQ ≥80<br>• No pregnancy<br>• No drug or alcohol dependence | Healthy=43;<br>22.23 (5.56)<br>CHR-P=57;<br>19.37 (4.97) | • Met SIPS criteria<br>• No other psychiatric disorder diagnosis | • No lifetime psychiatric diagnosis<br>• No psychotropic medication use<br>• No first-degree relatives with a psychotic disorder diagnosis | MWT-B<br>WISC |

**eTable 2. SIPS and CAARMS items and criteria for Clinical High-Risk status for Psychosis**

| | SIPS | CAARMS |
|---|---|---|
| Subscales | P1: Unusual Thought Content/Delusional Ideas | P1: Unusual Thought Content |
| | P2: Suspiciousness/Persecutory Ideas | P2: Non-Bizarre Ideas |
| | P3: Grandiose Ideas | |
| | P4: Perceptual Abnormalities/Hallucinations | P3: Perceptual Abnormalities |
| | P5: Disorganized Communication | P4: Disorganized Speech |
| Frequency | 1: at least several minutes per day at least 1/month<br>2: several minutes/day at least once/week in the past month<br>3: at least 1 hour/day for at least 4 days/week over 1 month | 0: absent<br>1: less than 1/month<br>2: 1/month to 2/weeks, <1 hour per occasion<br>3: 1/month to 2/weeks, >1 hour per occasion, OR 3 to 6/weeks <1 h per occasion<br>4: 3 to 6/week, > 1 hour per occasion, OR daily, <1 hour per occasion<br>5: daily, >1 hour per occasion, OR several times/ day<br>6: continuous |
| Substance use | Exclusion criterion if strongly intertwined with symptoms | 0: no relation to substance use noted<br>1: occurs in relation to substance use and at other times as well<br>2: noted only in relation to substance use |
| Distress | Subjective qualifier<br>Not used to determine an individual’s status | Rated on a scale of 0–100<br>Not used to determine an individual’s status |
| **Attenuated psychotic symptoms** | | |
| Inclusion criteria | Severity score of 3–5 on at least one of P1–P5<br>PLUS<br>Frequency score of 2 on P1, P2, P3, P4, and/or P5 | Subthreshold intensity<br>Severity score of 3–5 on P1, 3–5 on P2, 3-4 on P3, and/or 4-5 on P4<br>PLUS<br>Frequency score of 3–6 on P1, P2, P3, and/or P4<br>Subthreshold frequency<br>Severity score of 6 on at least one of P1, P2, and P4 and/or 5-6 on P3<br>PLUS<br>Frequency score of 3 on P1, P2, P3, and/or P4 |
| Onset | Symptoms should have begun within the past year OR currently rate one or more scale points higher compared to 12 months before<br>Symptoms that occurred over the past month only are rated | Symptoms should have been present in the previous 12 months and for not longer than 5 years |
| Level of functioning | No social/occupational dysfunction requirement | 30% drop in SOFAS score from premorbid level, sustained for a month, within the past 12 months OR SOFAS score <50 for the past 12 months or more |
| Exclusion criteria | Symptoms are strongly intertwined temporally with substance use episodes (substance-induced psychosis may be considered)<br>Symptoms are better accounted for by another DSM diagnosis<br>Past psychosis was ruled in according to information obtained through the initial screen and evaluated using the POPS | Symptoms occur only during peak intoxication from a substance known to be associated with psychotic experiences (e.g., hallucinogens, amphetamines, and cocaine)<br>—<br>The person has had a previous psychotic episode (treated or untreated) |

**eTable 2. SIPS and CAARMS items and criteria for Clinical High-Risk status for Psychosis**

| | SIPS | CAARMS |
|---|---|---|
| **Brief intermittent psychotic symptoms** | | |
| Inclusion criteria | Severity score of 6 on at least one of P1–P5<br>PLUS<br>Frequency score of 1 on P1, P2, P3, P4, and/or P5 | Severity score of 6 on at least one of P1, P2, and P4 and/or 5-6 on P3<br>PLUS<br>Frequency score of 4–6 on P1, P2, P3, and/or P4 |
| Onset | Symptoms should have reached a psychotic level of intensity in the previous 3 month | Symptoms should have been present in the previous 12 months and for not longer than 5 years |
| Duration | Up to 3 months | Up to 7 days |
| Level of functioning | No social/occupational dysfunction requirement | 30% drop in SOFAS score from premorbid level, sustained for a month, within the past 12 months<br>OR<br>SOFAS score <50 for the past 12 months or more |
| Exclusion criteria | Symptoms are strongly intertwined temporally with substance use episodes (substance-induced psychosis may be considered)<br>Symptoms are better accounted for by another DSM diagnosis<br>Past psychosis was ruled in according to information obtained through the initial screen and evaluated using the POPS<br>Symptoms are seriously disorganizing and dangerous | Symptoms occur only during peak intoxication from a substance known to be associated with psychotic experiences (e.g., hallucinogens, amphetamines, and cocaine)<br>—<br>Previous psychotic episode (treated or untreated)<br>—<br>Symptoms do not resolve spontaneously (without antipsychotic medication) |
| **Genetic risk and deterioration syndrome** | | |
| Inclusion criteria | Meets criteria for Schizotypal Personality Disorder<br>OR<br>Has a first-degree relative with a psychotic disorder | Schizotypal Personality Disorder in identified patient<br>OR<br>Family history of psychosis in a first-degree relative |
| Level of functioning | 30% drop in GAF score over the last month as compared to 12 months before | 30% drop in SOFAS score from premorbid level, sustained for a month, within the past 12 months OR SOFAS score <50 for the past 12 months or more |

CAARMS=Comprehensive Assessment of At-Risk Mental States; d=day; GAF=Global Assessment of Functioning; SIPS=Structured Interview for Psychosis-Risk Syndrome; SOFAS=Social and Occupational Functioning Assessment Scale

**eTable 3. Neuroimaging acquisition protocols and FreeSurfer version per site**

| Site | Scanner Vendor | Scanner Model | Magnet Strength (Tesla) | Repetition Time (ms) | Echo Time (ms) | Flip Angle | Voxel Size (mm) | FreeSurfer version |
|---|---|---|---|---|---|---|---|---|
| Amsterdam | Philips | Image MR Series | 3 | 8280 | 3.8 | • | • | 6.0.0 |
| Columbia1 | GE | Discovery MR750 | 3 | 7840 | 3.1 | 12° | 0.8x0.8x0.8 | 6.0.0 |
| Columbia3 | GE | Signa | 3 | • | • | 11° | 0.98x0.98x1.0 | 5.3 |
| Copenhagen | Philips | Achieva | 3 | 10.028 | 4.6 | 8° | 0.75x0.75x0.80 | 6.0.0 |
| | | Achieva | 3 | 10.01 | 4.6 | 8° | 0.75x0.75x0.80 | 6.0.0 |
| CSU | Siemens | Skyra | 3 | 2530 | 2.33 | 7° | 1.0x1.0x1.0 | 6.0.0 |
| Fetz Bern | Siemens | Magnetom Verio | 3 | 7920 | 2.48 | 16° | 1.0x1.0x1.0 | 6.0.0 |
| Glasgow | Siemens | Trio | 3 | 2250 | 2.6 | 9° | 1.0x1.0x1.0 | 6.0.0 |
| Heidelberg | Siemens | PET/MR | 3 | 2300 | 2.98 | 9° | 1.0x1.0x1.0 | 6.0.0 |
| | | Tim Trio | 3 | 2300 | 2.98 | 9° | 1.0x1.0x1.0 | 6.0.0 |
| IDIBAPS | Siemens | Tim Trio | 3 | 2300 | 3.01 | 9° | 0.94x0.94x1.0 | 6.0.0 |
| | | Prisma fit | 3 | 2300 | 3.01 | 9° | 0.94x0.94x1.0 | 6.0.0 |
| | | Prisma fit | 3 | 2300 | 2.98 | 9° | 1.0x1.0x1.2 | 6.0.0 |
| London | GE | Signa HDx | 3 | 7.144 | 2.85 | 20° | 1.1x1.1.x1.1 | 6.0.0 |
| Maastricht | Philips | Intera | 3 | 2250 | 4.6 | 8° | 1.17×1.17×1.20 | 6.0.0 |
| | | Achieva | 3 | 2250 | 4.6 | 8° | 1.2x0.8x0.8 | 6.0.0 |
| Melbourne | Siemens | Trio | 3 | 1900 | 2.15 | 90° | 0.5x0.5x1.0 | 6.0.0 |
| | | Trio | 3 | 2300 | 2.98 | 9° | 1.0x1.0x1.2 | 6.0.0 |
| | GE | Signa | 1.5 | 14.3 | 3.3 | 30° | 0.94x0.94x1.5 | 6.0.0 |
| | | LX Horizon | 3 | 36 | 9 | 30° | 0.49x0.49x2 | 6.0.0 |
| Mexico City | GE | Signa HDxt | 3 | 1340 | 5.7 | 20° | 1.2x1.2x1.2 | 6.0.0 |

| eTable 3. Neuroimaging acquisition protocols and FreeSurfer version per site | | | | | | | | |
|---|---|---|---|---|---|---|---|---|
| **Site** | **Scanner Vendor** | **Scanner Model** | **Magnet Strength (Tesla)** | **Repetition Time (ms)** | **Echo Time (ms)** | **Flip Angle** | **Voxel Size (mm)** | **FreeSurfer version** |
| MHRC | Philips | Achieva | 3 | 8.2 | 3.7 | 8° | 0.83×0.83x1.0 | 6.0.0 |
| MPRC | Siemens | Trio | 3 | 2400 | 2.2 | 8° | 0.8x0.8x0.8 | 6.0.0 |
| | | Prisma | 3 | 2400 | 2.2 | 8° | 0.8x0.8x0.8 | 6.0.0 |
| ISMMS | Siemens | Skyra | 3 | 2400 | 2.07 | 8° | 0.8x0.8x0.8 | 6.0.0 |
| Newcastle | Siemens | Avanto | 1.5 | 1980 | 4.3 | 15° | 0.98x0.98x1 | 6.0.0 |
| Oslo Region | GE | Signa HDxt | 3 | 7800 | 2.96 | 12° | 1.0×1.0×1.2 | 5.3 |
| | | Discovery MR750 | 3 | 8.16 | 3.18 | 12° | 1.0x1.0x1.0 | 5.3 |
| Pitt | Siemens | Prisma | 3 | 2400 | 2.2 | 8° | 0.8x0.8x0.8 | 6.0.0 |
| RUMC | Siemens | Verio | 3 | 2530 | 2.27 | 7° | 1.0x1.0x1.0 | 6.0.0 |
| Singapore | Siemens | Tim Trio | 3 | 2300 | 3 | 9° | 1.0x1.0x1.0 | 6.0.0 |
| SNUH | Siemens | Magnetom TrioTim | 3 | 1670 | 1.89 | 9° | 1x0.98x0.98 | 6.0.0 |
| Stavanger | GE | Discovery 450 | 1.5 | 7.9 | 3.1 | 12° | • | 5.3 |
| Toho | Toshiba | Excelart Vantage | 1.5 | • | • | 35° | 0.98×0.98×1.0 | 5.2 |
| Tokyo | GE | SIGNA HDx | 3 | 6.8 | 1.94 | 20° | 1.0x1.0x1.0 | 6.0.0 |
| | | Discovery MR750W | 3 | 8.46 | 3.25 | 20° | 1.0x1.0x1.0 | 6.0.0 |
| Toronto | GE | Discovery MR750 | 3 | 6736 | 2.99 | 8° | 0.9x0.9x0.9 | 6.0.0 |
| Toyama | Siemens | Magnetom Vision | 1.5 | 2400 | 5 | 40° | 1.0x1.0x1.0 | 6.0.0 |
| | | Magnetom Verio | 3 | 2300 | 2.9 | 9° | 1.0x1.0x1.2 | 6.0.0 |
| UCSF | Siemens | Magnetom TrioTim | 3 | 2300 | 2.95 | 9° | 1.0x1.0x1.2 | 5.1 |

| eTable 3. Neuroimaging acquisition protocols and FreeSurfer version per site | | | | | | | | |
|---|---|---|---|---|---|---|---|---|
| **Site** | **Scanner Vendor** | **Scanner Model** | **Magnet Strength (Tesla)** | **Repetition Time (ms)** | **Echo Time (ms)** | **Flip Angle** | **Voxel Size (mm)** | **FreeSurfer version** |
| Zurich | Philips | Achieva TX | 3 | 8.3 | 3.8 | 8° | 1.0x1.0x1.0 | 6.0.0 |
| Unavailable information left blank | | | | | | | | |

| eTable 4. FreeSurfer-derived morphometric measures | | |
|---|---|---|
| **Measure** | **Side** | **Regions** |
| **Cortical Thickness** | **L** | left_bankssts, left_caudalanteriorcingulate, left_caudalmiddlefrontal, left_cuneus, left_entorhinal, left_fusiform, left_inferiorparietal, left_inferiortemporal, left_isthmuscingulate, left_lateraloccipital, left_lateralorbitofrontal, left_lingual, left_medialorbitofrontal, left_middletemporal, left_parahippocampal, left_paracentral, left_parsopercularis, left_parsorbitalis, left_parstriangularis, left_pericalcarine, left_postcentral, left_posteriorcingulate, left_precentral, left_precuneus, left_rostralanteriorcingulate, left_rostralmiddlefrontal, left_superiorfrontal, left_superiorparietal, left_superiortemporal, left_supramarginal, left_frontalpole, left_temporalpole, left_transversetemporal, left_insula |
| | **R** | right_bankssts, right_caudalanteriorcingulate, right_caudalmiddlefrontal, right_cuneus, right_entorhinal, right_fusiform, right_inferiorparietal, right_inferiortemporal, right_isthmuscingulate, right_lateraloccipital, right_lateralorbitofrontal, right_lingual, right_medialorbitofrontal, right_middletemporal, right_parahippocampal, right_paracentral, right_parsopercularis, right_parsorbitalis, right_parstriangularis, right_pericalcarine, right_postcentral, right_posteriorcingulate, right_precentral, right_precuneus, right_rostralanteriorcingulate, right_rostralmiddlefrontal, right_superiorfrontal, right_superiorparietal, right_superiortemporal, right_supramarginal, right_frontalpole, right_temporalpole, right_transversetemporal, right_insula |
| **Surface Area** | **L** | left_bankssts, left_caudalanteriorcingulate, left_caudalmiddlefrontal, left_cuneus, left_entorhinal, left_fusiform, left_inferiorparietal, left_inferiortemporal, left_isthmuscingulate, left_lateraloccipital, left_lateralorbitofrontal, left_lingual, left_medialorbitofrontal, left_middletemporal, left_parahippocampal, left_paracentral, left_parsopercularis, left_parsorbitalis, left_parstriangularis, left_pericalcarine, left_postcentral, left_posteriorcingulate, left_precentral, left_precuneus, left_rostralanteriorcingulate, left_rostralmiddlefrontal, left_superiorfrontal, left_superiorparietal, left_superiortemporal, left_supramarginal, left_frontalpole, left_temporalpole, left_transversetemporal, left_insula |
| | **R** | right_bankssts, right_caudalanteriorcingulate, right_caudalmiddlefrontal, right_cuneus, right_entorhinal, right_fusiform, right_inferiorparietal, right_inferiortemporal, right_isthmuscingulate, right_lateraloccipital, right_lateralorbitofrontal, right_lingual, right_medialorbitofrontal, right_middletemporal, right_parahippocampal, right_paracentral, right_parsopercularis, right_parsorbitalis, right_parstriangularis, right_pericalcarine, right_postcentral, right_posteriorcingulate, right_precentral, right_precuneus, right_rostralanteriorcingulate, right_rostralmiddlefrontal, right_superiorfrontal, right_superiorparietal, right_superiortemporal, right_supramarginal, right_frontalpole, right_temporalpole, right_transversetemporal, right_insula |
| **Subcortical Volumes** | **L** | Left Thalamus, Left Caudate, Left Putamen, Left Pallidum, Left Hippocampus, Left Amygdala, Left Accumbens area |
| | **R** | Right Thalamus, Right Caudate, Right Putamen, Right Pallidum, Right Hippocampus, Right Amygdala, Right Accumbens area |

| eTable 5. Demographic characteristics of the participants in the study sample per site | | | | | | | |
|---|---|---|---|---|---|---|---|
| **#** | **Site** | **Healthy Individuals (N = 1237)** | | | **Clinical High-Risk for Psychosis (N = 1340)** | | |
| | | **N** | **Age (years) Mean (SD)** | **Sex # Male (%)** | **N** | **Age (years) Mean (SD)** | **Sex # Male (%)** |
| **1** | Amsterdam | 23 | 23.45 (2.80) | 10 (43.5 %) | 16 | 23.64 (2.51) | 6 (37.5 %) |
| **2** | Columbia1 | 9 | 24.35 (4.05) | 7 (77.8 %) | 17 | 22.89 (4.96) | 8 (47.1 %) |
| **3** | Columbia3 | 35 | 22.91 (3.68) | 23 (65.7 %) | 53 | 21.27 (4.01) | 39 (73.6 %) |
| **4** | Copenhagen | 58 | 24.78 (3.30) | 29 (50.0 %) | 158 | 24.21 (4.22) | 72 (45.6 %) |
| **5** | CSU | 55 | 21.47 (3.20) | 30 (54.5 %) | 49 | 19.49 (5.05) | 26 (53.1 %) |
| **6** | Bern | 15 | 20.11 (6.02) | 10 (66.7 %) | 39 | 19.14 (4.81) | 20 (51.3 %) |
| **7** | Glasgow | 45 | 22.89 (3.61) | 15 (33.3 %) | 75 | 22.45 (4.82) | 15 (20.0 %) |
| **8** | Heidelberg | 31 | 15.71 (0.90) | 15 (48.4 %) | 22 | 15.14 (1.08) | 9 (40.9 %) |
| **9** | IDIBAPS | 44 | 16.00 (1.53) | 14 (31.8 %) | 60 | 15.32 (1.80) | 20 (33.3 %) |
| **10** | ISMMS | 11 | 27.89 (3.95) | 6 (54.5 %) | 24 | 23.43 (5.55) | 11 (45.8 %) |
| **11** | London_1 | 12 | 26.42 (4.78) | 7 (58.3 %) | 46 | 23.52 (4.64) | 35 (76.1 %) |
| **12** | Maastricht | 37 | 25.52 (5.73) | 26 (70.3 %) | 39 | 20.13 (4.18) | 27 (69.2 %) |
| **13** | Melbourne | 90 | 21.76 (3.59) | 48 (53.3 %) | 18 | 18.39 (2.87) | 7 (38.9 %) |
| **14** | Mexico City | 37 | 20.97 (3.40) | 28 (75.7 %) | 30 | 19.70 (4.22) | 24 (80.0 %) |
| **15** | MHRC | 33 | 22.52 (2.52) | 33 (100.0 %) | 18 | 20.03 (2.61) | 18 (100.0 %) |
| **16** | MPRC | 19 | 18.00 (4.28) | 11 (57.9 %) | 29 | 17.28 (3.21) | 14 (48.3 %) |
| **17** | Newcastle | 17 | 20.25 (1.91) | 4 (23.5 %) | 43 | 19.53 (2.12) | 20 (46.5 %) |
| **18** | Oslo Region | 62 | 19.90 (3.62) | 39 (62.9 %) | 20 | 20.08 (3.61) | 12 (60.0 %) |
| **19** | Pitt | 60 | 22.93 (5.53) | 35 (58.3 %) | 25 | 20.87 (5.40) | 11 (44.0 %) |
| **20** | RUMC | 29 | 23.88 (3.22) | 10 (34.5 %) | 62 | 22.74 (3.93) | 33 (53.2 %) |
| **21** | Singapore | 52 | 22.01 (4.21) | 27 (51.9 %) | 99 | 21.93 (3.59) | 68 (68.7 %) |
| **22** | SNUH | 72 | 21.25 (2.51) | 49 (68.1 %) | 71 | 20.72 (3.81) | 52 (73.2 %) |
| **23** | Stavanger | 33 | 17.03 (3.10) | 17 (51.5 %) | 35 | 16.26 (1.88) | 13 (37.1 %) |
| **24** | Toho | 15 | 22.87 (2.64) | 8 (53.3 %) | 34 | 23.71 (6.90) | 7 (20.6 %) |
| **25** | Tokyo | 25 | 22.08 (2.84) | 13 (52.0 %) | 38 | 20.92 (3.55) | 20 (52.6 %) |
| **26** | Toronto | 36 | 25.37 (5.10) | 20 (55.6 %) | 25 | 20.84 (1.90) | 14 (56.0 %) |
| **27** | Toyama | 139 | 25.04 (4.24) | 72 (51.8 %) | 73 | 18.59 (4.11) | 40 (54.8 %) |
| **28** | UCSF | 100 | 23.90 (7.60) | 57 (57.0 %) | 65 | 19.55 (4.46) | 35 (53.8 %) |
| **29** | Zurich | 43 | 22.23 (5.56) | 21 (48.8 %) | 57 | 19.37 (4.97) | 33 (57.9 %) |

| **eTable 6. Characteristics of Clinical High-Risk for Psychosis (CHR-P) individuals in the study sample defined with either the SIPS or CAARMS criteria** | | | | |
|---|---|---|---|---|
| | **CHR-P individuals meeting SIPS criteria** | | **CHR-P individuals meeting CAARMS criteria** | |
| | **N** | | **N** | |
| **Age in years, mean (SD)** | 806 | 19.90 (4.69) | 534 | 22.03 (4.52) |
| **Sex, number male (%)** | 806 | 444 (55.09%) | 534 | 265 (49.63%) |
| **Positive symptoms score, mean (SD)** | 806 | 10.93 (4.66) | 534 | 10.37 (4.03) |
| **Prescribed typical antipsychotics, number (%)** | 744 | 21 (2.82%) | 526 | 1 (0.19%) |
| **Prescribed atypical antipsychotics, number (%)** | 748 | 169 (22.59%) | 526 | 62 (11.79) |
| **Converters, N (%)** | 641 | 115 (17.94%) | 456 | 42 (9.21%) |
| **Follow-up period in months, mean (SD)[a]** | 597 | 19.07 (10.67) | 392 | 20.60 (17.54) |

[a]The duration of follow-up was unavailable in 108 CHR-P individuals.

| eTable 7. Percentage of individuals with infra- or supranormal normative regional z-scores based on group | | | | | |
|---|---|---|---|---|---|
| | | Normative z-scores in HI | | Normative z-scores in CHR-P | |
| Region | Hemi | infranormal z-scores (%) | supranormal z-scores (%) | infranormal z-scores (%) | supranormal z-scores (%) |
| **Subcortical Volume** | | | | | |
| nucleus accumbens | L | 4.85 | 0.49 | 5.82 | 0.52 |
| | R | 5.17 | 0.40 | 7.01 | 0.07 |
| amygdala | L | 3.31 | 0.40 | 3.73 | 0.60 |
| | R | 2.67 | 1.37 | 3.73 | 2.01 |
| caudate | L | 3.31 | 0.89 | 5.97 | 0.60 |
| | R | 2.75 | 0.81 | 4.93 | 1.12 |
| hippocampus | L | 3.07 | 0.24 | 4.78 | 0.75 |
| | R | 3.07 | 0.65 | 3.88 | 0.67 |
| pallidum | L | 3.96 | 0.40 | 5.67 | 0.37 |
| | R | 4.20 | 0.81 | 6.04 | 0.97 |
| putamen | L | 8.57 | 0.40 | 10.00 | 0.37 |
| | R | 9.30 | 0.24 | 11.42 | 0.22 |
| thalamus | L | 6.95 | 0.08 | 8.28 | 0.82 |
| | R | 6.22 | 0.57 | 8.58 | 0.30 |
| **Cortical Thickness** | | | | | |
| banks superior temporal sulcus | L | 2.26 | 2.34 | 2.31 | 1.57 |
| | R | 2.75 | 2.18 | 3.43 | 2.24 |
| caudal anterior cingulate | L | 2.67 | 4.45 | 2.54 | 3.13 |
| | R | 3.15 | 4.37 | 3.66 | 4.40 |
| caudal middle frontal | L | 2.83 | 1.37 | 2.46 | 1.12 |
| | R | 2.83 | 1.54 | 2.84 | 1.64 |
| cuneus | L | 1.37 | 1.94 | 1.87 | 2.31 |
| | R | 0.81 | 2.51 | 1.87 | 3.06 |
| entorhinal cortex | L | 1.46 | 3.31 | 2.46 | 4.18 |
| | R | 2.02 | 2.67 | 2.69 | 3.66 |
| fusiform gyrus | L | 1.78 | 2.43 | 1.42 | 2.99 |
| | R | 1.94 | 2.43 | 1.87 | 3.13 |
| inferior parietal | L | 1.21 | 2.34 | 1.57 | 2.61 |
| | R | 1.94 | 1.21 | 2.09 | 1.94 |
| inferior temporal | L | 1.13 | 2.75 | 1.27 | 3.28 |
| | R | 0.89 | 2.51 | 1.94 | 4.85 |
| isthmus cingulate | L | 4.20 | 1.37 | 5.52 | 1.64 |
| | R | 4.61 | 2.43 | 4.85 | 1.49 |
| lateral occipital | L | 1.21 | 5.50 | 0.60 | 4.78 |
| | R | 0.57 | 4.45 | 0.90 | 4.85 |
| lateral orbitofrontal | L | 2.91 | 1.94 | 1.64 | 1.79 |
| | R | 1.37 | 2.59 | 1.64 | 2.31 |
| lingual gyrus | L | 2.99 | 2.59 | 3.51 | 2.09 |
| | R | 2.18 | 2.34 | 2.16 | 2.24 |

| eTable 7. Percentage of individuals with infra- or supranormal normative regional z-scores based on group | | | | | |
|---|---|---|---|---|---|
| | | Normative z-scores in HI | | Normative z-scores in CHR-P | |
| Region | Hemi | infranormal z-scores (%) | supranormal z-scores (%) | infranormal z-scores (%) | supranormal z-scores (%) |
| medial orbitofrontal | L | 1.62 | 2.34 | 2.09 | 3.28 |
| | R | 1.46 | 2.67 | 1.34 | 3.13 |
| middle temporal | L | 1.54 | 0.97 | 2.24 | 1.79 |
| | R | 1.29 | 1.21 | 2.01 | 1.57 |
| parahippocampal | L | 1.62 | 1.94 | 2.31 | 1.87 |
| | R | 2.43 | 2.75 | 2.16 | 2.84 |
| paracentral | L | 0.97 | 1.54 | 1.57 | 1.94 |
| | R | 1.46 | 1.21 | 1.94 | 1.12 |
| pars opercularis | L | 4.77 | 0.81 | 3.81 | 1.19 |
| | R | 3.80 | 0.97 | 3.81 | 0.97 |
| pars orbitalis | L | 2.75 | 2.26 | 2.61 | 1.79 |
| | R | 2.18 | 1.54 | 1.64 | 2.76 |
| pars triangularis | L | 3.80 | 0.65 | 3.13 | 1.87 |
| | R | 3.88 | 0.81 | 3.28 | 1.12 |
| pericalcarine | L | 3.48 | 3.64 | 4.25 | 3.58 |
| | R | 3.23 | 3.07 | 2.99 | 5.15 |
| postcentral | L | 1.13 | 1.94 | 0.82 | 2.31 |
| | R | 0.49 | 1.13 | 1.12 | 1.57 |
| posterior cingulate | L | 5.01 | 1.70 | 5.67 | 2.01 |
| | R | 3.31 | 1.78 | 4.18 | 1.49 |
| precentral | L | 2.34 | 0.81 | 2.84 | 0.90 |
| | R | 2.02 | 0.32 | 2.76 | 0.52 |
| precuneus | L | 2.18 | 1.21 | 1.42 | 2.09 |
| | R | 1.46 | 0.81 | 2.31 | 1.72 |
| rostral anterior cingulate | L | 2.26 | 1.70 | 3.43 | 2.54 |
| | R | 1.86 | 1.86 | 2.91 | 2.54 |
| rostral middle frontal | L | 0.81 | 1.86 | 1.64 | 2.01 |
| | R | 1.05 | 2.67 | 1.27 | 2.39 |
| superior frontal | L | 4.53 | 0.89 | 3.36 | 1.57 |
| | R | 3.40 | 1.21 | 2.76 | 1.64 |
| superior parietal | L | 0.57 | 2.51 | 0.52 | 2.91 |
| | R | 0.57 | 1.62 | 0.82 | 2.61 |
| superior temporal | L | 3.15 | 0.49 | 3.13 | 0.60 |
| | R | 2.34 | 1.29 | 3.36 | 0.90 |
| supramarginal gyrus | L | 2.99 | 1.21 | 2.39 | 1.04 |
| | R | 2.43 | 1.13 | 4.10 | 1.49 |
| frontal pole | L | 2.83 | 2.99 | 2.46 | 3.88 |
| | R | 2.43 | 3.56 | 2.31 | 3.66 |
| temporal pole | L | 2.02 | 2.34 | 1.27 | 2.54 |
| | R | 2.26 | 1.29 | 3.06 | 1.94 |

| eTable 7. Percentage of individuals with infra- or supranormal normative regional z-scores based on group | | | | | |
|---|---|---|---|---|---|
| | | Normative z-scores in HI | | Normative z-scores in CHR-P | |
| Region | Hemi | infranormal z-scores (%) | supranormal z-scores (%) | infranormal z-scores (%) | supranormal z-scores (%) |
| transverse temporal | L | 2.83 | 1.37 | 2.61 | 1.57 |
| | R | 2.43 | 1.13 | 1.64 | 1.87 |
| insula | L | 2.99 | 1.21 | 4.48 | 0.60 |
| | R | 3.48 | 0.57 | 4.48 | 0.37 |
| **Surface Area** | | | | | |
| banks superior temporal sulcus | L | 1.54 | 2.75 | 2.39 | 1.79 |
| | R | 2.99 | 2.75 | 2.01 | 2.24 |
| caudal anterior cingulate | L | 0.65 | 4.28 | 0.30 | 3.21 |
| | R | 0.89 | 2.34 | 0.60 | 3.13 |
| caudal middle frontal | L | 1.46 | 2.26 | 1.64 | 2.39 |
| | R | 0.89 | 2.26 | 1.49 | 2.99 |
| cuneus | L | 1.54 | 2.75 | 2.46 | 2.69 |
| | R | 1.70 | 2.59 | 1.72 | 2.91 |
| entorhinal cortex | L | 2.51 | 6.95 | 2.61 | 5.52 |
| | R | 1.37 | 5.01 | 1.34 | 5.07 |
| fusiform gyrus | L | 1.70 | 2.02 | 2.91 | 2.24 |
| | R | 2.91 | 1.70 | 2.61 | 1.87 |
| inferior parietal | L | 2.02 | 1.94 | 3.13 | 1.87 |
| | R | 2.43 | 2.10 | 3.06 | 2.84 |
| inferior temporal | L | 3.64 | 2.59 | 2.31 | 2.76 |
| | R | 2.99 | 2.59 | 2.46 | 2.16 |
| isthmus cingulate | L | 1.05 | 3.23 | 0.75 | 3.06 |
| | R | 1.29 | 3.23 | 0.82 | 3.81 |
| lateral occipital | L | 1.94 | 2.99 | 2.09 | 3.06 |
| | R | 1.46 | 2.43 | 1.64 | 2.61 |
| lateral orbitofrontal | L | 2.67 | 1.21 | 2.46 | 2.24 |
| | R | 2.51 | 2.26 | 2.84 | 3.28 |
| lingual gyrus | L | 2.51 | 2.51 | 2.99 | 2.01 |
| | R | 2.75 | 2.59 | 2.84 | 3.06 |
| medial orbitofrontal | L | 1.70 | 3.96 | 1.79 | 3.21 |
| | R | 1.94 | 2.99 | 2.46 | 2.69 |
| middle temporal | L | 3.88 | 1.62 | 3.06 | 3.21 |
| | R | 2.91 | 2.18 | 2.54 | 2.69 |
| parahippocampal | L | 1.46 | 2.59 | 2.09 | 2.16 |
| | R | 2.67 | 2.75 | 3.58 | 3.13 |
| paracentral | L | 1.13 | 2.34 | 0.82 | 2.91 |
| | R | 1.54 | 2.83 | 0.82 | 3.06 |
| pars opercularis | L | 1.54 | 3.72 | 1.94 | 2.39 |
| | R | 1.05 | 3.80 | 0.97 | 3.28 |
| pars orbitalis | L | 2.51 | 1.70 | 2.39 | 2.84 |
| | R | 2.67 | 1.62 | 1.72 | 2.24 |

| eTable 7. Percentage of individuals with infra- or supranormal normative regional z-scores based on group | | | | | |
|---|---|---|---|---|---|
| | | Normative z-scores in HI | | Normative z-scores in CHR-P | |
| Region | Hemi | infranormal z-scores (%) | supranormal z-scores (%) | infranormal z-scores (%) | supranormal z-scores (%) |
| pars triangularis | L | 2.83 | 2.18 | 2.84 | 1.49 |
| | R | 2.91 | 1.70 | 1.72 | 1.94 |
| pericalcarine | L | 1.86 | 2.99 | 1.49 | 2.99 |
| | R | 2.26 | 2.51 | 2.24 | 2.01 |
| postcentral | L | 0.97 | 4.12 | 0.90 | 5.45 |
| | R | 1.21 | 4.28 | 1.79 | 3.43 |
| posterior cingulate | L | 1.62 | 2.51 | 1.42 | 2.91 |
| | R | 1.54 | 2.34 | 1.49 | 2.46 |
| precentral | L | 2.10 | 3.88 | 1.27 | 4.70 |
| | R | 1.21 | 3.23 | 0.45 | 3.96 |
| precuneus | L | 1.37 | 2.02 | 2.24 | 2.91 |
| | R | 1.54 | 1.21 | 1.94 | 2.91 |
| rostral anterior cingulate | L | 0.97 | 3.88 | 1.34 | 3.28 |
| | R | 0.73 | 2.02 | 0.60 | 2.24 |
| rostral middle frontal | L | 2.18 | 1.86 | 2.69 | 2.61 |
| | R | 1.54 | 2.10 | 2.31 | 3.13 |
| superior frontal | L | 2.99 | 3.07 | 3.66 | 3.28 |
| | R | 2.26 | 2.67 | 2.69 | 2.31 |
| superior parietal | L | 1.86 | 2.10 | 1.64 | 2.61 |
| | R | 1.94 | 1.62 | 2.24 | 3.06 |
| superior temporal | L | 2.10 | 2.91 | 2.54 | 3.43 |
| | R | 2.02 | 3.07 | 1.79 | 2.99 |
| supramarginal gyrus | L | 2.43 | 5.09 | 2.61 | 3.81 |
| | R | 1.62 | 2.26 | 1.79 | 1.87 |
| frontal pole | L | 1.29 | 1.78 | 1.79 | 1.12 |
| | R | 0.65 | 1.21 | 1.12 | 1.42 |
| temporal pole | L | 1.05 | 1.29 | 1.12 | 1.94 |
| | R | 1.29 | 2.99 | 0.90 | 2.99 |
| transverse temporal | L | 0.81 | 2.83 | 1.19 | 3.36 |
| | R | 0.73 | 1.94 | 0.75 | 2.31 |
| insula | L | 0.81 | 6.22 | 1.34 | 6.72 |
| | R | 1.54 | 6.31 | 1.12 | 7.01 |

[a] significant two-proportion z-tests difference in infranormal z-scores between CHR-P and HI at $P_{FDR} < 0.05$; [b] significant two-proportion z-tests difference in supranormal z-scores between CHR-P and HI at $P_{FDR} < 0.05$; CHR-P = clinical high-risk for psychosis; Hemi = hemisphere; HI = healthy individuals; L = left; R = right.

| eTable 8. Percentage of CHR-P with infra- or supranormal normative regional z-scores according to medication status. | | | | | |
|---|---|---|---|---|---|
| | | Unmediated CHR-P Individuals (N = 1061) | | Mediated CHR-P Individuals (N = 243) | |
| Region | Hemi | infranormal z-scores (%) | supranormal z-scores (%) | infranormal z-scores (%) | supranormal z-scores (%) |
| **Subcortical Volume** | | | | | |
| nucleus accumbens | L | 5.00 | 0.57 | 8.64 | 0.00 |
| | R | 6.03 | 0.00 | 11.11 | 0.41 |
| amygdala | L | 3.49 | 0.57 | 4.53 | 0.82 |
| | R | 3.30 | 2.17 | 5.76 | 1.65 |
| caudate | L | 5.56 | 0.57 | 7.82 | 0.82 |
| | R | 4.43 | 0.94 | 7.41 | 2.06 |
| hippocampus | L | 4.05 | 0.75 | 7.82 | 0.82 |
| | R | 3.49 | 0.66 | 5.76 | 0.82 |
| pallidum | L | 5.00 | 0.28 | 8.64 | 0.82 |
| | R | 5.18 | 1.13 | 9.47 | 0.41 |
| putamen | L | 9.61 | 0.47 | 11.93 | 0.00 |
| | R | 11.50 | 0.28 | 11.93 | 0.00 |
| thalamus | L | 7.35 | 0.66 | 11.52 | 1.65 |
| | R | 7.35 | 0.38 | 14.40 | 0.00 |
| **Cortical Thickness** | | | | | |
| banks superior temporal sulcus | L | 2.26 | 1.60 | 2.06 | 1.23 |
| | R | 3.49 | 2.07 | 2.88 | 2.88 |
| caudal anterior cingulate | L | 2.64 | 3.02 | 2.47 | 4.12 |
| | R | 3.86 | 4.52 | 2.88 | 4.53 |
| caudal middle frontal | L | 2.64 | 1.04 | 1.65 | 1.65 |
| | R | 2.83 | 1.79 | 2.47 | 1.23 |
| cuneus | L | 1.79 | 2.73 | 2.47 | 0.82 |
| | R | 1.98 | 2.92 | 1.65 | 3.70 |
| entorhinal cortex | L | 2.73 | 4.71 | 1.23 | 2.06 |
| | R | 2.73 | 4.05 | 2.88 | 2.47 |
| fusiform gyrus | L | 1.51 | 2.83 | 1.23 | 4.12 |
| | R | 1.98 | 2.73 | 1.65 | 4.94 |
| inferior parietal | L | 1.23 | 2.92 | 3.29 | 1.23 |
| | R | 1.89 | 2.17 | 2.88 | 0.82 |
| inferior temporal | L | 1.23 | 3.20 | 1.65 | 3.70 |
| | R | 1.70 | 5.66 | 3.29 | 1.65 |
| isthmus cingulate | L | 5.28 | 1.51 | 4.53 | 2.47 |
| | R | 4.62 | 1.51 | 5.35 | 1.23 |
| lateral occipital | L | 0.66 | 4.62 | 0.41 | 4.53 |
| | R | 0.85 | 4.52 | 1.23 | 5.76 |
| lateral orbitofrontal | L | 1.89 | 1.79 | 0.82 | 2.06 |
| | R | 1.79 | 2.17 | 1.23 | 2.06 |
| lingual gyrus | L | 3.02 | 1.89 | 5.76 | 3.29 |

| eTable 8. Percentage of CHR-P with infra- or supranormal normative regional z-scores according to medication status. | | | | | |
|---|---|---|---|---|---|
| | | Unmediated CHR-P Individuals (N = 1061) | | Mediated CHR-P Individuals (N = 243) | |
| Region | Hemi | infranormal z-scores (%) | supranormal z-scores (%) | infranormal z-scores (%) | supranormal z-scores (%) |
| | R | 2.17 | 2.17 | 1.65 | 2.88 |
| medial orbitofrontal | L | 1.98 | 3.30 | 2.47 | 3.29 |
| | R | 1.41 | 3.20 | 1.23 | 2.88 |
| middle temporal | L | 1.98 | 1.89 | 3.29 | 1.65 |
| | R | 1.79 | 1.70 | 2.88 | 1.23 |
| parahippocampal | L | 1.79 | 1.51 | 4.94 | 2.47 |
| | R | 1.98 | 2.92 | 3.29 | 2.88 |
| paracentral | L | 1.41 | 2.07 | 2.06 | 0.82 |
| | R | 1.89 | 1.04 | 1.65 | 0.41 |
| pars opercularis | L | 3.77 | 1.13 | 4.12 | 0.82 |
| | R | 3.86 | 1.13 | 4.12 | 0.41 |
| pars orbitalis | L | 2.36 | 1.89 | 3.70 | 0.82 |
| | R | 1.51 | 2.83 | 2.06 | 2.06 |
| pars triangularis | L | 3.30 | 1.98 | 2.47 | 1.23 |
| | R | 3.58 | 1.13 | 2.47 | 1.23 |
| pericalcarine | L | 4.34 | 3.68 | 3.70 | 3.29 |
| | R | 2.92 | 5.56 | 2.88 | 3.29 |
| postcentral | L | 0.94 | 2.64 | 0.41 | 1.23 |
| | R | 1.41 | 1.51 | 0.00 | 1.65 |
| posterior cingulate | L | 5.47 | 2.17 | 6.58 | 1.65 |
| | R | 4.24 | 1.51 | 3.29 | 1.65 |
| precentral | L | 2.83 | 0.85 | 2.88 | 0.82 |
| | R | 3.11 | 0.47 | 1.65 | 0.82 |
| precuneus | L | 1.32 | 1.98 | 1.65 | 1.65 |
| | R | 2.26 | 1.89 | 2.88 | 1.23 |
| rostral anterior cingulate | L | 3.49 | 2.26 | 3.29 | 3.70 |
| | R | 3.02 | 3.02 | 2.88 | 0.41 |
| rostral middle frontal | L | 1.70 | 1.89 | 1.23 | 2.06 |
| | R | 1.32 | 2.26 | 0.82 | 2.88 |
| superior frontal | L | 3.49 | 1.60 | 2.88 | 1.65 |
| | R | 3.30 | 1.60 | 0.82 | 1.65 |
| superior parietal | L | 0.57 | 2.83 | 0.41 | 2.88 |
| | R | 0.85 | 2.36 | 0.82 | 3.29 |
| superior temporal | L | 2.64 | 0.75 | 4.53 | 0.00 |
| | R | 3.30 | 1.04 | 3.29 | 0.41 |
| supramarginal gyrus | L | 2.45 | 0.85 | 2.47 | 2.06 |
| | R | 3.58 | 1.41 | 6.58 | 2.06 |
| frontal pole | L | 2.73 | 3.58 | 1.65 | 4.53 |
| | R | 2.73 | 3.77 | 0.82 | 3.70 |

| **eTable 8. Percentage of CHR-P with infra- or supranormal normative regional z-scores according to medication status.** | | | | | |
|---|---|---|---|---|---|
| | | **Unmediated CHR-P Individuals (N = 1061)** | | **Mediated CHR-P Individuals (N = 243)** | |
| **Region** | **Hemi** | **infranormal z-scores (%)** | **supranormal z-scores (%)** | **infranormal z-scores (%)** | **supranormal z-scores (%)** |
| temporal pole | L | 1.23 | 2.54 | 1.65 | 2.47 |
| | R | 3.30 | 1.89 | 2.06 | 1.65 |
| transverse temporal | L | 2.73 | 1.70 | 2.06 | 0.82 |
| | R | 1.70 | 2.07 | 1.65 | 1.23 |
| insula | L | 4.62 | 0.47 | 3.70 | 0.82 |
| | R | 4.15 | 0.19 | 5.35 | 1.23 |
| **Surface Area** | | | | | |
| banks superior temporal sulcus | L | 2.73 | 1.60 | 0.82 | 2.88 |
| | R | 2.17 | 2.07 | 1.23 | 2.88 |
| caudal anterior cingulate | L | 0.19 | 3.02 | 0.41 | 4.53 |
| | R | 0.66 | 3.11 | 0.00 | 3.70 |
| caudal middle frontal | L | 1.51 | 2.36 | 2.47 | 2.47 |
| | R | 1.60 | 2.92 | 0.82 | 2.88 |
| cuneus | L | 2.54 | 2.83 | 2.47 | 1.65 |
| | R | 1.60 | 3.11 | 2.06 | 2.06 |
| entorhinal cortex | L | 2.92 | 5.47 | 1.23 | 6.17 |
| | R | 1.32 | 5.09 | 1.65 | 4.12 |
| fusiform gyrus | L | 2.83 | 2.36 | 3.29 | 2.06 |
| | R | 2.36 | 1.79 | 3.70 | 2.06 |
| inferior parietal | L | 3.30 | 1.89 | 2.88 | 2.06 |
| | R | 3.02 | 2.83 | 3.29 | 3.29 |
| inferior temporal | L | 2.17 | 2.45 | 2.88 | 3.70 |
| | R | 2.07 | 2.07 | 3.29 | 2.88 |
| isthmus cingulate | L | 0.75 | 2.54 | 0.82 | 5.76 |
| | R | 0.57 | 3.20 | 2.06 | 6.58 |
| lateral occipital | L[b] | 2.26 | 2.17 | 0.82 | 7.00 |
| | R | 1.23 | 2.26 | 3.29 | 2.88 |
| lateral orbitofrontal | L | 2.83 | 2.07 | 0.82 | 2.47 |
| | R | 2.73 | 3.11 | 3.29 | 3.70 |
| lingual gyrus | L | 2.92 | 1.98 | 3.70 | 2.06 |
| | R | 2.54 | 3.11 | 4.53 | 2.47 |
| medial orbitofrontal | L | 1.79 | 3.58 | 2.06 | 1.23 |
| | R | 2.36 | 2.26 | 3.29 | 4.53 |
| middle temporal | L | 3.20 | 2.83 | 2.47 | 4.53 |
| | R | 2.83 | 2.73 | 1.65 | 2.47 |
| parahippocampal | L | 2.07 | 1.89 | 2.06 | 3.29 |
| | R | 3.68 | 3.11 | 3.70 | 3.29 |
| paracentral | L | 0.85 | 2.73 | 0.82 | 3.70 |
| | R | 0.94 | 3.20 | 0.41 | 2.47 |
| pars opercularis | L | 1.70 | 2.07 | 3.29 | 3.70 |

| **eTable 8. Percentage of CHR-P with infra- or supranormal normative regional z-scores according to medication status.** | | | | | |
|---|---|---|---|---|---|
| | | **Unmediated CHR-P Individuals (N = 1061)** | | **Mediated CHR-P Individuals (N = 243)** | |
| **Region** | **Hemi** | **infranormal z-scores (%)** | **supranormal z-scores (%)** | **infranormal z-scores (%)** | **supranormal z-scores (%)** |
| | R | 0.85 | 3.30 | 1.65 | 3.29 |
| pars orbitalis | L | 2.36 | 2.73 | 2.06 | 3.29 |
| | R | 1.60 | 2.54 | 2.06 | 1.23 |
| pars triangularis | L | 3.02 | 1.51 | 1.65 | 1.65 |
| | R | 1.70 | 2.07 | 1.65 | 1.23 |
| pericalcarine | L | 1.32 | 2.83 | 2.06 | 3.29 |
| | R | 2.26 | 2.07 | 1.65 | 2.06 |
| postcentral | L | 0.75 | 5.37 | 1.65 | 5.76 |
| | R | 1.98 | 3.30 | 1.23 | 3.70 |
| posterior cingulate | L | 1.41 | 2.92 | 1.65 | 2.47 |
| | R | 1.41 | 2.45 | 2.06 | 2.88 |
| precentral | L | 0.94 | 4.43 | 2.47 | 5.76 |
| | R | 0.47 | 3.96 | 0.41 | 4.12 |
| precuneus | L | 2.26 | 2.73 | 2.06 | 3.29 |
| | R | 2.07 | 2.73 | 1.65 | 2.88 |
| rostral anterior cingulate | L | 1.13 | 3.30 | 2.47 | 3.70 |
| | R | 0.75 | 2.54 | 0.00 | 1.23 |
| rostral middle frontal | L | 2.92 | 2.73 | 1.65 | 2.47 |
| | R | 2.26 | 2.73 | 2.47 | 4.94 |
| superior frontal | L | 3.96 | 3.49 | 2.47 | 2.06 |
| | R | 2.54 | 2.17 | 3.29 | 2.88 |
| superior parietal | L | 1.60 | 2.64 | 2.06 | 2.47 |
| | R | 1.98 | 3.02 | 3.29 | 2.88 |
| superior temporal | L | 2.36 | 3.49 | 3.29 | 3.70 |
| | R | 1.98 | 2.64 | 1.23 | 4.12 |
| supramarginal gyrus | L | 2.83 | 3.86 | 1.65 | 4.12 |
| | R | 1.60 | 2.17 | 2.88 | 0.41 |
| frontal pole | L | 1.70 | 1.04 | 2.06 | 1.65 |
| | R | 1.23 | 1.32 | 0.41 | 2.06 |
| temporal pole | L | 0.75 | 2.17 | 1.65 | 0.41 |
| | R | 0.94 | 3.11 | 0.82 | 2.88 |
| transverse temporal | L | 1.32 | 3.39 | 0.82 | 3.29 |
| | R | 0.75 | 2.73 | 0.82 | 0.41 |
| insula | L | 1.32 | 6.03 | 1.65 | 9.88 |
| | R | 0.94 | 6.88 | 2.06 | 7.82 |

[a] significant two-proportion z-tests difference in infranormal z-scores between CHR-P with and without antipsychotic medication exposure at $P_{FDR} < 0.05$; [b] significant two-proportion z-tests difference in supranormal z-scores between CHR-P with and without antipsychotic medication exposure at $P_{FDR} < 0.05$; CHR-P = clinical high-risk for psychosis; Hemi = hemisphere; L = left; R = right.

| eTable 9. Percentage of individuals with infra- or supranormal normative average deviation scores based on group | | | | |
|---|---|---|---|---|
| | Normative z-scores in HI (N = 1237) | | Normative z-scores in CHR-P (N = 1340) | |
| Region | infranormal z-scores (%) | supranormal z-scores (%) | infranormal z-scores (%) | supranormal z-scores (%) |
| **Average Deviation Score (ADS)** | | | | |
| $ADS_G$[a] | 1.70 | 2.18 | 4.18 | 2.01 |
| $ADS_{SV}$[a] | 1.62 | 2.59 | 3.13 | 2.69 |
| $ADS_{CT}$ | 1.94 | 2.67 | 3.21 | 2.31 |
| $ADS_{SA}$[a] | 1.05 | 2.83 | 3.51 | 2.69 |

[a] significant two-proportion z-tests difference in infranormal ADS between CHR-P and HI at $P_{FDR} < 0.05$; [b] significant two-proportion z-tests difference in supranormal ADS between CHR-P and HI at $P_{FDR} < 0.05$; $ASD_{CT}$=average deviation score-cortical thickness; $ADS_G$=Average deviation score-global; $ADS_{SA}$=average deviation score-surface area; $ADS_{SV}$=average deviation score-subcortical volume; CHR-P = clinical high-risk for psychosis; Hemi = hemisphere; HI = healthy individuals; L = left; R = right.

| eTable 10. Percentage of CHR-P with infra- or supranormal normative average deviation scores according to medication status | | | | |
|---|---|---|---|---|
| | Unmediated CHR-P Individuals (N = 1061) | | Mediated CHR-P Individuals (N = 243) | |
| Region | infranormal z-scores (%) | supranormal z-scores (%) | infranormal z-scores (%) | supranormal z-scores (%) |
| **Average Deviation Score (ADS)** | | | | |
| $ADS_G$ | 2.92 | 1.60 | 5.35 | 2.06 |
| $ADS_{SV}$ | 1.98 | 2.07 | 4.11 | 2.88 |
| $ADS_{CT}$ | 2.83 | 2.17 | 2.47 | 2.06 |
| $ADS_{SA}$ | 2.83 | 2.36 | 2.47 | 2.88 |

[a] significant two-proportion z-tests difference in infranormal ADS between CHR-P with and without antipsychotic medication exposure at $P_{FDR}$ < 0.05; [b] significant two-proportion z-tests difference in supranormal ADS between CHR-P with and without antipsychotic medication exposure at $P_{FDR}$ < 0.05; $ASD_{CT}$=average deviation score-cortical thickness; $ADS_G$=Average deviation score-global; $ADS_{SA}$=average deviation score-surface area; $ADS_{SV}$=average deviation score-subcortical volume; CHR-P = clinical high-risk for psychosis; Hemi = hemisphere; L = left; R = right.

| eTable 11. Associations between regional normative z-scores and positive symptoms and IQ in CHR-P | | | | | |
|---|---|---|---|---|---|
| | | Positive Symptoms | | IQ | |
| Region | Hemi | β Estimate | Uncorrected P values | β Estimate | Uncorrected P values |
| **Subcortical Volume** | | | | | |
| nucleus accumbens | L | 1.53E-03 | 0.96 | 0.02 | 0.64 |
| | R | -0.02 | 0.46 | 0.01 | 0.73 |
| amygdala | L | -0.01 | 0.73 | 0.03 | 0.32 |
| | R | 1.81E-03 | 0.95 | 0.02 | 0.55 |
| caudate | L | -0.02 | 0.53 | 0.11 | 6.00E-04[a] |
| | R | -0.03 | 0.24 | 0.10 | 2.60E-03 |
| hippocampus | L | -0.04 | 0.20 | -0.01 | 0.73 |
| | R | -0.03 | 0.21 | -0.01 | 0.79 |
| pallidum | L | -0.01 | 0.66 | 0.03 | 0.43 |
| | R | -0.02 | 0.49 | -0.01 | 0.79 |
| putamen | L | 0.01 | 0.78 | -0.02 | 0.58 |
| | R | -0.01 | 0.61 | 1.92E-03 | 0.95 |
| thalamus | L | -0.03 | 0.32 | 0.07 | 0.02 |
| | R | -0.02 | 0.38 | 0.06 | 0.07 |
| **Cortical Thickness** | | | | | |
| banks superior temporal sulcus | L | 0.01 | 0.82 | 0.01 | 0.85 |
| | R | 0.01 | 0.73 | -0.03 | 0.40 |
| caudal anterior cingulate | L | -0.02 | 0.57 | -0.01 | 0.68 |
| | R | -0.01 | 0.73 | 0.03 | 0.36 |
| caudal middle frontal | L | -0.04 | 0.19 | 0.05 | 0.15 |
| | R | 0.01 | 0.64 | 0.05 | 0.15 |
| cuneus | L | -0.03 | 0.28 | 0.06 | 0.06 |
| | R | 0.01 | 0.69 | 0.04 | 0.25 |
| entorhinal cortex | L | -0.02 | 0.49 | 0.05 | 0.14 |
| | R | 0.03 | 0.22 | 0.04 | 0.26 |
| fusiform gyrus | L | -0.04 | 0.16 | 0.03 | 0.30 |
| | R | -0.02 | 0.46 | -3.01E-03 | 0.93 |
| inferior parietal | L | 0.01 | 0.63 | -0.06 | 0.07 |
| | R | 0.01 | 0.82 | -0.07 | 0.04 |
| inferior temporal | L | -0.02 | 0.58 | -3.98E-03 | 0.90 |
| | R | -0.03 | 0.25 | 0.03 | 0.37 |
| isthmus cingulate | L | -5.49E-05 | 1.00 | -0.02 | 0.51 |
| | R | -0.03 | 0.23 | 0.03 | 0.37 |
| lateral occipital | L | -0.07 | 0.02 | -0.06 | 0.10 |
| | R | -0.04 | 0.11 | -0.02 | 0.47 |
| lateral orbitofrontal | L | -0.03 | 0.30 | -0.02 | 0.53 |
| | R | -3.76E-03 | 0.89 | 0.03 | 0.35 |
| lingual gyrus | L | 0.02 | 0.52 | 0.08 | 0.02 |
| | R | -0.01 | 0.74 | 0.05 | 0.13 |

| eTable 11. Associations between regional normative z-scores and positive symptoms and IQ in CHR-P | | | | | |
|---|---|---|---|---|---|
| | | **Positive Symptoms** | | **IQ** | |
| **Region** | **Hemi** | **β Estimate** | **Uncorrected P values** | **β Estimate** | **Uncorrected P values** |
| medial orbitofrontal | L | 1.69E-03 | 0.95 | -0.04 | 0.20 |
| | R | 0.02 | 0.45 | -0.02 | 0.57 |
| middle temporal | L | -3.07E-03 | 0.91 | -0.01 | 0.73 |
| | R | 0.02 | 0.43 | -0.04 | 0.18 |
| parahippocampal | L | 0.01 | 0.71 | 0.04 | 0.19 |
| | R | -1.02E-03 | 0.97 | -0.01 | 0.80 |
| paracentral | L | -0.01 | 0.79 | -0.01 | 0.86 |
| | R | 0.04 | 0.11 | -0.05 | 0.10 |
| pars opercularis | L | 0.02 | 0.46 | 3.03E-03 | 0.93 |
| | R | -0.01 | 0.80 | 0.08 | 0.02 |
| pars orbitalis | L | -0.01 | 0.73 | -0.02 | 0.50 |
| | R | -0.03 | 0.30 | -0.03 | 0.30 |
| pars triangularis | L | 0.02 | 0.48 | 0.02 | 0.59 |
| | R | 0.04 | 0.17 | -0.03 | 0.42 |
| pericalcarine | L | 0.01 | 0.67 | 0.03 | 0.33 |
| | R | 0.01 | 0.81 | 0.02 | 0.50 |
| postcentral | L | -0.02 | 0.47 | 0.05 | 0.18 |
| | R | 0.06 | 0.04 | 0.05 | 0.14 |
| posterior cingulate | L | 0.04 | 0.12 | -0.01 | 0.84 |
| | R | -0.02 | 0.54 | -0.02 | 0.64 |
| precentral | L | -0.01 | 0.60 | 0.05 | 0.11 |
| | R | 0.01 | 0.74 | 0.02 | 0.64 |
| precuneus | L | 0.02 | 0.38 | 0.06 | 0.09 |
| | R | 0.02 | 0.37 | -0.03 | 0.29 |
| rostral anterior cingulate | L | 0.02 | 0.57 | -0.01 | 0.68 |
| | R | 3.15E-03 | 0.91 | -0.03 | 0.43 |
| rostral middle frontal | L | -0.03 | 0.25 | -0.02 | 0.60 |
| | R | -0.01 | 0.67 | -0.05 | 0.10 |
| superior frontal | L | -0.01 | 0.79 | -0.02 | 0.46 |
| | R | -0.01 | 0.79 | -0.01 | 0.67 |
| superior parietal | L | -0.01 | 0.83 | -0.08 | 0.02 |
| | R | 0.05 | 0.09 | -0.03 | 0.41 |
| superior temporal | L | -0.02 | 0.43 | 0.03 | 0.37 |
| | R | 0.03 | 0.34 | 0.04 | 0.25 |
| supramarginal gyrus | L | 0.04 | 0.13 | 0.03 | 0.42 |
| | R | 0.06 | 0.03 | 0.01 | 0.67 |
| frontal pole | L | 0.01 | 0.69 | -0.03 | 0.42 |
| | R | -0.02 | 0.45 | 0.04 | 0.27 |
| temporal pole | L | 0.03 | 0.36 | 2.99E-03 | 0.93 |
| | R | 0.02 | 0.42 | 0.04 | 0.25 |

| eTable 11. Associations between regional normative z-scores and positive symptoms and IQ in CHR-P | | | | | |
|---|---|---|---|---|---|
| | | **Positive Symptoms** | | **IQ** | |
| **Region** | **Hemi** | **β Estimate** | **Uncorrected P values** | **β Estimate** | **Uncorrected P values** |
| transverse temporal | L | -0.01 | 0.83 | -0.03 | 0.39 |
| | R | 0.03 | 0.23 | 0.01 | 0.84 |
| insula | L | -3.09E-04 | 0.99 | -0.02 | 0.57 |
| | R | -2.04E-03 | 0.94 | 0.07 | 0.03 |
| **Surface Area** | | | | | |
| banks superior temporal | L | 0.02 | 0.53 | -0.01 | 0.83 |
| sulcus | R | 0.02 | 0.54 | -0.07 | 0.05 |
| caudal anterior cingulate | L | -0.01 | 0.59 | 0.03 | 0.39 |
| | R | -0.01 | 0.66 | 0.05 | 0.17 |
| caudal middle frontal | L | -0.06 | 0.04 | 0.04 | 0.28 |
| | R | -3.80E-03 | 0.89 | 0.01 | 0.80 |
| cuneus | L | -2.72E-03 | 0.92 | 0.11 | 5.64E-04[a] |
| | R | 0.01 | 0.76 | 0.02 | 0.53 |
| entorhinal cortex | L | 1.22E-03 | 0.96 | 0.02 | 0.63 |
| | R | -0.01 | 0.80 | 4.72E-03 | 0.89 |
| fusiform gyrus | L | 0.03 | 0.24 | 0.09 | 0.01 |
| | R | 0.03 | 0.27 | 0.05 | 0.15 |
| inferior parietal | L | 0.05 | 0.08 | -0.03 | 0.30 |
| | R | -0.01 | 0.80 | -0.01 | 0.83 |
| inferior temporal | L | 0.02 | 0.53 | 0.04 | 0.23 |
| | R | -3.74E-03 | 0.89 | 0.07 | 0.03 |
| isthmus cingulate | L | -3.82E-03 | 0.89 | -0.04 | 0.24 |
| | R | 0.03 | 0.22 | -0.02 | 0.56 |
| lateral occipital | L | 0.01 | 0.66 | 0.01 | 0.66 |
| | R | -1.66E-03 | 0.95 | 0.06 | 0.06 |
| lateral orbitofrontal | L | -6.00E-04 | 0.98 | 4.52E-03 | 0.89 |
| | R | 0.01 | 0.67 | 0.02 | 0.58 |
| lingual gyrus | L | -0.01 | 0.84 | 0.07 | 0.03 |
| | R | -0.02 | 0.46 | 0.04 | 0.18 |
| medial orbitofrontal | L | -0.02 | 0.51 | 0.04 | 0.22 |
| | R | -0.04 | 0.13 | 0.04 | 0.21 |
| middle temporal | L | 0.02 | 0.45 | -0.01 | 0.79 |
| | R | 0.03 | 0.31 | 0.02 | 0.50 |
| parahippocampal | L | -0.03 | 0.35 | 0.02 | 0.53 |
| | R | -0.05 | 0.06 | 4.24E-03 | 0.90 |
| paracentral | L | 0.03 | 0.30 | -0.06 | 0.06 |
| | R | -0.05 | 0.08 | -0.08 | 0.02 |
| pars opercularis | L | -0.01 | 0.78 | 0.03 | 0.37 |
| | R | -2.56E-03 | 0.93 | 0.04 | 0.25 |
| pars orbitalis | L | -0.06 | 0.04 | 0.05 | 0.11 |

| eTable 11. Associations between regional normative z-scores and positive symptoms and IQ in CHR-P | | | | | |
|---|---|---|---|---|---|
| | | Positive Symptoms | | IQ | |
| Region | Hemi | β Estimate | Uncorrected P values | β Estimate | Uncorrected P values |
| | R | -0.05 | 0.10 | 0.06 | 0.06 |
| pars triangularis | L | -0.02 | 0.44 | 0.03 | 0.36 |
| | R | -0.02 | 0.46 | 0.01 | 0.67 |
| pericalcarine | L | 0.01 | 0.81 | 0.08 | 0.02 |
| | R | 3.51E-03 | 0.90 | 0.02 | 0.61 |
| postcentral | L | -0.03 | 0.28 | 0.01 | 0.70 |
| | R | -0.03 | 0.25 | -0.02 | 0.61 |
| posterior cingulate | L | -2.80E-03 | 0.92 | -0.01 | 0.68 |
| | R | 0.01 | 0.76 | 0.01 | 0.70 |
| precentral | L | -0.04 | 0.16 | 0.02 | 0.61 |
| | R | -0.01 | 0.60 | -1.18E-03 | 0.97 |
| precuneus | L | 0.01 | 0.75 | -0.09 | 0.00 |
| | R | -0.02 | 0.44 | 0.02 | 0.63 |
| rostral anterior cingulate | L | -0.04 | 0.20 | 0.09 | 0.01 |
| | R | -0.04 | 0.13 | 8.64E-04 | 0.98 |
| rostral middle frontal | L | 0.02 | 0.52 | 0.04 | 0.29 |
| | R | 0.02 | 0.39 | 0.03 | 0.44 |
| superior frontal | L | 0.01 | 0.78 | -3.30E-04 | 0.99 |
| | R | -0.03 | 0.27 | -0.01 | 0.84 |
| superior parietal | L | -0.02 | 0.48 | -0.02 | 0.50 |
| | R | -0.01 | 0.84 | -0.04 | 0.19 |
| superior temporal | L | -0.03 | 0.31 | -0.03 | 0.30 |
| | R | 0.03 | 0.29 | -0.04 | 0.27 |
| supramarginal gyrus | L | -0.03 | 0.21 | -0.03 | 0.36 |
| | R | -0.03 | 0.21 | -0.03 | 0.45 |
| frontal pole | L | -0.03 | 0.33 | -0.01 | 0.81 |
| | R | 0.01 | 0.82 | -0.01 | 0.74 |
| temporal pole | L | 0.02 | 0.43 | -0.01 | 0.67 |
| | R | 4.22E-03 | 0.88 | -0.06 | 0.09 |
| transverse temporal | L | -0.03 | 0.22 | -0.03 | 0.41 |
| | R | -0.01 | 0.78 | -0.04 | 0.26 |
| insula | L | -0.02 | 0.55 | -0.01 | 0.69 |
| | R | -0.05 | 0.08 | -0.04 | 0.23 |
| [a] significant at $P_{FDR}<0.05$; hemi = hemisphere; CHR-P = clinical high-risk for psychosis; IQ = intelligence quotient; L = left; R = right | | | | | |

| eTable 12. Associations between observed brain morphometric measures with positive symptoms and IQ in CHR-P | | | | | |
|---|---|---|---|---|---|
| | | Positive Symptoms Score | | IQ | |
| **Region** | **Hemi** | **β Estimate** | **Uncorrected P values** | **β Estimate** | **Uncorrected P values** |
| **Subcortical Volume** | | | | | |
| nucleus accumbens | L | -4.90E-03 | 0.85 | 0.01 | 0.83 |
| | R | -0.02 | 0.46 | 0.01 | 0.74 |
| amygdala | L | -0.01 | 0.76 | 0.02 | 0.39 |
| | R | -0.01 | 0.80 | 0.01 | 0.67 |
| caudate | L | -0.01 | 0.55 | 0.09 | 1.24E-03 |
| | R | -0.03 | 0.19 | 0.08 | 3.42E-03 |
| hippocampus | L | -0.04 | 0.06 | -0.02 | 0.44 |
| | R | -0.04 | 0.08 | -0.01 | 0.61 |
| pallidum | L | 3.14E-03 | 0.90 | 0.03 | 0.37 |
| | R | -0.02 | 0.45 | -0.01 | 0.83 |
| putamen | L | -5.93E-04 | 0.98 | -0.02 | 0.53 |
| | R | -0.02 | 0.40 | -0.01 | 0.75 |
| thalamus | L | -0.03 | 0.10 | 0.05 | 0.07 |
| | R | -0.03 | 0.19 | 0.03 | 0.17 |
| **Cortical Thickness** | | | | | |
| banks superior temporal sulcus | L | 0.01 | 0.68 | -4.95E-03 | 0.86 |
| | R | 0.01 | 0.58 | -0.03 | 0.33 |
| caudal anterior cingulate | L | -0.01 | 0.81 | -0.03 | 0.41 |
| | R | -0.01 | 0.65 | 0.03 | 0.39 |
| caudal middle frontal | L | -0.04 | 0.04 | 0.02 | 0.30 |
| | R | -0.01 | 0.77 | 0.03 | 0.23 |
| cuneus | L | -0.03 | 0.25 | 0.06 | 0.04 |
| | R | 0.01 | 0.82 | 0.04 | 0.17 |
| entorhinal cortex | L | -0.01 | 0.64 | 0.06 | 0.08 |
| | R | 0.05 | 0.07 | 0.03 | 0.34 |
| fusiform gyrus | L | -0.02 | 0.25 | 0.03 | 0.25 |
| | R | -0.01 | 0.59 | 0.01 | 0.75 |
| inferior parietal | L | 8.91E-04 | 0.96 | -0.04 | 0.04 |
| | R | 4.69E-03 | 0.78 | -0.05 | 0.01 |
| inferior temporal | L | -0.01 | 0.57 | -0.01 | 0.72 |
| | R | -0.02 | 0.34 | 0.02 | 0.53 |
| isthmus cingulate | L | 0.01 | 0.79 | -0.02 | 0.48 |
| | R | -0.03 | 0.25 | 0.02 | 0.50 |
| lateral occipital | L | -0.05 | 0.01 | -0.03 | 0.21 |
| | R | -0.04 | 0.06 | -0.01 | 0.58 |
| lateral orbitofrontal | L | -0.02 | 0.41 | -0.02 | 0.53 |
| | R | -4.99E-03 | 0.82 | 0.03 | 0.34 |
| lingual gyrus | L | 0.01 | 0.56 | 0.08 | 0.01 |
| | R | -0.01 | 0.60 | 0.05 | 0.06 |

| eTable 12. Associations between observed brain morphometric measures with positive symptoms and IQ in CHR-P | | | | | |
|---|---|---|---|---|---|
| | | **Positive Symptoms Score** | | **IQ** | |
| **Region** | **Hemi** | **β Estimate** | **Uncorrected P values** | **β Estimate** | **Uncorrected P values** |
| medial orbitofrontal | L | -3.64E-03 | 0.88 | -0.03 | 0.26 |
| | R | 0.01 | 0.82 | -0.02 | 0.57 |
| middle temporal | L | 0.01 | 0.74 | -0.01 | 0.64 |
| | R | 0.02 | 0.29 | -0.04 | 0.10 |
| parahippocampal | L | 0.02 | 0.48 | 0.04 | 0.23 |
| | R | 0.01 | 0.80 | -0.01 | 0.84 |
| paracentral | L | -0.01 | 0.64 | -0.01 | 0.71 |
| | R | 0.03 | 0.15 | -0.04 | 0.09 |
| pars opercularis | L | 0.01 | 0.74 | 4.25E-04 | 0.99 |
| | R | -0.01 | 0.79 | 0.05 | 0.05 |
| pars orbitalis | L | -0.01 | 0.65 | -0.02 | 0.56 |
| | R | -0.02 | 0.38 | -0.04 | 0.17 |
| pars triangularis | L | 0.02 | 0.43 | 0.01 | 0.78 |
| | R | 0.03 | 0.15 | -0.02 | 0.33 |
| pericalcarine | L | 0.01 | 0.64 | 0.03 | 0.29 |
| | R | 4.48E-03 | 0.87 | 0.02 | 0.46 |
| postcentral | L | -0.02 | 0.44 | 0.03 | 0.29 |
| | R | 0.05 | 0.03 | 0.03 | 0.18 |
| posterior cingulate | L | 0.04 | 0.07 | -0.02 | 0.54 |
| | R | -0.02 | 0.51 | -0.03 | 0.32 |
| precentral | L | -0.01 | 0.53 | 0.03 | 0.15 |
| | R | 1.98E-03 | 0.92 | 0.01 | 0.70 |
| precuneus | L | 0.01 | 0.40 | 0.03 | 0.19 |
| | R | 0.01 | 0.46 | -0.03 | 0.20 |
| rostral anterior cingulate | L | 0.01 | 0.58 | -0.02 | 0.54 |
| | R | 1.13E-03 | 0.96 | -0.02 | 0.60 |
| rostral middle frontal | L | -0.03 | 0.08 | -0.01 | 0.66 |
| | R | -0.01 | 0.56 | -0.04 | 0.07 |
| superior frontal | L | -0.01 | 0.45 | -0.03 | 0.16 |
| | R | -0.01 | 0.42 | -0.02 | 0.29 |
| superior parietal | L | -0.01 | 0.74 | -0.05 | 0.01 |
| | R | 0.03 | 0.09 | -0.02 | 0.26 |
| superior temporal | L | -0.02 | 0.29 | 0.02 | 0.47 |
| | R | 0.02 | 0.35 | 0.02 | 0.44 |
| supramarginal gyrus | L | 0.02 | 0.24 | 0.01 | 0.61 |
| | R | 0.04 | 0.02 | 0.01 | 0.79 |
| frontal pole | L | 0.01 | 0.76 | -0.03 | 0.27 |
| | R | -0.03 | 0.28 | 0.03 | 0.34 |
| temporal pole | L | 0.03 | 0.30 | 4.79E-03 | 0.88 |
| | R | 0.03 | 0.26 | 0.04 | 0.18 |

| **eTable 12. Associations between observed brain morphometric measures with positive symptoms and IQ in CHR-P** | | | | | |
|---|---|---|---|---|---|
| | | **Positive Symptoms Score** | | **IQ** | |
| **Region** | **Hemi** | **β Estimate** | **Uncorrected P values** | **β Estimate** | **Uncorrected P values** |
| transverse temporal | L | 1.23E-04 | 1.00 | -0.02 | 0.42 |
| | R | 0.03 | 0.27 | 4.82E-03 | 0.87 |
| insula | L | -0.01 | 0.59 | -0.01 | 0.84 |
| | R | -0.01 | 0.64 | 0.07 | 0.01 |
| **Surface Area** | | | | | |
| banks superior temporal sulcus | L | 0.01 | 0.71 | -2.35E-03 | 0.94 |
| | R | 0.01 | 0.56 | -0.06 | 0.03 |
| caudal anterior cingulate | L | -0.01 | 0.62 | 0.01 | 0.67 |
| | R | -0.01 | 0.72 | 0.02 | 0.49 |
| caudal middle frontal | L | -0.04 | 0.05 | 0.02 | 0.44 |
| | R | -1.95E-05 | 1.00 | 0.02 | 0.54 |
| cuneus | L | 2.41E-03 | 0.92 | 0.09 | 9.12E-04 |
| | R | 0.01 | 0.60 | 0.02 | 0.58 |
| entorhinal cortex | L | 0.01 | 0.72 | 0.02 | 0.62 |
| | R | 0.01 | 0.65 | -3.46E-03 | 0.91 |
| fusiform gyrus | L | 0.03 | 0.11 | 0.05 | 0.03 |
| | R | 0.03 | 0.12 | 0.02 | 0.43 |
| inferior parietal | L | 0.05 | 0.03 | -0.02 | 0.44 |
| | R | -0.01 | 0.62 | -0.02 | 0.48 |
| inferior temporal | L | 0.02 | 0.40 | 0.01 | 0.64 |
| | R | 0.01 | 0.58 | 0.03 | 0.14 |
| isthmus cingulate | L | -0.01 | 0.67 | -0.02 | 0.36 |
| | R | 0.02 | 0.31 | -0.02 | 0.55 |
| lateral occipital | L | 0.01 | 0.50 | -2.82E-03 | 0.91 |
| | R | 0.01 | 0.74 | 0.03 | 0.19 |
| lateral orbitofrontal | L | 0.01 | 0.77 | -3.43E-03 | 0.87 |
| | R | 4.74E-03 | 0.80 | 2.60E-03 | 0.91 |
| lingual gyrus | L | 2.16E-04 | 0.99 | 0.05 | 0.08 |
| | R | -0.01 | 0.73 | 0.04 | 0.14 |
| medial orbitofrontal | L | -0.01 | 0.72 | 0.02 | 0.47 |
| | R | -0.02 | 0.36 | 0.01 | 0.58 |
| middle temporal | L | 0.01 | 0.45 | -0.02 | 0.48 |
| | R | 0.02 | 0.16 | 0.01 | 0.73 |
| parahippocampal | L | -0.02 | 0.42 | 0.01 | 0.72 |
| | R | -0.04 | 0.14 | -0.01 | 0.79 |
| paracentral | L | 0.03 | 0.24 | -0.05 | 0.09 |
| | R | -0.04 | 0.09 | -0.06 | 0.02 |
| pars opercularis | L | -3.97E-03 | 0.86 | 0.01 | 0.71 |
| | R | -2.81E-03 | 0.91 | 0.04 | 0.14 |
| pars orbitalis | L | -0.04 | 0.06 | 0.04 | 0.14 |

| eTable 12. Associations between observed brain morphometric measures with positive symptoms and IQ in CHR-P | | | | | |
|---|---|---|---|---|---|
| | | Positive Symptoms Score | | IQ | |
| Region | Hemi | β Estimate | Uncorrected P values | β Estimate | Uncorrected P values |
| | R | -0.03 | 0.16 | 0.05 | 0.08 |
| pars triangularis | L | -0.01 | 0.61 | 0.03 | 0.28 |
| | R | -0.02 | 0.34 | 0.02 | 0.50 |
| pericalcarine | L | 0.01 | 0.76 | 0.07 | 0.02 |
| | R | 2.92E-04 | 0.99 | 0.02 | 0.54 |
| postcentral | L | -0.01 | 0.41 | 1.78E-03 | 0.94 |
| | R | -0.01 | 0.62 | -0.02 | 0.46 |
| posterior cingulate | L | -3.34E-03 | 0.88 | -0.02 | 0.51 |
| | R | 0.01 | 0.63 | -0.01 | 0.81 |
| precentral | L | -0.02 | 0.27 | 0.01 | 0.80 |
| | R | -0.01 | 0.76 | 0.01 | 0.82 |
| precuneus | L | 0.01 | 0.76 | -0.06 | 3.67E-03 |
| | R | -0.01 | 0.54 | 0.01 | 0.81 |
| rostral anterior cingulate | L | -0.02 | 0.29 | 0.06 | 0.01 |
| | R | -0.03 | 0.15 | -0.01 | 0.63 |
| rostral middle frontal | L | 0.01 | 0.59 | 0.01 | 0.48 |
| | R | 0.01 | 0.65 | 0.01 | 0.81 |
| superior frontal | L | 0.01 | 0.74 | 5.10E-04 | 0.98 |
| | R | -0.01 | 0.38 | -4.34E-03 | 0.83 |
| superior parietal | L | -0.01 | 0.59 | -0.01 | 0.63 |
| | R | -4.29E-03 | 0.83 | -0.03 | 0.17 |
| superior temporal | L | -0.01 | 0.49 | -0.03 | 0.26 |
| | R | 0.02 | 0.19 | -0.03 | 0.20 |
| supramarginal gyrus | L | -0.02 | 0.32 | -0.03 | 0.23 |
| | R | -0.02 | 0.22 | -0.01 | 0.74 |
| frontal pole | L | -0.01 | 0.77 | 0.01 | 0.74 |
| | R | 0.01 | 0.73 | -5.70E-05 | 1.00 |
| temporal pole | L | 0.02 | 0.48 | -0.01 | 0.65 |
| | R | 0.02 | 0.45 | -0.06 | 0.05 |
| transverse temporal | L | -0.02 | 0.39 | -0.03 | 0.35 |
| | R | 0.01 | 0.80 | -0.03 | 0.33 |
| insula | L | 0.01 | 0.77 | -0.01 | 0.54 |
| | R | -0.02 | 0.29 | -0.03 | 0.22 |

[a] significant at $P_{FDR}<0.05$; Hemi = hemisphere; CHR-P = clinical high-risk for psychosis; IQ = intelligence quotient; L = left; R = right

| eTable 13. Associations between either regional z-scores or observed brain morphometric measures with IQ in healthy individuals | | | | | |
|---|---|---|---|---|---|
| | | Associations with normative z-scores | | Associations with observed values | |
| **Region** | **Hemi** | **β Estimate** | **Uncorrected P values** | **β Estimate** | **Uncorrected P values** |
| **Subcortical Volume** | | | | | |
| nucleus accumbens | L | -0.01 | 0.81 | -0.02 | 0.50 |
| | R | 0.01 | 0.85 | 4.61E-03 | 0.89 |
| amygdala | L | -0.01 | 0.69 | -0.02 | 0.62 |
| | R | -0.02 | 0.52 | -0.01 | 0.66 |
| caudate | L | 0.03 | 0.47 | 0.02 | 0.51 |
| | R | 0.03 | 0.34 | 0.02 | 0.42 |
| hippocampus | L | 0.02 | 0.56 | 0.02 | 0.53 |
| | R | 0.05 | 0.20 | 0.03 | 0.26 |
| pallidum | L | -0.03 | 0.33 | -0.01 | 0.66 |
| | R | 0.03 | 0.40 | 0.04 | 0.18 |
| putamen | L | 2.87E-04 | 0.99 | 9.12E-04 | 0.98 |
| | R | -3.88E-03 | 0.91 | -2.65E-03 | 0.93 |
| thalamus | L | 0.02 | 0.61 | 0.02 | 0.54 |
| | R | 0.06 | 0.08 | 0.04 | 0.14 |
| **Cortical Thickness** | | | | | |
| banks superior temporal sulcus | L | 0.01 | 0.85 | 0.01 | 0.85 |
| | R | -0.04 | 0.29 | -0.03 | 0.29 |
| caudal anterior cingulate | L | -0.05 | 0.19 | -0.04 | 0.19 |
| | R | -0.08 | 0.02 | -0.08 | 0.02 |
| caudal middle frontal | L | 0.03 | 0.39 | 0.03 | 0.24 |
| | R | 0.05 | 0.21 | 0.04 | 0.13 |
| cuneus | L | 0.10 | 0.01 | 0.08 | 0.01 |
| | R | 0.07 | 0.04 | 0.06 | 0.06 |
| entorhinal cortex | L | 0.02 | 0.65 | 0.02 | 0.59 |
| | R | -4.52E-03 | 0.90 | -1.31E-03 | 0.97 |
| fusiform gyrus | L | -0.04 | 0.22 | -0.02 | 0.50 |
| | R | -0.04 | 0.28 | -0.02 | 0.60 |
| inferior parietal | L | -0.10 | 3.71E-03 | -0.06 | 0.01 |
| | R | -0.07 | 0.04 | -0.04 | 0.05 |
| inferior temporal | L | -0.03 | 0.34 | -0.03 | 0.34 |
| | R | -4.64E-03 | 0.90 | -0.01 | 0.76 |
| isthmus cingulate | L | -0.02 | 0.57 | -0.01 | 0.80 |
| | R | -0.04 | 0.24 | -0.02 | 0.61 |
| lateral occipital | L | -0.10 | 0.01 | -0.06 | 0.04 |
| | R | -4.14E-04 | 0.99 | 0.01 | 0.76 |
| lateral orbitofrontal | L | 0.02 | 0.66 | 0.01 | 0.81 |
| | R | -0.02 | 0.49 | -0.02 | 0.45 |
| lingual gyrus | L | 0.11 | 1.37E-03 | 0.09 | 2.50E-03 |

| eTable 13. Associations between either regional z-scores or observed brain morphometric measures with IQ in healthy individuals | | | | | |
|---|---|---|---|---|---|
| | | Associations with normative z-scores | | Associations with observed values | |
| Region | Hemi | β Estimate | Uncorrected P values | β Estimate | Uncorrected P values |
| | R | 0.08 | 0.02 | 0.07 | 0.02 |
| medial orbitofrontal | L | -0.01 | 0.82 | -0.02 | 0.58 |
| | R | 0.01 | 0.71 | 0.01 | 0.84 |
| middle temporal | L | -0.05 | 0.16 | -0.04 | 0.19 |
| | R | -0.06 | 0.07 | -0.05 | 0.04 |
| parahippocampal | L | -0.03 | 0.39 | -0.03 | 0.36 |
| | R | -0.01 | 0.82 | -0.01 | 0.72 |
| paracentral | L | 0.04 | 0.29 | 0.03 | 0.27 |
| | R | -0.01 | 0.82 | -0.02 | 0.51 |
| pars opercularis | L | 0.01 | 0.84 | 0.02 | 0.58 |
| | R | 0.06 | 0.12 | 0.05 | 0.09 |
| pars orbitalis | L | -0.02 | 0.55 | -0.02 | 0.63 |
| | R | -0.04 | 0.31 | -0.03 | 0.23 |
| pars triangularis | L | -0.01 | 0.77 | 5.32E-04 | 0.98 |
| | R | 0.02 | 0.64 | 0.01 | 0.73 |
| pericalcarine | L | 0.10 | 0.01 | 0.07 | 0.04 |
| | R | 0.08 | 0.02 | 0.06 | 0.09 |
| postcentral | L | 0.09 | 0.02 | 0.06 | 0.02 |
| | R | 0.09 | 0.01 | 0.08 | 0.01 |
| posterior cingulate | L | -2.35E-03 | 0.95 | 0.01 | 0.86 |
| | R | 0.01 | 0.73 | 0.02 | 0.53 |
| precentral | L | 0.05 | 0.14 | 0.03 | 0.22 |
| | R | 0.04 | 0.33 | 0.01 | 0.63 |
| precuneus | L | -0.01 | 0.83 | 0.01 | 0.79 |
| | R | -0.06 | 0.10 | -0.02 | 0.33 |
| rostral anterior cingulate | L | -4.14E-03 | 0.91 | -0.01 | 0.86 |
| | R | -0.05 | 0.19 | -0.03 | 0.31 |
| rostral middle frontal | L | -0.03 | 0.40 | -0.01 | 0.71 |
| | R | -0.03 | 0.47 | -0.02 | 0.49 |
| superior frontal | L | -0.07 | 0.04 | -0.04 | 0.07 |
| | R | -0.08 | 0.02 | -0.04 | 0.04 |
| superior parietal | L | 0.01 | 0.77 | 8.89E-04 | 0.97 |
| | R | -2.07E-04 | 1.00 | 4.24E-03 | 0.85 |
| superior temporal | L | 0.02 | 0.53 | 0.02 | 0.43 |
| | R | 0.04 | 0.28 | 0.03 | 0.32 |
| supramarginal gyrus | L | -0.02 | 0.55 | -0.02 | 0.50 |
| | R | 0.05 | 0.20 | 0.04 | 0.14 |
| frontal pole | L | -0.03 | 0.44 | -0.01 | 0.83 |
| | R | -0.04 | 0.30 | -0.04 | 0.26 |

| eTable 13. Associations between either regional z-scores or observed brain morphometric measures with IQ in healthy individuals | | | | | |
|---|---|---|---|---|---|
| | | Associations with normative z-scores | | Associations with observed values | |
| Region | Hemi | β Estimate | Uncorrected P values | β Estimate | Uncorrected P values |
| temporal pole | L | 0.06 | 0.08 | 0.06 | 0.09 |
| | R | 0.03 | 0.44 | 0.02 | 0.56 |
| transverse temporal | L | 0.01 | 0.85 | 0.01 | 0.76 |
| | R | 0.05 | 0.14 | 0.05 | 0.09 |
| insula | L | 0.07 | 0.05 | 0.07 | 0.02 |
| | R | 0.04 | 0.25 | 0.05 | 0.11 |
| **Surface Area** | | | | | |
| banks superior temporal sulcus | L | -0.03 | 0.40 | -0.02 | 0.45 |
| | R | -0.03 | 0.40 | -0.02 | 0.63 |
| caudal anterior cingulate | L | 0.04 | 0.27 | 0.06 | 0.09 |
| | R | -0.02 | 0.56 | -0.02 | 0.48 |
| caudal middle frontal | L | 0.09 | 0.01 | 0.07 | 0.01 |
| | R | -9.58E-04 | 0.98 | 3.75E-03 | 0.90 |
| cuneus | L | -0.02 | 0.62 | -0.01 | 0.72 |
| | R | -0.01 | 0.73 | -0.02 | 0.40 |
| entorhinal cortex | L | 0.05 | 0.13 | 0.03 | 0.30 |
| | R | 0.02 | 0.66 | 0.01 | 0.69 |
| fusiform gyrus | L | 0.01 | 0.87 | 4.26E-03 | 0.87 |
| | R | -0.02 | 0.50 | -0.02 | 0.49 |
| inferior parietal | L | -1.32E-03 | 0.97 | 2.26E-03 | 0.94 |
| | R | -0.05 | 0.13 | -0.04 | 0.15 |
| inferior temporal | L | 0.03 | 0.47 | -0.01 | 0.80 |
| | R | -0.01 | 0.81 | -0.03 | 0.25 |
| isthmus cingulate | L | -1.60E-06 | 1.00 | 1.46E-03 | 0.96 |
| | R | 0.03 | 0.34 | 0.02 | 0.48 |
| lateral occipital | L | -0.02 | 0.62 | -0.02 | 0.36 |
| | R | 0.04 | 0.26 | 0.01 | 0.68 |
| lateral orbitofrontal | L | -0.01 | 0.76 | -0.01 | 0.63 |
| | R | 0.02 | 0.65 | 0.02 | 0.53 |
| lingual gyrus | L | 0.02 | 0.54 | 0.02 | 0.49 |
| | R | -0.01 | 0.82 | 1.48E-03 | 0.96 |
| medial orbitofrontal | L | 0.02 | 0.66 | 2.92E-03 | 0.91 |
| | R | -4.35E-03 | 0.90 | -0.02 | 0.53 |
| middle temporal | L | 0.01 | 0.76 | -3.68E-03 | 0.88 |
| | R | 0.03 | 0.47 | 0.01 | 0.81 |
| parahippocampal | L | 0.08 | 0.03 | 0.08 | 0.01 |
| | R | -0.01 | 0.72 | 2.47E-04 | 0.99 |
| paracentral | L | 1.19E-03 | 0.97 | 3.49E-06 | 1.00 |
| | R | 3.84E-03 | 0.91 | 3.03E-03 | 0.92 |
| pars opercularis | L | 0.01 | 0.80 | 0.01 | 0.76 |

| eTable 13. Associations between either regional z-scores or observed brain morphometric measures with IQ in healthy individuals | | | | | |
|---|---|---|---|---|---|
| | | Associations with normative z-scores | | Associations with observed values | |
| Region | Hemi | β Estimate | Uncorrected P values | β Estimate | Uncorrected P values |
| | R | -4.79E-03 | 0.89 | -6.62E-04 | 0.98 |
| pars orbitalis | L | -0.04 | 0.22 | -0.04 | 0.11 |
| | R | 0.01 | 0.79 | 0.01 | 0.75 |
| pars triangularis | L | 0.02 | 0.51 | 0.02 | 0.46 |
| | R | 0.05 | 0.20 | 0.04 | 0.24 |
| pericalcarine | L | -3.03E-03 | 0.93 | -3.48E-03 | 0.91 |
| | R | -0.01 | 0.71 | -0.01 | 0.77 |
| postcentral | L | 0.08 | 0.02 | 0.06 | 0.01 |
| | R | 0.06 | 0.12 | 0.04 | 0.09 |
| posterior cingulate | L | 0.05 | 0.13 | 0.04 | 0.20 |
| | R | 0.02 | 0.49 | 0.01 | 0.71 |
| precentral | L | -3.51E-03 | 0.92 | -0.01 | 0.82 |
| | R | 0.02 | 0.57 | 0.01 | 0.56 |
| precuneus | L | -0.03 | 0.45 | -0.01 | 0.59 |
| | R | 0.05 | 0.18 | 0.03 | 0.18 |
| rostral anterior cingulate | L | 0.01 | 0.80 | 4.90E-03 | 0.85 |
| | R | -0.03 | 0.46 | -0.02 | 0.39 |
| rostral middle frontal | L | 0.05 | 0.19 | 0.01 | 0.59 |
| | R | 0.01 | 0.81 | -1.52E-04 | 0.99 |
| superior frontal | L | 0.01 | 0.83 | -4.96E-03 | 0.82 |
| | R | 9.11E-04 | 0.98 | -0.01 | 0.76 |
| superior parietal | L | 0.01 | 0.74 | 0.01 | 0.69 |
| | R | -6.40E-04 | 0.99 | 3.44E-03 | 0.90 |
| superior temporal | L | -0.06 | 0.11 | -0.03 | 0.15 |
| | R | 1.24E-03 | 0.97 | 0.01 | 0.80 |
| supramarginal gyrus | L | -0.04 | 0.27 | -0.03 | 0.21 |
| | R | -0.06 | 0.12 | -0.03 | 0.22 |
| frontal pole | L | 0.06 | 0.10 | 0.04 | 0.16 |
| | R | -0.02 | 0.62 | -0.02 | 0.53 |
| temporal pole | L | 0.01 | 0.81 | -7.80E-04 | 0.98 |
| | R | -0.01 | 0.70 | -0.01 | 0.77 |
| transverse temporal | L | -0.06 | 0.12 | -0.04 | 0.17 |
| | R | -2.75E-04 | 0.99 | -1.78E-03 | 0.95 |
| insula | L | -0.02 | 0.51 | -0.02 | 0.33 |
| | R | -0.02 | 0.57 | -0.02 | 0.52 |
| [a] significant at $P_{FDR}<0.05$; Hemi = hemisphere; IQ = intelligence quotient; L = left; R = right | | | | | |

| **eTable 14. Associations between average deviation scores with positive symptoms and IQ in CHR-P and healthy individuals** | | | | | | |
|---|---|---|---|---|---|---|
| | **CHR-P individuals** | | | | **Healthy individuals** | |
| **Average Deviation Scores** | **Positive Symptoms (N = 1340)** | | **IQ (N = 924)** | | **IQ (N = 797)** | |
| | **β Estimate** | **Uncorrected P-value** | **β Estimate** | **Uncorrected P-value** | **β Estimate** | **Uncorrected P-value** |
| $ADS_G$ | -0.05 | 0.08 | 0.10 | 3.25E-03[a] | 0.05 | 0.14 |
| $ADS_{SV}$ | -0.03 | 0.31 | 0.05 | 0.13 | 0.02 | 0.57 |
| $ADS_{CT}$ | 0.01 | 0.61 | 0.04 | 0.25 | 0.02 | 0.65 |
| $ADS_{SA}$ | -0.08 | 0.01[a] | 0.09 | 0.01[a] | 0.05 | 0.13 |

[a] significant at $P_{FDR} < 0.05$; $ASD_{CT}$=average deviation score-cortical thickness; $ADS_G$=Average deviation score-global; $ADS_{SA}$=average deviation score-surface area; $ADS_{SV}$=average deviation score-subcortical volume; CHR-P = clinical high-risk for psychosis; IQ = intelligence quotient

| eTable 15. Percentage of Clinical High-Risk for Psychosis (CHR-P) with infra- or supranormal normative regional z-score according to conversion status. | | | | | |
|---|---|---|---|---|---|
| | | CHR-PC (N = 157) | | CHR-PNC (N = 940) | |
| **Region** | **Hemi** | **infranormal z-scores (%)** | **supranormal z-scores (%)** | **infranormal z-scores (%)** | **supranormal z-scores (%)** |
| **Subcortical Volume** | | | | | |
| nucleus accumbens | L | 5.73 | 0.00 | 4.89 | 0.53 |
| | R | 8.28 | 0.00 | 6.17 | 0.11 |
| amygdala | L | 4.46 | 1.27 | 3.72 | 0.53 |
| | R | 5.10 | 3.82 | 3.19 | 1.70 |
| caudate | L | 8.92 | 0.64 | 5.96 | 0.53 |
| | R | 5.73 | 1.27 | 5.11 | 1.17 |
| hippocampus | L | 7.64 | 1.91 | 4.57 | 0.53 |
| | R | 5.73 | 1.91 | 3.83 | 0.53 |
| pallidum | L | 7.64 | 0.00 | 4.68 | 0.43 |
| | R | 10.19 | 0.00 | 5.21 | 1.28 |
| putamen | L | 8.28 | 0.64 | 10.00 | 0.32 |
| | R | 11.46 | 0.00 | 11.70 | 0.21 |
| thalamus | L | 9.55 | 0.64 | 8.51 | 0.96 |
| | R | 12.74 | 0.64 | 8.09 | 0.32 |
| **Cortical Thickness** | | | | | |
| banks superior temporal sulcus | L | 4.46 | 0.64 | 2.23 | 1.38 |
| | R | 4.46 | 1.91 | 3.19 | 1.81 |
| caudal anterior cingulate | L | 4.46 | 3.18 | 2.02 | 3.30 |
| | R | 3.18 | 3.82 | 3.51 | 4.57 |
| caudal middle frontal | L | 1.91 | 0.00 | 2.45 | 1.17 |
| | R | 1.91 | 1.91 | 3.30 | 1.70 |
| cuneus | L | 1.91 | 3.82 | 1.70 | 2.02 |
| | R | 3.82 | 0.64 | 1.81 | 2.98 |
| entorhinal cortex | L | 3.82 | 4.46 | 2.34 | 4.57 |
| | R | 2.55 | 1.91 | 2.45 | 4.26 |
| fusiform gyrus | L | 0.64 | 0.00 | 1.60 | 3.51 |
| | R | 2.55 | 1.91 | 1.70 | 3.51 |
| inferior parietal | L | 0.64 | 1.91 | 1.60 | 2.66 |
| | R | 3.18 | 1.91 | 1.81 | 1.81 |
| inferior temporal | L | 0.64 | 4.46 | 1.28 | 3.62 |
| | R[a] | 5.10 | 3.82 | 1.28 | 4.47 |
| isthmus cingulate | L | 5.10 | 1.27 | 5.11 | 1.91 |
| | R | 5.10 | 3.18 | 4.79 | 1.38 |
| lateral occipital | L | 0.64 | 1.91 | 0.64 | 4.57 |
| | R | 0.00 | 5.10 | 0.64 | 4.57 |
| lateral orbitofrontal | L | 1.91 | 1.91 | 1.49 | 2.02 |
| | R | 0.00 | 2.55 | 1.81 | 2.13 |
| lingual gyrus | L | 5.10 | 2.55 | 3.19 | 1.81 |

**eTable 15. Percentage of Clinical High-Risk for Psychosis (CHR-P) with infra- or supranormal normative regional z-score according to conversion status.**

| | | CHR-PC (N = 157) | | CHR-PNC (N = 940) | |
|---|---|---|---|---|---|
| **Region** | **Hemi** | **infranormal z-scores (%)** | **supranormal z-scores (%)** | **infranormal z-scores (%)** | **supranormal z-scores (%)** |
| | R | 3.82 | 1.91 | 2.23 | 1.81 |
| medial orbitofrontal | L | 1.27 | 6.37 | 2.02 | 2.87 |
| | R | 1.27 | 3.82 | 1.49 | 2.87 |
| middle temporal | L | 1.27 | 2.55 | 2.23 | 2.13 |
| | R | 1.91 | 1.27 | 2.23 | 1.60 |
| parahippocampal | L | 3.18 | 0.64 | 2.45 | 2.02 |
| | R | 2.55 | 3.18 | 1.91 | 2.98 |
| paracentral | L | 2.55 | 1.27 | 1.49 | 2.23 |
| | R | 3.82 | 1.27 | 1.60 | 1.17 |
| pars opercularis | L | 3.82 | 0.64 | 3.83 | 1.28 |
| | R | 3.82 | 0.64 | 3.94 | 0.96 |
| pars orbitalis | L | 3.18 | 1.91 | 2.66 | 1.70 |
| | R | 0.64 | 1.27 | 1.60 | 2.66 |
| pars triangularis | L | 6.37 | 2.55 | 2.98 | 2.02 |
| | R | 3.18 | 0.64 | 3.83 | 1.28 |
| pericalcarine | L | 4.46 | 4.46 | 4.15 | 3.51 |
| | R | 1.91 | 5.73 | 2.77 | 4.47 |
| postcentral | L | 1.91 | 0.64 | 0.74 | 2.55 |
| | R | 1.27 | 0.64 | 1.28 | 1.60 |
| posterior cingulate | L | 3.18 | 1.91 | 5.85 | 2.23 |
| | R | 4.46 | 0.64 | 4.26 | 1.70 |
| precentral | L | 1.91 | 1.91 | 3.09 | 0.85 |
| | R | 1.27 | 0.00 | 2.87 | 0.74 |
| precuneus | L | 1.27 | 1.27 | 1.70 | 2.34 |
| | R | 1.27 | 3.18 | 2.66 | 1.49 |
| rostral anterior cingulate | L | 1.91 | 2.55 | 3.83 | 2.87 |
| | R | 4.46 | 2.55 | 3.19 | 2.77 |
| rostral middle frontal | L | 1.27 | 2.55 | 1.60 | 1.81 |
| | R | 1.27 | 3.18 | 1.49 | 2.13 |
| superior frontal | L | 1.27 | 2.55 | 3.94 | 1.60 |
| | R | 2.55 | 0.64 | 3.19 | 1.70 |
| superior parietal | L | 0.64 | 3.82 | 0.53 | 2.55 |
| | R | 0.64 | 5.10 | 0.74 | 2.13 |
| superior temporal | L | 4.46 | 0.64 | 2.87 | 0.74 |
| | R | 4.46 | 0.00 | 3.40 | 1.28 |
| supramarginal gyrus | L | 4.46 | 1.91 | 2.13 | 0.96 |
| | R | 3.82 | 1.91 | 4.36 | 1.38 |
| frontal pole | L | 3.82 | 4.46 | 2.34 | 3.72 |
| | R | 3.18 | 6.37 | 2.34 | 3.40 |

| eTable 15. Percentage of Clinical High-Risk for Psychosis (CHR-P) with infra- or supranormal normative regional z-score according to conversion status. | | | | | |
|---|---|---|---|---|---|
| | | CHR-PC (N = 157) | | CHR-PNC (N = 940) | |
| **Region** | **Hemi** | **infranormal z-scores (%)** | **supranormal z-scores (%)** | **infranormal z-scores (%)** | **supranormal z-scores (%)** |
| temporal pole | L | 1.27 | 2.55 | 0.85 | 2.66 |
| | R | 4.46 | 1.91 | 2.45 | 1.91 |
| transverse temporal | L | 1.27 | 0.64 | 3.09 | 1.49 |
| | R | 2.55 | 1.27 | 1.38 | 1.60 |
| insula | L | 4.46 | 0.64 | 4.15 | 0.64 |
| | R | 1.27 | 0.64 | 4.36 | 0.43 |
| **Surface Area** | | | | | |
| banks superior temporal sulcus | L | 1.91 | 1.27 | 2.55 | 1.91 |
| | R[e] | 7.01 | 2.55 | 1.38 | 2.23 |
| caudal anterior cingulate | L | 0.00 | 1.27 | 0.11 | 3.62 |
| | R | 0.64 | 3.18 | 0.43 | 3.19 |
| caudal middle frontal | L | 2.55 | 2.55 | 1.49 | 2.55 |
| | R | 0.64 | 2.55 | 1.28 | 3.09 |
| cuneus | L | 3.82 | 1.91 | 2.66 | 2.77 |
| | R | 1.27 | 1.91 | 1.38 | 3.09 |
| entorhinal cortex | L | 4.46 | 6.37 | 2.45 | 5.11 |
| | R | 0.00 | 7.01 | 1.60 | 4.68 |
| fusiform gyrus | L | 4.46 | 3.82 | 2.45 | 1.91 |
| | R | 1.91 | 1.91 | 2.45 | 1.60 |
| inferior parietal | L | 1.27 | 3.18 | 3.51 | 1.70 |
| | R | 2.55 | 0.64 | 3.09 | 3.19 |
| inferior temporal | L | 1.91 | 1.27 | 2.45 | 2.45 |
| | R | 1.91 | 3.18 | 2.77 | 1.91 |
| isthmus cingulate | L | 1.27 | 1.27 | 0.74 | 3.72 |
| | R | 1.27 | 3.18 | 0.85 | 4.26 |
| lateral occipital | L | 1.91 | 3.82 | 2.13 | 3.09 |
| | R | 3.18 | 2.55 | 1.38 | 2.34 |
| lateral orbitofrontal | L | 3.82 | 1.91 | 2.55 | 2.55 |
| | R | 1.91 | 3.82 | 2.66 | 3.62 |
| lingual gyrus | L | 1.91 | 4.46 | 3.19 | 1.60 |
| | R | 3.18 | 1.27 | 2.55 | 3.30 |
| medial orbitofrontal | L | 0.64 | 2.55 | 2.13 | 2.98 |
| | R | 4.46 | 3.82 | 2.23 | 2.45 |
| middle temporal | L | 5.73 | 1.91 | 3.09 | 3.30 |
| | R | 3.18 | 1.27 | 2.66 | 2.87 |
| parahippocampal | L | 0.64 | 3.18 | 2.66 | 1.60 |
| | R | 5.73 | 2.55 | 3.30 | 2.98 |
| paracentral | L | 1.27 | 3.82 | 0.96 | 3.19 |
| | R | 0.64 | 1.27 | 0.85 | 3.51 |
| pars opercularis | L | 1.27 | 3.82 | 1.70 | 2.45 |

| eTable 15. Percentage of Clinical High-Risk for Psychosis (CHR-P) with infra- or supranormal normative regional z-score according to conversion status. | | | | | |
|---|---|---|---|---|---|
| | | CHR-PC (N = 157) | | CHR-PNC (N = 940) | |
| **Region** | **Hemi** | **infranormal z-scores (%)** | **supranormal z-scores (%)** | **infranormal z-scores (%)** | **supranormal z-scores (%)** |
| | R | 1.27 | 2.55 | 0.96 | 3.19 |
| pars orbitalis | L | 0.64 | 3.82 | 2.77 | 2.98 |
| | R | 1.27 | 1.91 | 1.38 | 2.34 |
| pars triangularis | L | 1.27 | 4.46 | 3.19 | 1.28 |
| | R | 1.91 | 1.27 | 1.81 | 2.02 |
| pericalcarine | L | 0.64 | 0.64 | 1.81 | 3.30 |
| | R | 3.18 | 1.27 | 2.23 | 2.13 |
| postcentral | L | 1.27 | 6.37 | 0.96 | 5.53 |
| | R | 2.55 | 3.18 | 1.70 | 3.40 |
| posterior cingulate | L | 1.91 | 3.82 | 1.38 | 2.66 |
| | R | 0.00 | 3.82 | 1.38 | 2.55 |
| precentral | L | 1.91 | 3.82 | 1.28 | 4.47 |
| | R | 0.64 | 6.37 | 0.43 | 3.62 |
| precuneus | L | 3.82 | 5.73 | 2.02 | 1.81 |
| | R | 2.55 | 2.55 | 2.02 | 3.19 |
| rostral anterior cingulate | L | 0.64 | 3.18 | 1.60 | 3.62 |
| | R | 0.00 | 1.27 | 0.74 | 2.55 |
| rostral middle frontal | L | 2.55 | 3.18 | 2.55 | 2.87 |
| | R | 3.18 | 3.18 | 2.23 | 3.19 |
| superior frontal | L | 5.10 | 1.27 | 3.30 | 3.72 |
| | R | 0.64 | 2.55 | 2.66 | 2.13 |
| superior parietal | L | 1.91 | 1.91 | 1.81 | 2.13 |
| | R | 1.91 | 3.18 | 2.34 | 2.45 |
| superior temporal | L | 0.64 | 3.82 | 2.77 | 3.40 |
| | R | 2.55 | 1.91 | 1.60 | 3.40 |
| supramarginal gyrus | L | 3.18 | 1.27 | 2.34 | 4.47 |
| | R | 0.00 | 1.27 | 2.13 | 2.13 |
| frontal pole | L | 1.91 | 0.64 | 1.81 | 1.28 |
| | R | 1.27 | 2.55 | 1.28 | 1.17 |
| temporal pole | L | 0.64 | 1.91 | 0.85 | 2.13 |
| | R | 0.64 | 3.82 | 1.06 | 2.66 |
| transverse temporal | L | 1.91 | 2.55 | 0.85 | 4.26 |
| | R | 0.00 | 4.46 | 0.74 | 2.45 |
| insula | L | 1.27 | 3.82 | 1.49 | 6.70 |
| | R | 1.27 | 7.64 | 0.96 | 6.91 |
| Average Deviation Scores | | | | | |
| $ADS_G$[a,c] | | 5.73 | 1.91 | 3.51 | 2.13 |
| $ADS_{SV}$[a] | | 5.10 | 3.18 | 2.87 | 2.34 |
| $ADS_{CT}$ | | 4.46 | 3.18 | 2.98 | 1.91 |

| eTable 15. Percentage of Clinical High-Risk for Psychosis (CHR-P) with infra- or supranormal normative regional z-score according to conversion status. | | | | | |
|---|---|---|---|---|---|
| | | CHR-PC (N = 157) | | CHR-PNC (N = 940) | |
| **Region** | **Hemi** | **infranormal z-scores (%)** | **supranormal z-scores (%)** | **infranormal z-scores (%)** | **supranormal z-scores (%)** |
| $ADS_{SA}$[c] | | 3.18 | 2.55 | 2.98 | 3.19 |

[a] significant two-proportion z-tests difference in infranormal z-scores between CHR-PC and healthy individuals at $P_{FDR} < 0.05$; [b] significant two-proportion z-tests difference in supranormal z-scores between CHR-PC and healthy individuals at $P_{FDR} < 0.05$; [c] significant two-proportion z-tests difference in infranormal z-scores between CHR-PNC and healthy individuals at $P_{FDR} < 0.05$; [d] significant two-proportion z-tests difference in supranormal z-scores between CHR-PNC and healthy individuals at $P_{FDR} < 0.05$; [e] significant two-proportion z-tests difference in infranormal z-scores between CHR-PC and CHR-PNC at $P_{FDR} < 0.05$; [f] significant two-proportion z-tests difference in supranormal z-scores between CHR-PC and CHR-PNC at $P_{FDR} < 0.05$; $ASD_{CT}$=average deviation score-cortical thickness; $ADS_{G}$=Average deviation score-global; $ADS_{SA}$=average deviation score-surface area; $ADS_{SV}$=average deviation score-subcortical volume; CHR-PC = clinical high-risk for psychosis individuals that converted to a psychotic disorder; CHR-PNC = clinical high-risk individuals that did not convert to a psychotic disorder; Hemi = hemisphere; L = left; R = right.

**eTable 16. Associations between regional z-scores and average deviation scores with positive symptoms and IQ based on conversion status**

| | Hemi | All CHR-P N=1340 | CHR-PC N=157 | CHR-PNC N=940 | All CHR-P N=1340 | CHR-PC N=157 | CHR-PNC N=940 |
|---|---|---|---|---|---|---|---|
| | | ***β* Estimate for Positive Symptoms** | | | ***β* Estimate for IQ** | | |
| **Subcortical volumes** | | | | | | | |
| nucleus accumbens | L | 1.53E-03 | -0.12 | 0.03 | 0.02 | 0.12 | -0.01 |
| | R | -0.02 | 0.01 | -0.03 | 0.01 | 0.02 | 0.02 |
| amygdala | L | -0.01 | -0.14 | 0.02 | 0.03 | 0.09 | 0.03 |
| | R | 1.81E-03 | -0.13 | 0.02 | 0.02 | 0.05 | 0.02 |
| caudate | L | -0.02 | -0.13 | -0.02 | 0.11[a] | 0.23 | 0.09 |
| | R | -0.03 | -0.14 | -0.05 | 0.10 | 0.20 | 0.09 |
| hippocampus | L | -0.04 | -0.12 | -0.02 | -0.01 | -0.01 | 0.01 |
| | R | -0.03 | -0.16 | -0.01 | -0.01 | 3.45E-03 | 0.02 |
| pallidum | L | -0.01 | -0.04 | -0.03 | 0.03 | 0.17 | 0.03 |
| | R | -0.02 | -0.02 | -0.04 | -0.01 | 0.07 | -0.01 |
| putamen | L | 0.01 | -0.07 | 0.01 | -0.02 | 0.10 | -0.05 |
| | R | -0.01 | -0.06 | -0.01 | 1.92E-03 | 0.09 | -0.02 |
| thalamus | L | -0.03 | -0.07 | -0.02 | 0.07 | 0.20 | 0.06 |
| | R | -0.02 | -0.11 | -0.03 | 0.06 | 0.26 | 0.03 |
| **Cortical Thickness** | | | | | | | |
| banks superior temporal sulcus | L | 0.01 | 0.04 | 0.01 | 0.01 | -0.03 | 4.25E-03 |
| | R | 0.01 | -0.11 | 0.05 | -0.03 | -0.16 | 4.34E-04 |
| caudal anterior cingulate | L | -0.02 | -0.01 | -0.01 | -0.01 | -0.25 | 0.02 |
| | R | -0.01 | 0.13 | -0.02 | 0.03 | 0.01 | 0.04 |

| eTable 16. Associations between regional z-scores and average deviation scores with positive symptoms and IQ based on conversion status | | | | | | | |
|---|---|---|---|---|---|---|---|
| | **Hemi** | **All CHR-P N=1340** | **CHR-PC N=157** | **CHR-PNC N=940** | **All CHR-P N=1340** | **CHR-PC N=157** | **CHR-PNC N=940** |
| | | ***β* Estimate for Positive Symptoms** | | | ***β* Estimate for IQ** | | |
| caudal middle frontal | L | -0.04 | -0.07 | -0.05 | 0.05 | 0.17 | 0.05 |
| | R | 0.01 | -0.09 | 0.02 | 0.05 | 0.27 | 0.01 |
| cuneus | L | -0.03 | -0.04 | -0.02 | 0.06 | 0.09 | 0.04 |
| | R | 0.01 | 1.57E-03 | 0.03 | 0.04 | 0.03 | 0.03 |
| entorhinal cortex | L | -0.02 | -0.01 | -0.03 | 0.05 | -0.04 | 0.03 |
| | R | 0.03 | -0.05 | 0.05 | 0.04 | -4.68E-03 | 0.04 |
| fusiform gyrus | L | -0.04 | -0.06 | -0.02 | 0.03 | 0.04 | 0.01 |
| | R | -0.02 | 0.03 | 4.92E-03 | -3.01E-03 | -4.77E-03 | -0.04 |
| inferior parietal | L | 0.01 | -0.02 | 0.01 | -0.06 | -0.06 | -0.09 |
| | R | 0.01 | -0.02 | 1.98E-03 | -0.07 | -0.07 | -0.06 |
| inferior temporal | L | -0.02 | -0.01 | -0.02 | -3.98E-03 | -0.09 | -0.02 |
| | R | -0.03 | -0.02 | -0.02 | 0.03 | -0.06 | 0.02 |
| isthmus cingulate | L | -5.49E-05 | 0.09 | 0.01 | -0.02 | 0.04 | -0.02 |
| | R | -0.03 | 0.06 | -0.04 | 0.03 | 0.16 | 0.01 |
| lateral occipital | L | -0.07 | -0.02 | -0.08 | -0.06 | -3.10E-04 | -0.05 |
| | R | -0.04 | 0.09 | -0.07 | -0.02 | -0.03 | -0.04 |
| lateral orbitofrontal | L | -0.03 | -0.03 | 3.23E-03 | -0.02 | -0.19 | -0.03 |
| | R | -3.76E-03 | 0.05 | 0.01 | 0.03 | -0.11 | 0.03 |
| lingual gyrus | L | 0.02 | 0.05 | 0.02 | 0.08 | 0.03 | 0.09 |
| | R | -0.01 | 0.02 | 2.73E-03 | 0.05 | 0.08 | 0.05 |

**eTable 16. Associations between regional z-scores and average deviation scores with positive symptoms and IQ based on conversion status**

| | Hemi | All CHR-P N=1340 | CHR-PC N=157 | CHR-PNC N=940 | All CHR-P N=1340 | CHR-PC N=157 | CHR-PNC N=940 |
|---|---|---|---|---|---|---|---|
| | | $\beta$ Estimate for Positive Symptoms | | | $\beta$ Estimate for IQ | | |
| medial orbitofrontal | L | 1.69E-03 | 0.10 | -3.03E-03 | -0.04 | -0.07 | -0.06 |
| | R | 0.02 | 0.18 | 3.54E-03 | -0.02 | -0.28 | -2.61E-03 |
| middle temporal | L | -3.07E-03 | -0.07 | 8.68E-04 | -0.01 | -0.17 | -0.01 |
| | R | 0.02 | 0.04 | 0.03 | -0.04 | -0.05 | -0.05 |
| parahippocampal | L | 0.01 | 0.02 | 1.47E-03 | 0.04 | 0.02 | 0.05 |
| | R | -1.02E-03 | -0.07 | 0.01 | -0.01 | 0.01 | -0.02 |
| paracentral | L | -0.01 | -0.01 | -0.02 | -0.01 | -0.02 | -0.01 |
| | R | 0.04 | 0.07 | 0.03 | -0.05 | -0.19 | -0.03 |
| pars opercularis | L | 0.02 | 0.17 | 0.01 | 3.03E-03 | -0.05 | 0.05 |
| | R | -0.01 | 0.05 | -0.01 | 0.08 | 0.03 | 0.10 |
| pars orbitalis | L | -0.01 | 0.05 | -0.01 | -0.02 | -0.03 | -0.01 |
| | R | -0.03 | -0.02 | -0.01 | -0.03 | 0.11 | -0.07 |
| pars triangularis | L | 0.02 | 0.04 | 1.07E-03 | 0.02 | 0.10 | 0.02 |
| | R | 0.04 | 0.02 | 0.06 | -0.03 | 0.03 | -0.02 |
| pericalcarine | L | 0.01 | -0.01 | 0.01 | 0.03 | 0.11 | 0.02 |
| | R | 0.01 | -0.05 | 2.41E-03 | 0.02 | 0.09 | 0.01 |
| postcentral | L | -0.02 | -0.03 | -0.02 | 0.05 | 0.11 | 0.04 |
| | R | 0.06 | 0.09 | 0.04 | 0.05 | 0.09 | 0.05 |
| posterior cingulate | L | 0.04 | 0.01 | 0.07 | -0.01 | -0.05 | 0.05 |
| | R | -0.02 | 0.03 | -0.02 | -0.02 | -0.22 | 0.01 |

**eTable 16. Associations between regional z-scores and average deviation scores with positive symptoms and IQ based on conversion status**

| | Hemi | All CHR-P N=1340 | CHR-PC N=157 | CHR-PNC N=940 | All CHR-P N=1340 | CHR-PC N=157 | CHR-PNC N=940 |
|---|---|---|---|---|---|---|---|
| | | **$\beta$ Estimate for Positive Symptoms** | | | **$\beta$ Estimate for IQ** | | |
| precentral | L | -0.01 | 3.60E-03 | -0.03 | 0.05 | -0.02 | 0.08 |
| | R | 0.01 | 0.12 | -0.02 | 0.02 | -0.02 | 0.06 |
| precuneus | L | 0.02 | 0.05 | 0.01 | 0.06 | 0.04 | 0.04 |
| | R | 0.02 | 4.61E-03 | 0.01 | -0.03 | 0.05 | -0.04 |
| rostral anterior cingulate | L | 0.02 | -0.08 | 0.04 | -0.01 | -0.11 | -0.02 |
| | R | 3.15E-03 | 0.14 | 5.68E-05 | -0.03 | -0.13 | -0.01 |
| rostral middle frontal | L | -0.03 | 0.07 | -0.05 | -0.02 | 0.20 | -0.07 |
| | R | -0.01 | 0.03 | -0.02 | -0.05 | 0.03 | -0.09 |
| superior frontal | L | -0.01 | 0.10 | -0.03 | -0.02 | -0.02 | -0.02 |
| | R | -0.01 | -0.06 | -1.61E-03 | -0.01 | 0.05 | -0.04 |
| superior parietal | L | -0.01 | -0.18 | -0.02 | -0.08 | -0.06 | -0.08 |
| | R | 0.05 | 0.05 | 0.01 | -0.03 | -4.99E-03 | -0.01 |
| superior temporal | L | -0.02 | -0.11 | 0.03 | 0.03 | 0.01 | 0.03 |
| | R | 0.03 | -0.09 | 0.08 | 0.04 | 0.10 | 0.05 |
| supramarginal gyrus | L | 0.04 | 0.01 | 0.04 | 0.03 | 0.03 | 0.05 |
| | R | 0.06 | 0.10 | 0.05 | 0.01 | 0.04 | 4.32E-03 |
| frontal pole | L | 0.01 | 0.08 | 0.01 | -0.03 | -0.16 | -0.02 |
| | R | -0.02 | 0.02 | -0.01 | 0.04 | 0.14 | -0.01 |
| temporal pole | L | 0.03 | -0.08 | 0.06 | 2.99E-03 | 0.14 | -0.03 |
| | R | 0.02 | -0.06 | 0.05 | 0.04 | 0.22 | 0.02 |

| eTable 16. Associations between regional z-scores and average deviation scores with positive symptoms and IQ based on conversion status | | | | | | | |
|---|---|---|---|---|---|---|---|
| | Hemi | All CHR-P N=1340 | CHR-PC N=157 | CHR-PNC N=940 | All CHR-P N=1340 | CHR-PC N=157 | CHR-PNC N=940 |
| | | *β* Estimate for Positive Symptoms | | | *β* Estimate for IQ | | |
| transverse temporal | L | -0.01 | 0.02 | 0.01 | -0.03 | -0.05 | -0.03 |
| | R | 0.03 | -0.04 | 0.04 | 0.01 | 0.05 | 0.04 |
| insula | L | -3.09E-04 | 0.13 | -0.01 | -0.02 | -0.13 | 0.03 |
| | R | -2.04E-03 | 0.13 | -0.01 | 0.07 | -0.14 | 0.12 |
| **Cortical Surface Area** | | | | | | | |
| banks superior temporal sulcus | L | 0.02 | -0.07 | 0.02 | -0.01 | 0.05 | 0.01 |
| | R | 0.02 | 0.10 | 0.01 | -0.07 | 9.42E-05 | -0.04 |
| caudal anterior cingulate | L | -0.01 | -0.09 | 0.01 | 0.03 | 0.10 | 0.02 |
| | R | -0.01 | 0.06 | -0.03 | 0.05 | 0.05 | 0.04 |
| caudal middle frontal | L | -0.06 | 6.91E-04 | -0.07 | 0.04 | 0.06 | 0.03 |
| | R | -3.80E-03 | 0.07 | -0.03 | 0.01 | -0.20 | 0.03 |
| cuneus | L | -2.72E-03 | -0.14 | 0.03 | 0.11[a] | 0.21 | 0.10 |
| | R | 0.01 | -0.04 | 0.02 | 0.02 | 0.12 | -0.03 |
| entorhinal cortex | L | 1.22E-03 | 0.10 | -0.04 | 0.02 | 0.19 | 0.01 |
| | R | -0.01 | 0.06 | -0.04 | 4.72E-03 | 0.14 | -0.01 |
| fusiform gyrus | L | 0.03 | 0.03 | 0.04 | 0.09 | 0.19 | 0.08 |
| | R | 0.03 | 0.02 | 0.04 | 0.05 | 0.16 | 0.06 |
| inferior parietal | L | 0.05 | 0.16 | 0.05 | -0.03 | 0.10 | -0.03 |
| | R | -0.01 | 0.05 | -0.02 | -0.01 | 0.17 | 2.00E-03 |
| inferior temporal | L | 0.02 | -0.09 | 0.03 | 0.04 | 0.06 | 0.08 |

**eTable 16. Associations between regional z-scores and average deviation scores with positive symptoms and IQ based on conversion status**

| | Hemi | All CHR-P N=1340 | CHR-PC N=157 | CHR-PNC N=940 | All CHR-P N=1340 | CHR-PC N=157 | CHR-PNC N=940 |
|---|---|---|---|---|---|---|---|
| | | ***β* Estimate for Positive Symptoms** | | | ***β* Estimate for IQ** | | |
| | R | -3.74E-03 | -0.09 | -0.01 | 0.07 | -0.02 | 0.07 |
| isthmus cingulate | L | -3.82E-03 | 0.05 | -0.01 | -0.04 | 0.03 | -0.08 |
| | R | 0.03 | 0.06 | 0.03 | -0.02 | 0.01 | -0.02 |
| lateral occipital | L | 0.01 | 0.02 | -2.75E-03 | 0.01 | -0.02 | 0.03 |
| | R | -1.66E-03 | 0.02 | -2.88E-03 | 0.06 | 0.12 | 0.05 |
| lateral orbitofrontal | L | -6.00E-04 | -0.10 | 0.01 | 4.52E-03 | 0.03 | -0.02 |
| | R | 0.01 | -0.10 | 0.02 | 0.02 | 0.09 | -0.01 |
| lingual gyrus | L | -0.01 | 0.04 | -0.01 | 0.07 | 0.11 | 0.07 |
| | R | -0.02 | -0.02 | -0.02 | 0.04 | 0.12 | 0.02 |
| medial orbitofrontal | L | -0.02 | -0.05 | -0.02 | 0.04 | -0.03 | 0.06 |
| | R | -0.04 | -0.21 | -0.01 | 0.04 | 0.28 | 0.01 |
| middle temporal | L | 0.02 | 0.01 | 0.02 | -0.01 | 0.04 | 0.02 |
| | R | 0.03 | -0.03 | 0.05 | 0.02 | 0.02 | 0.05 |
| parahippocampal | L | -0.03 | -0.17 | -0.01 | 0.02 | 0.16 | 0.03 |
| | R | -0.05 | -0.02 | -0.05 | 4.24E-03 | 0.09 | 0.02 |
| paracentral | L | 0.03 | 0.18 | 0.02 | -0.06 | -0.12 | -0.06 |
| | R | -0.05 | -0.03 | -0.03 | -0.08 | -0.19 | -0.05 |
| pars opercularis | L | -0.01 | -0.11 | -0.01 | 0.03 | 0.06 | -1.93E-03 |
| | R | -2.56E-03 | -0.04 | -0.02 | 0.04 | -0.05 | 0.04 |
| pars orbitalis | L | -0.06 | -0.04 | -0.07 | 0.05 | -0.04 | 0.06 |

| eTable 16. Associations between regional z-scores and average deviation scores with positive symptoms and IQ based on conversion status | | | | | | | |
|---|---|---|---|---|---|---|---|
| | **Hemi** | **All CHR-P N=1340** | **CHR-PC N=157** | **CHR-PNC N=940** | **All CHR-P N=1340** | **CHR-PC N=157** | **CHR-PNC N=940** |
| | | **β Estimate for Positive Symptoms** | | | **β Estimate for IQ** | | |
| | R | -0.05 | 0.04 | -0.08 | 0.06 | 0.11 | 0.03 |
| pars triangularis | L | -0.02 | 0.01 | -0.02 | 0.03 | 0.09 | 0.02 |
| | R | -0.02 | -0.05 | -0.02 | 0.01 | -0.08 | 0.01 |
| pericalcarine | L | 0.01 | -0.06 | 0.02 | 0.08 | 0.11 | 0.06 |
| | R | 3.51E-03 | -0.14 | 0.03 | 0.02 | 0.14 | -0.04 |
| postcentral | L | -0.03 | 0.02 | -0.02 | 0.01 | -0.01 | 3.05E-03 |
| | R | -0.03 | 0.03 | -0.02 | -0.02 | 0.07 | -0.04 |
| posterior cingulate | L | -2.80E-03 | 0.06 | -0.03 | -0.01 | -0.06 | -0.03 |
| | R | 0.01 | 0.12 | -0.02 | 0.01 | 0.10 | 0.01 |
| precentral | L | -0.04 | -0.12 | -0.03 | 0.02 | 0.08 | -0.02 |
| | R | -0.01 | -0.13 | -0.01 | -1.18E-03 | -0.09 | 0.02 |
| precuneus | L | 0.01 | -0.04 | -0.01 | -0.09 | -0.02 | -0.09 |
| | R | -0.02 | -0.07 | -0.03 | 0.02 | -0.04 | 0.05 |
| rostral anterior cingulate | L | -0.04 | -0.08 | 2.32E-03 | 0.09 | 0.28 | 0.07 |
| | R | -0.04 | -0.09 | -0.04 | 8.64E-04 | 0.03 | -0.02 |
| rostral middle frontal | L | 0.02 | 0.09 | 0.01 | 0.04 | -0.11 | 0.07 |
| | R | 0.02 | 0.04 | 0.02 | 0.03 | 0.12 | 0.01 |
| superior frontal | L | 0.01 | 0.03 | 5.56E-05 | -3.30E-04 | -0.05 | 2.16E-03 |
| | R | -0.03 | -0.01 | -0.03 | -0.01 | -0.18 | 0.02 |
| superior parietal | L | -0.02 | 0.01 | -0.02 | -0.02 | -0.09 | -0.04 |

| **eTable 16. Associations between regional z-scores and average deviation scores with positive symptoms and IQ based on conversion status** | | | | | | | |
|---|---|---|---|---|---|---|---|
| | **Hemi** | **All CHR-P N=1340** | **CHR-PC N=157** | **CHR-PNC N=940** | **All CHR-P N=1340** | **CHR-PC N=157** | **CHR-PNC N=940** |
| | | **$\beta$ Estimate for Positive Symptoms** | | | **$\beta$ Estimate for IQ** | | |
| | R | -0.01 | -0.05 | -4.86E-03 | -0.04 | -0.04 | -0.05 |
| superior temporal | L | -0.03 | -0.04 | -0.03 | -0.03 | -0.05 | -0.06 |
| | R | 0.03 | 0.10 | 0.04 | -0.04 | -0.14 | -0.03 |
| supramarginal gyrus | L | -0.03 | 0.11 | -0.08 | -0.03 | -0.13 | -0.02 |
| | R | -0.03 | 0.13 | -0.05 | -0.03 | -0.12 | -0.03 |
| frontal pole | L | -0.03 | 0.02 | -0.07 | -0.01 | -0.13 | 0.03 |
| | R | 0.01 | -0.01 | 2.96E-03 | -0.01 | -0.16 | -0.02 |
| temporal pole | L | 0.02 | 0.03 | 0.01 | -0.01 | 0.13 | -0.04 |
| | R | 4.22E-03 | -0.08 | -7.68E-04 | -0.06 | -0.15 | -0.08 |
| transverse temporal | L | -0.03 | -0.05 | -0.03 | -0.03 | -0.10 | -0.05 |
| | R | -0.01 | 0.04 | -4.31E-03 | -0.04 | -0.26 | -0.05 |
| insula | L | -0.02 | -0.08 | -0.01 | -0.01 | 0.10 | -0.04 |
| | R | -0.05 | -0.16 | -0.03 | -0.04 | 0.07 | -0.08 |
| **Average Deviation Scores** | | | | | | | |
| $ADS_G$ | | -0.05 | -0.07 | -0.04 | 0.10[a] | 0.21[a] | 0.07 |
| $ADS_{SV}$ | | -0.03 | -0.15 | -0.02 | 0.05 | 0.20[a] | 0.04 |
| $ADS_{CT}$ | | 0.01 | 0.12 | 0.03 | 0.04 | -0.03 | 0.04 |
| $ADS_{SA}$ | | -0.08[a] | -0.12 | -0.09[a] | 0.08[a] | 0.26[a] | 0.04 |

[a] significant associations at $P_{FDR}<0.05$; $ADS_G$=Average deviation score-global; $ASD_{CT}$=average deviation score-cortical thickness; $ADS_{SA}$=average deviation score-surface area; $ADS_{SV}$=average deviation score-subcortical volume; CHR-P = clinical high-risk for psychosis; CHR-PC = clinical high-risk for psychosis converters; CHR-PNC = clinical high-risk for psychosis non-converters; L=left; R=right

| **eTable 17. Effect size (Cohen's d) of group differences** | | | | |
|---|---|---|---|---|
| **Region** | **Hemi** | **Cohen's *d*** <br><br> **All CHR-P vs Healthy individuals** | **Cohen's *d*** <br><br> **CHR-PC vs Healthy individuals** | **Cohen's *d*** <br><br> **medicated CHR-P vs unmedicated CHR-P** |
| **Subcortical Volumes** | | | | |
| nucleus accumbens | L | -3.42E-03 | -0.11 | -0.20 |
| | R | -0.01 | -0.06 | -0.14 |
| amygdala | L | -0.03 | -0.08 | -0.17 |
| | R | -0.07 | -0.10 | -0.15 |
| caudate | L | 0.01 | -0.02 | -0.06 |
| | R | -0.02 | -0.04 | -0.07 |
| hippocampus | L | -0.11 | -0.21 | -0.14 |
| | R | -0.15 | -0.26 | -0.14 |
| pallidum | L | -0.05 | -0.13 | -0.02 |
| | R | -0.04 | -0.20 | -0.01 |
| putamen | L | -0.02 | -0.03 | -0.06 |
| | R | -0.05 | -0.08 | -0.01 |
| thalamus | L | -0.03 | -0.20 | -0.05 |
| | R | -0.09 | -0.22 | -0.14 |
| **Cortical Thickness** | | | | |
| banks superior temporal sulcus | L | -0.04 | -0.09 | 0.01 |
| | R | -0.03 | -0.12 | -0.05 |
| caudal anterior cingulate | L | -0.03 | 0.02 | -0.03 |
| | R | -0.04 | -0.04 | 0.08 |
| caudal middle frontal | L | 0.03 | -0.06 | -0.04 |
| | R | -0.04 | 0.02 | -0.02 |
| cuneus | L | 0.02 | -0.03 | -0.13 |
| | R | -0.01 | -0.10 | -0.08 |
| entorhinal cortex | L | 0.02 | -0.04 | 0.01 |
| | R | 0.01 | -0.10 | 0.11 |
| fusiform gyrus | L | -0.06 | -0.15 | 0.06 |
| | R | -0.02 | -0.14 | 0.01 |
| inferior parietal | L | -0.05 | -0.06 | 0.03 |
| | R | 0.04 | 0.12 | -0.04 |

| eTable 17. Effect size (Cohen's d) of group differences | | | | |
|---|---|---|---|---|
| **Region** | **Hemi** | **Cohen's *d*** **All CHR-P vs Healthy individuals** | **Cohen's *d*** **CHR-PC vs Healthy individuals** | **Cohen's *d*** **medicated CHR-P vs unmedicated CHR-P** |
| inferior temporal | L | 0.01 | 0.03 | 0.03 |
| | R | -0.01 | -4.70E-03 | -0.06 |
| isthmus cingulate | L | -1.84E-03 | -0.01 | -0.05 |
| | R | -3.40E-03 | 0.03 | -1.90E-03 |
| lateral occipital | L | -0.01 | -0.11 | -1.43E-03 |
| | R | -0.04 | -0.10 | 0.02 |
| lateral orbitofrontal | L | -0.03 | -0.03 | -0.06 |
| | R | 4.36E-03 | 0.13 | 0.01 |
| lingual gyrus | L | -0.05 | -0.14 | -0.02 |
| | R | 4.83E-03 | -0.07 | -0.05 |
| medial orbitofrontal | L | -0.05 | 0.08 | 0.04 |
| | R | 3.58E-03 | 0.07 | 0.06 |
| middle temporal | L | 0.03 | 0.06 | -0.01 |
| | R | -0.01 | -0.07 | -0.02 |
| parahippocampal | L | -0.07 | -0.10 | -0.07 |
| | R | -0.03 | -0.05 | -0.10 |
| paracentral | L | -0.05 | -0.21 | 0.03 |
| | R | -0.01 | -0.17 | -2.78E-03 |
| pars opercularis | L | 0.06 | 0.19 | -0.05 |
| | R | 0.02 | 1.11E-03 | 0.03 |
| pars orbitalis | L | 0.02 | 0.04 | -0.03 |
| | R | 0.10 | 0.04 | -0.01 |
| pars triangularis | L | 0.10 | 0.09 | -0.03 |
| | R | 0.05 | 0.03 | 0.05 |
| pericalcarine | L | 0.01 | -0.04 | -0.03 |
| | R | 0.02 | 0.01 | 0.03 |
| postcentral | L | 0.03 | -0.01 | -0.13 |
| | R | -0.03 | -0.06 | -0.20 |
| posterior cingulate | L | -0.04 | 0.02 | -0.10 |
| | R | -0.07 | -0.10 | 0.06 |

| eTable 17. Effect size (Cohen's d) of group differences | | | | |
|---|---|---|---|---|
| **Region** | **Hemi** | **Cohen's *d*** <br> **All CHR-P vs Healthy individuals** | **Cohen's *d*** <br> **CHR-PC vs Healthy individuals** | **Cohen's *d*** <br> **medicated CHR-P vs unmedicated CHR-P** |
| precentral | L | 0.03 | 0.10 | 0.02 |
| | R | -3.97E-03 | -0.05 | 0.03 |
| precuneus | L | -8.14E-04 | 0.08 | 0.06 |
| | R | -0.04 | 0.04 | -0.13 |
| rostral anterior cingulate | L | -0.02 | -0.04 | 0.02 |
| | R | -1.61E-03 | -0.05 | 0.03 |
| rostral middle frontal | L | -0.01 | 0.05 | -0.24 |
| | R | 1.62E-03 | 0.03 | -0.09 |
| superior frontal | L | 0.07 | 0.09 | 0.04 |
| | R | 0.07 | 0.01 | 0.01 |
| superior parietal | L | 0.03 | 0.09 | 0.02 |
| | R | 0.02 | 0.07 | -0.04 |
| superior temporal | L | 1.95E-03 | -0.15 | -0.05 |
| | R | -0.01 | -0.22 | -0.11 |
| supramarginal gyrus | L | 0.04 | 0.07 | 0.04 |
| | R | 0.01 | 0.12 | 0.05 |
| frontal pole | L | 0.04 | -0.01 | 0.06 |
| | R | -0.01 | 0.03 | 0.01 |
| temporal pole | L | 1.44E-03 | -0.04 | -0.09 |
| | R | -0.02 | -0.05 | -0.01 |
| transverse temporal | L | 0.03 | 0.03 | 0.02 |
| | R | -0.02 | -0.10 | 0.12 |
| insula | L | -0.10 | -0.03 | -0.01 |
| | R | -0.07 | -0.04 | -0.05 |
| **Cortical Surface Area** | | | | |
| banks superior temporal sulcus | L | -0.07 | -0.05 | 0.17 |
| | R | -0.04 | -0.21 | 0.04 |
| caudal anterior cingulate | L | -0.01 | -0.06 | -0.06 |
| | R | -0.03 | -0.05 | -0.01 |
| caudal middle frontal | L | 1.92E-03 | 0.06 | 0.05 |

| eTable 17. Effect size (Cohen's d) of group differences | | | | |
|---|---|---|---|---|
| **Region** | **Hemi** | **Cohen's *d*** <br> **All CHR-P vs Healthy individuals** | **Cohen's *d*** <br> **CHR-PC vs Healthy individuals** | **Cohen's *d*** <br> **medicated CHR-P vs unmedicated CHR-P** |
| | R | 0.01 | 0.07 | 2.71E-03 |
| cuneus | L | -0.06 | -0.05 | 0.06 |
| | R | -0.04 | -0.05 | -0.08 |
| entorhinal cortex | L | -0.08 | -0.13 | 0.03 |
| | R | -0.03 | 0.08 | 0.21 |
| fusiform gyrus | L | -0.03 | -0.03 | 0.01 |
| | R | -0.05 | -0.01 | -0.01 |
| inferior parietal | L | -0.06 | 0.07 | -0.07 |
| | R | -0.06 | -0.05 | 0.03 |
| inferior temporal | L | 3.78E-03 | 0.04 | 0.04 |
| | R | -0.04 | 0.02 | 0.04 |
| isthmus cingulate | L | 0.03 | -0.04 | -0.09 |
| | R | 0.08 | 0.08 | 0.12 |
| lateral occipital | L | 0.01 | -0.03 | -1.50E-03 |
| | R | 0.02 | -0.04 | -0.02 |
| lateral orbitofrontal | L | 0.02 | -0.04 | 0.04 |
| | R | -0.02 | -0.02 | -0.05 |
| lingual gyrus | L | -0.03 | -0.08 | -0.02 |
| | R | 0.02 | -0.07 | -0.06 |
| medial orbitofrontal | L | -0.06 | -0.01 | 0.01 |
| | R | -0.10 | -0.09 | 0.05 |
| middle temporal | L | 0.06 | 0.07 | 0.03 |
| | R | -0.01 | -0.19 | -0.06 |
| parahippocampal | L | -0.02 | 0.09 | -0.05 |
| | R | 0.04 | 0.03 | -0.04 |
| paracentral | L | 0.04 | 0.18 | -0.01 |
| | R | 0.07 | 0.02 | -0.04 |
| pars opercularis | L | -0.06 | -0.02 | -0.02 |
| | R | 0.02 | 0.03 | 0.05 |
| pars orbitalis | L | 0.06 | 0.12 | 0.05 |

| eTable 17. Effect size (Cohen's d) of group differences | | | | |
|---|---|---|---|---|
| **Region** | **Hemi** | **Cohen's *d*** All CHR-P vs Healthy individuals | **Cohen's *d*** CHR-PC vs Healthy individuals | **Cohen's *d*** medicated CHR-P vs unmedicated CHR-P |
| | R | 0.07 | 0.04 | 0.05 |
| pars triangularis | L | -0.04 | 2.41E-03 | 0.06 |
| | R | 0.04 | 0.16 | -0.01 |
| pericalcarine | L | -0.04 | -0.10 | -0.04 |
| | R | -0.06 | -0.15 | -0.01 |
| postcentral | L | 0.06 | 0.12 | 0.03 |
| | R | -0.02 | -0.02 | -0.01 |
| posterior cingulate | L | -0.02 | 0.06 | 0.08 |
| | R | -0.03 | 0.14 | -0.01 |
| precentral | L | 0.09 | 0.04 | -0.02 |
| | R | 0.07 | 0.13 | -0.04 |
| precuneus | L | 0.03 | 0.06 | 0.01 |
| | R | 0.01 | 0.04 | 0.08 |
| rostral anterior cingulate | L | -0.09 | -0.11 | -0.07 |
| | R | -0.04 | 0.04 | 0.03 |
| rostral middle frontal | L | 1.31E-03 | 0.17 | 0.03 |
| | R | 0.04 | 0.13 | 0.02 |
| superior frontal | L | 0.01 | 0.01 | -0.06 |
| | R | -0.08 | -0.05 | 0.02 |
| superior parietal | L | 0.05 | 0.07 | -0.01 |
| | R | 0.06 | 0.12 | -0.01 |
| superior temporal | L | -2.16E-05 | -0.08 | -0.05 |
| | R | -0.02 | -0.17 | 0.04 |
| supramarginal gyrus | L | -0.05 | -0.11 | 0.02 |
| | R | -0.03 | -0.02 | 0.08 |
| frontal pole | L | 0.04 | 0.11 | -0.05 |
| | R | 0.04 | 0.11 | -0.10 |
| temporal pole | L | -0.01 | -0.04 | -0.02 |
| | R | 0.02 | 0.05 | -0.03 |
| transverse temporal | L | -0.03 | -0.02 | 0.02 |

| eTable 17. Effect size (Cohen's d) of group differences | | | | |
|---|---|---|---|---|
| Region | Hemi | Cohen's *d*<br>All CHR-P vs Healthy individuals | Cohen's *d*<br>CHR-PC vs Healthy individuals | Cohen's *d*<br>medicated CHR-P vs unmedicated CHR-P |
| | R | 0.01 | 0.10 | -0.10 |
| insula | L | -0.01 | -0.12 | -0.05 |
| | R | -0.03 | -0.12 | 0.04 |
| **Average Deviation Scores** | | | | |
| $ADS_G$ | | -0.09 | -0.22 | -0.15 |
| $ADS_{SV}$ | | -0.08 | -0.22 | -0.16 |
| $ADS_{CT}$ | | -0.02 | -0.15 | -0.12 |
| $ADS_{SA}$ | | -0.05 | 0.02 | 0.04 |

$ASD_{CT}$=average deviation score-cortical thickness; $ADS_G$=Average deviation score-global; $ADS_{SA}$=average deviation score-surface area; $ADS_{SV}$=average deviation score-subcortical volume; CHR-P = clinical high-risk for psychosis; CHR-PC = clinical high-risk for psychosis converters; L=left; R=right

| eTable 18. Association between positive or negative average deviation scores with positive symptoms and IQ | | | | | | |
|---|---|---|---|---|---|---|
| | CHR-P Individuals | | | | Healthy Individuals | |
| | Positive Symptoms | | IQ | | IQ | |
| Average Deviation Score | β Estimate | Uncorrected P value | β Estimate | Uncorrected P value | β Estimate | Uncorrected P value |
| **P-ADS$_G$** | 0.02 | 0.54 | -2.41E-03 | 0.94 | -0.04 | 0.26 |
| **P-ADS$_{SV}$** | -0.02 | 0.61 | 0.06 | 0.07 | -0.01 | 0.80 |
| **P-ADS$_{CT}$** | 0.02 | 0.51 | -0.09 | 0.01[a] | -0.03 | 0.34 |
| **P-ADS$_{SA}$** | 0.01 | 0.67 | 0.07 | 0.04 | -0.01 | 0.72 |
| **N-ADS$_G$** | -0.02 | 0.38 | 0.06 | 0.05 | 0.03 | 0.46 |
| **N-ADS$_{SV}$** | -0.04 | 0.14 | 0.04 | 0.27 | 0.05 | 0.18 |
| **N-ADS$_{CT}$** | -0.01 | 0.71 | 0.10 | 1.37E-03[a] | 0.01 | 0.72 |
| **N-ADS$_{SA}$** | -0.01 | 0.84 | -0.03 | 0.37 | -0.01 | 0.82 |

[a] significant at $P_{FDR}<0.05$; CHR-P = clinical high-risk for psychosis; IQ = intelligence quotient; N-$ASD_{CT}$ = Negative Average Deviation Score-cortical thickness; N-$ASD_G$ = Negative Average Deviation Score-global; N-$ASD_{SA}$ = Negative Average Deviation Score-cortical surface area; N-$ASD_{SV}$ = Negative Average Deviation Score-subcortical volumes; P-$ASD_{CT}$ = Positive Average Deviation Score-cortical thickness; P-$ASD_G$ = Positive Average Deviation Score-global; P-$ASD_{SA}$ = Positive Average Deviation Score-cortical surface area; P-$ASD_{SV}$ = Positive Average Deviation Score-subcortical volumes.

**eFigure 1.** Flow diagram for study sample selection of Clinical High-Risk for Psychosis (CHR-P) individuals and healthy individuals used as controls (HI).

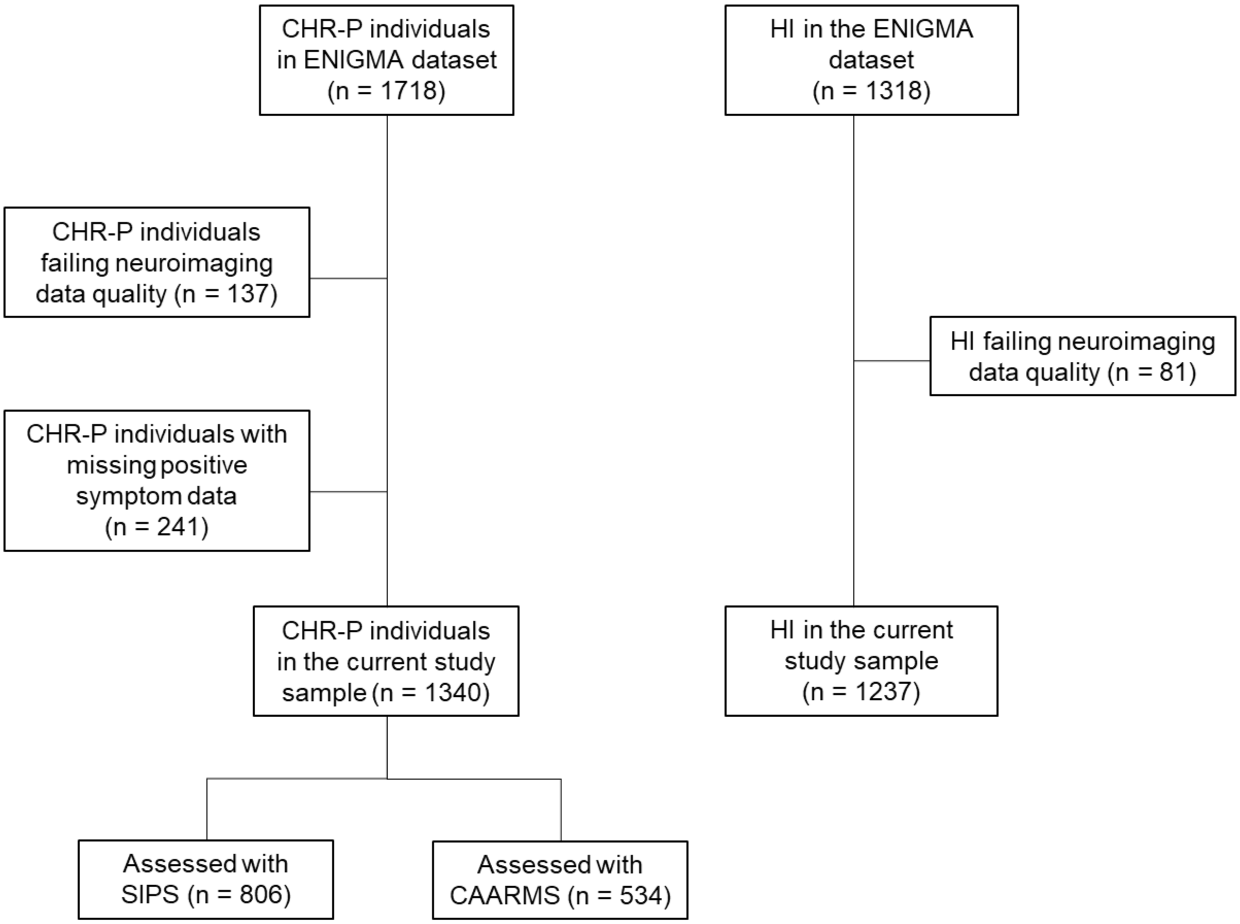

**eFigure 2.** Normative modeling of regional brain morphometric measures.

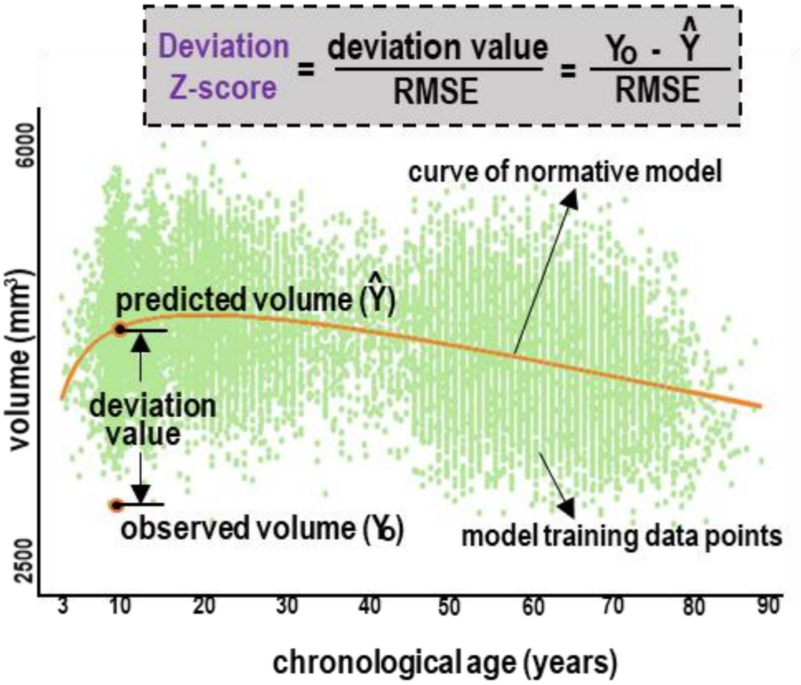


The figure illustrates the computation of the deviation z-score using the right hippocampus as an example. The green dots represent the observed hippocampal volumes and the orange curve is the FPR estimated mean.

**eFigure 3.** Distribution of normative regional z-scores for cortical thickness based on the Schaefer 400 parcellation for the A) left and B) right hemisphere.

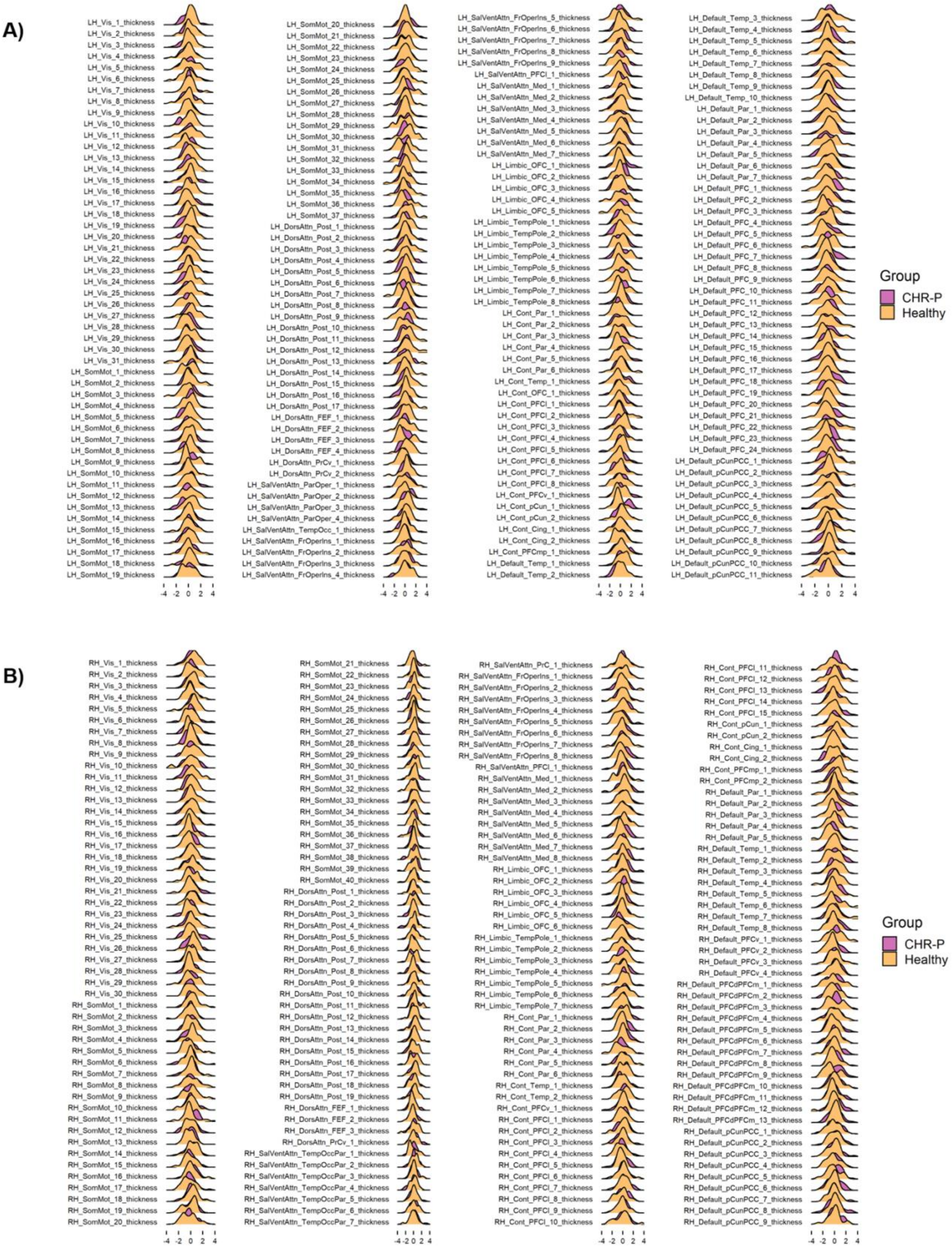

**eFigure 4.** Distribution of normative regional z-scores for cortical surface area based on the Schaefer 400 parcellation for the A) left and B) right hemisphere.

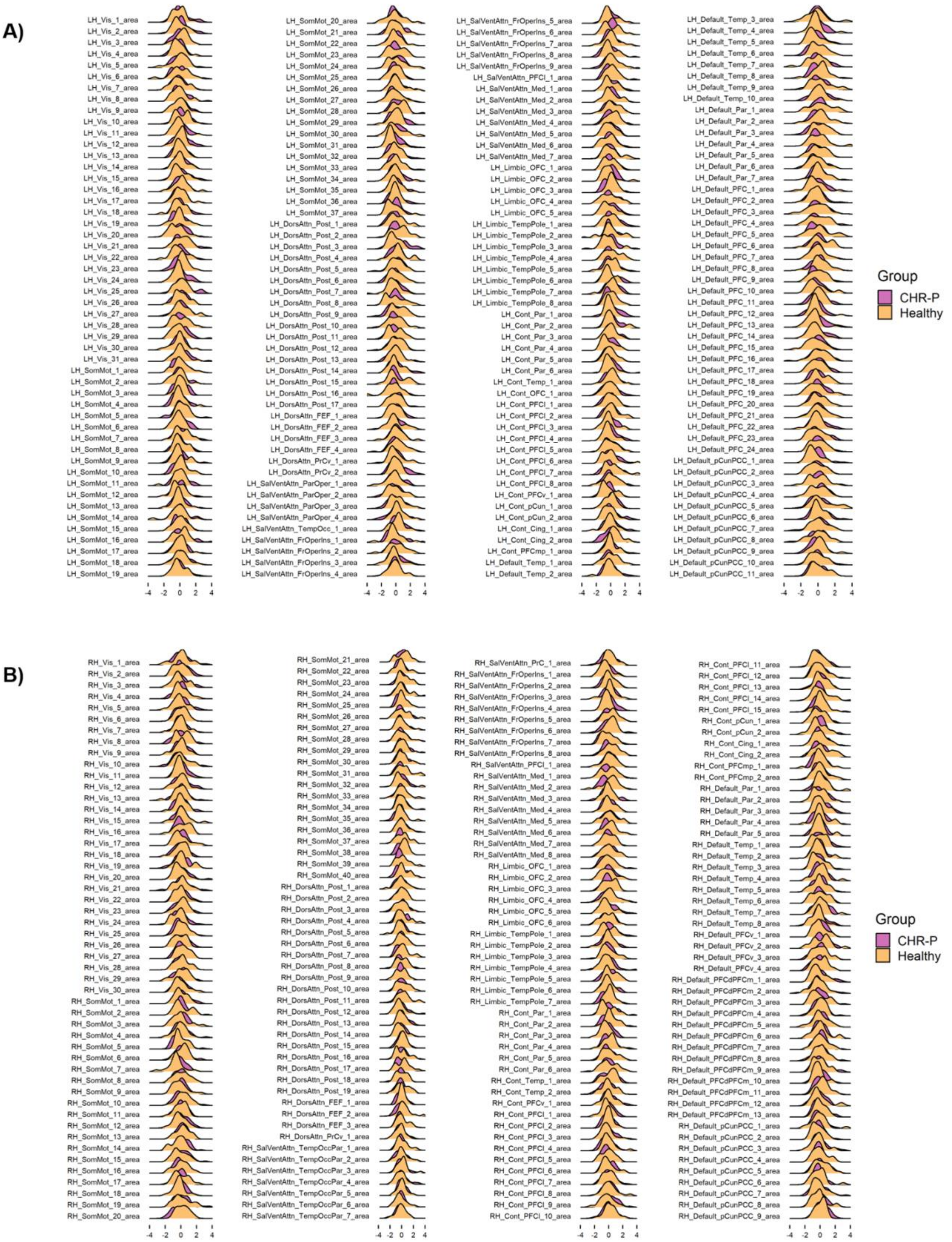

**eFigure 5. Associations between regional and average deviation scores and clinical measures in CHR-P.**

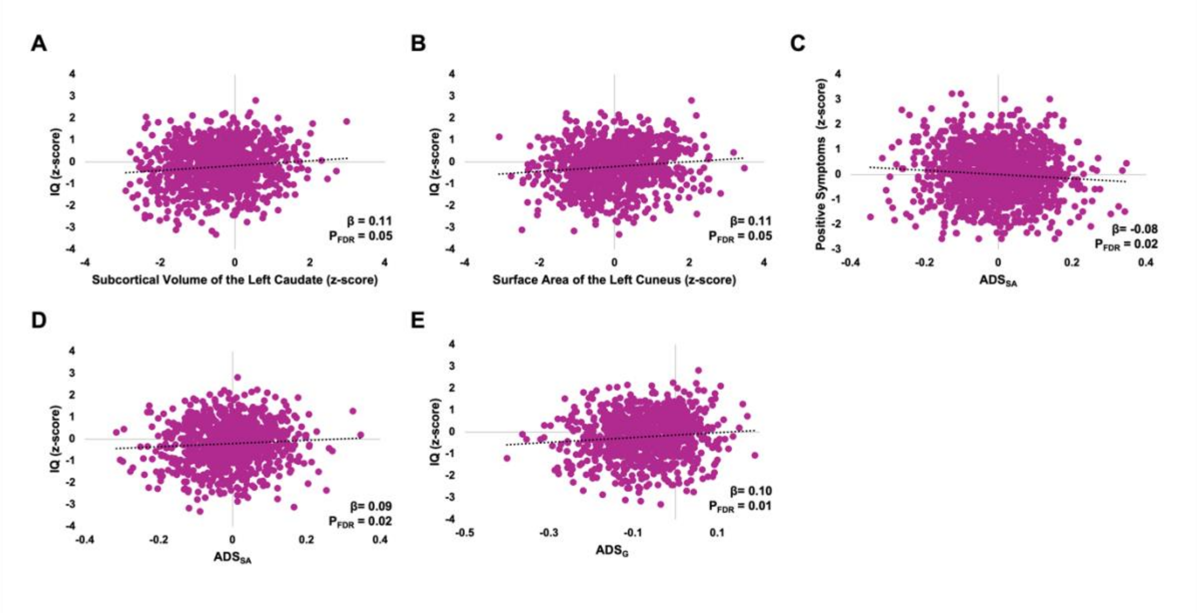


A-B, The associations between IQ and regional normative z-scores of the A) left caudate subcortical volume and B) the surface area of the left cuneus across individuals at clinical high-risk for psychosis (CHR-P). C, The association between C) positive symptoms and cortical surface area average deviation score ($ADS_{SA}$) in CHR-P. D-E, The association between IQ and $ADS_{SA}$ and global average deviation score ($ADS_G$) across CHR-P.

**eFigure 6**. Associations between average deviation scores with the positive symptoms and IQ based on medication exposure and removing one site at a time.

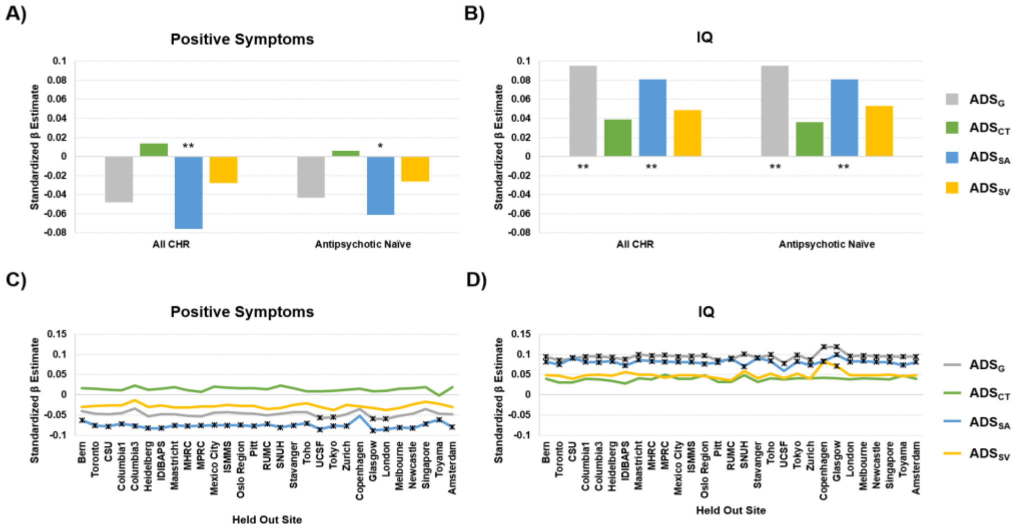

Standardized beta (β) estimates are presented in the y-axis; The effect of antipsychotic medication exposure is shown on the x-axis for A and B, the site left-out is shown on the x-axis for C and D; $ADS_G$=Average deviation score-global; $ADS_{CT}$=Average deviation score-cortical thickness; $ADS_{SA}$=Average deviation score-cortical surface area; $ADS_{SV}$= Average deviation score-subcortical volume; IQ = intelligence quotient, *significant at uncorrected P < 0.05; ** significant at $P_{FDR}$ < 0.05.

**eReferences.**
